# Aging Fly Cell Atlas Identifies Exhaustive Aging Features at Cellular Resolution

**DOI:** 10.1101/2022.12.06.519355

**Authors:** Tzu-Chiao Lu, Maria Brbić, Ye-Jin Park, Tyler Jackson, Jiaye Chen, Sai Saroja Kolluru, Yanyan Qi, Nadja Sandra Katheder, Xiaoyu Tracy Cai, Seungjae Lee, Yen- Chung Chen, Niccole Auld, Chung-Yi Liang, Sophia H. Ding, Doug Welsch, Samuel D’Souza, Angela Oliveira Pisco, Robert C. Jones, Jure Leskovec, Eric C. Lai, Hugo J. Bellen, Liqun Luo, Heinrich Jasper, Stephen R. Quake, Hongjie Li

## Abstract

Aging is characterized by a decline in tissue function, but the underlying changes at cellular resolution across the organism remain unclear. Here, we present the Aging Fly Cell Atlas, a single-nucleus transcriptomic map of the whole aging *Drosophila*. We characterize 163 distinct cell types and perform an in-depth analysis of changes in tissue cell composition, gene expression, and cell identities. We further develop aging clock models to predict the fly age and show that ribosomal gene expression is a conserved predictive factor for age. Combining all aging features, we find unique cell type-specific aging patterns. This atlas provides a valuable resource for studying fundamental principles of aging in complex organisms.

## Introduction

Aging is characterized by the progressive decline in tissue function across the entire body. It is a major risk factor for a wide range of diseases, including cardiovascular diseases, cancers, and neurodegenerative diseases (*1, 2*). Aging phenotypes have been observed and described for centuries and a number of different aging hypotheses have been proposed (*3*). However, critical questions remain largely unaddressed in complex organisms: How does aging impact cell composition and the maintenance of specific cell types? Do different cell types age at the same rate? Can we use one cell type’s transcriptome to predict age? What genes and signaling pathways drive aging in different cell types?

The fruit fly *Drosophila melanogaster* has been at the basis of many key discoveries in genetics, neurobiology, development, and aging. About 75% of human disease-associated genes have functional homologs in the fly (*4, 5*). Many of the age-related functional changes in humans are also observed in flies, including a decline in motor activity, learning and memory, cardiac function, and fertility (*6*). Hence, a proper description of the molecular and genetic basis of the age-related decline in flies should provide an important resource for aging studies not only in flies but also in other organisms.

The recent development of single-cell RNA sequencing (scRNA-seq) technologies and the establishment of the Fly Cell Atlas (FCA) (*7*), a single-nucleus transcriptomic atlas of *Drosophila* at the age of 5 days (5d hereafter), have made it possible to investigate aging phenotypes across the whole organism at single-cell resolution. Here, we present the Aging Fly Cell Atlas (AFCA), a single-nucleus transcriptomic map describing age-related changes in most tissues, including sex differences. We performed an in-depth analysis of age-related gene expression and cell composition changes across the entire fly, as well as cell type-specific and common pathways that correlate with aging. Interestingly, we observed a significant increase of fat body nuclei and a dramatic decrease in muscle nuclei with age. Furthermore, we developed aging clock models that predict the animal’s biological age from the single-nucleus transcriptomic data. In addition, we found aging variances in expressed gene number, as well as cell-type identity. Our analysis revealed that different cell types are uniquely impacted by different aging features. The aging atlas provides a valuable resource for the *Drosophila* and aging community as a reference to study aging and age-related diseases, and to evaluate the success of anti-aging regimens. We developed a website portal for data visualization and custom analyses, and made data available at the CELLxGENE portal (**fig. S1 and S2**). All resources can be accessed at https://hongjielilab.org/afca.

## Results

### Single-Nucleus Transcriptomes of the Entire Fly from Different Ages

To generate the AFCA, we applied the same single-nucleus RNA sequencing (snRNA-seq) pipeline used for the Fly Cell Atlas (FCA, 5d adults) (*7*) and profiled the whole head and body from three additional ages (30d, 50d and 70d). These time points were chosen to cover the lifespan trajectory of a fly (**Fig. 1A**), including 70d old flies, the estimated equivalent of 80-90 year old humans. Male and female flies were sequenced separately, allowing the investigation of sexual dimorphism during aging (**Fig. 1B**). To achieve the most reliable analyses of aging features, we performed preprocessing of the new aging data similarly to the young FCA data (**fig. S1A**). Consistent with a previous scRNA-seq study of the aging fly brain (*8*), we found that young and old cells have a similar distribution in the tSNE space, suggesting the whole organism largely maintains its cell types during aging (**Fig. 1C**). Overall, we obtained more than 868,000 nuclei covering all 17 broad cell-type classes (**Fig. 1, D and E**). The detected numbers of expressed genes and unique molecular identifiers (UMIs) were largely consistent across different ages (**fig. S3**). The most abundant cell classes were neurons, epithelial cells, muscle, and fat cells. Next, we annotated those broad cell classes into detailed cell types.

**Figure 1.**
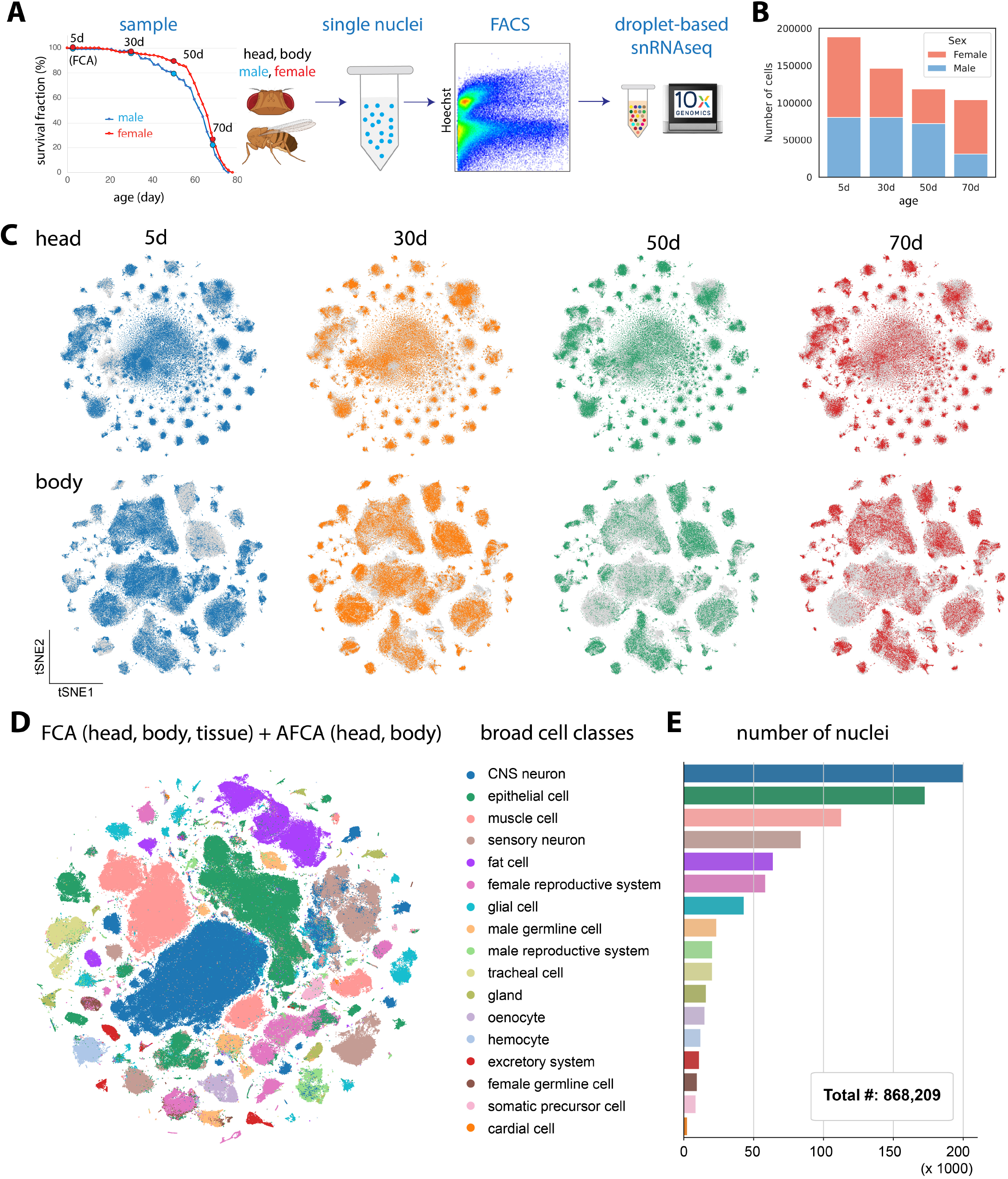
Overview of AFCA. **A)** Flowchart of snRNA-seq experiment. Flies are collected at 30, 50, and 70 days. The heads and bodies of males and females are processed separately. 5d samples are from the FCA. **B)** Number of nuclei collected from different ages and sexes. **C)** tSNE visualizations of the head and body samples from different ages. **D)** tSNE visualizations showing broad cell classes of datasets integrated across different time points. **E)** Number of nuclei for each broad cell class shown in 1D.

### AFCA Cell Type Annotation and Resource for Studying Cell Type-Specific Aging

Since cell type-specific aging analysis largely depends on accurate cell-type annotation, we took multiple approaches to ensure that our new AFCA data are annotated with high confidence. We first co-clustered our aging data with the annotated FCA data, from either the head/body or individual tissues. Then we transferred AFCA annotations using both a cluster-centered method and a supervised machine-learning-based method (**fig. S4 and S5**). Overall, we found that these two approaches agree well with about 80% overlap (**Fig. 2A**). The discrepancies of the non-overlapping annotations were mostly due to uncharacterized cell types in the FCA data or cell types with aging differences. Next, we manually validated each annotation using cell type-specific markers. Marker validation confirmed the accuracy of our automatic annotation procedure, with a few exceptions such as indirect flight muscles due to age-related loss of specific markers (**fig. S6, A and B**) and gut cell types due to the high similarity between intestinal stem cells and renal stem cells (**fig. S6C**). These cell types were then manually added and corrected (**fig. S5 and S7**). Overall, we characterized 163 distinct cell types, including 91 cell types from the aging head and 72 cell types from the aging body (**Fig. 2B and fig. S8 and S9**).

**Figure 2.**
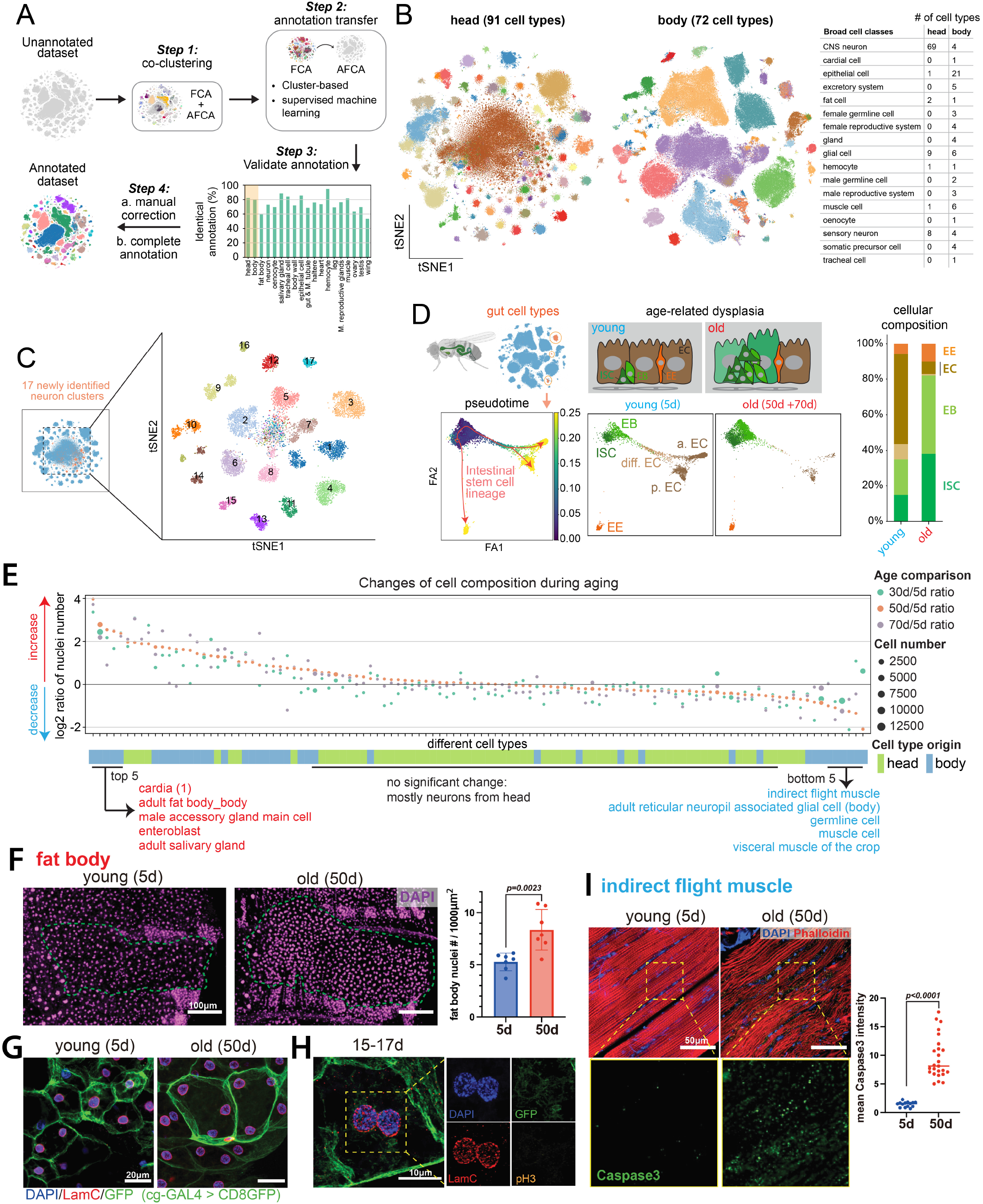
AFCA resource and changes of cell composition during aging. **A)** Flowchart of transferring annotations from FCA to AFCA. **B)** Cell types annotated in the AFCA head and body shown on tSNE. The number of annotated cell types corresponding to the broad cell classes is shown in the table. **C)** Identification of 17 new neuronal clusters after combining AFCA and FCA head data. **D)** Pseudotime and cellular composition of ISC and ISC-differentiated cell types. ISC, intestinal stem cell; EB, enteroblast; EC, enterocyte; EE, enteroendocrine cell; a. EC, anterior EC; p. EC, posterior EC; diff. EC, differentiating EC. **E)** Changes of cellular composition during aging. Each dot represents one cell type. Each color compares one aged sample and the 5d sample. Dot sizes reflect the nuclear numbers of the corresponding cell type from the aged population. Tissue origins are indicated. **F)** Comparison of the number of nuclei of the fat body from young and old flies. Nuclei are stained by DAPI and counted in each fly. The nuclear number is significantly increased in the 50d population (t-test, 50d vs. 5d, P value=0.0023). Error bar, standard deviation (SD). **G)** Representative confocal images showing nuclei in young and old fat body cells. The membrane is labeled by *cg-GAL4 > UAS-CD8GFP*. Nuclei are stained by DAPI and the LamC antibody. **H)** Fat body cells with segregating nuclei stained by pH3, DAPI, LamC, and GFP. **I)** Indirect flight muscle stained with cleaved-Caspase3 antibody, DAPI, and Phalloidin. Cleaved-Caspase3 signals are significantly increased in the aged population (t-test, 50d vs. 5d, P value<0.0001). Median numbers indicated.

For complex tissues such as the brain, more cell types can emerge when significantly more cells are sequenced (*9*–*11*). Indeed, 17 new neuronal cell types emerged after combining young and old head data (**Fig. 2C)**. Among them, 4 types are GABAergic neurons (*Gad1+*), 2 are Glutamatergic neurons (*VGlut+*), and the remaining 11 types are cholinergic neurons (*VAChT+*) (**fig. S10**).

Next, we assessed the reliability of AFCA data to investigate age-related changes in specific cell types. We focused on the fly gut as a case study, where the somatic stem cell lineage and its aging have been well characterized (*12*). In a healthy young fly gut, intestinal stem cells (ISCs) maintain gut homeostasis through proper proliferation and differentiation. In old flies, ISCs exhibit a high proliferation rate, and their daughter cells, enteroblasts (EBs), do not properly differentiate into mature enterocytes, leading to a dysplasia phenotype (*13*–*15*). We first extracted six major gut cell types and performed pseudotime trajectory analysis (*16, 17*). There is a significant increase of ISCs and EBs along with a decrease of fully differentiated enterocytes, consistent with previous in vivo studies (**Fig. 2D and fig. S11**) (*18*) and we could identify genes that showed different dynamic patterns between young and old flies (**fig. S11E**). In summary, the detailed annotations in our AFCA data offer a valuable resource to explore cell type- and tissue-specific aging signatures.

### Cell Composition Changes During Aging

In complex organisms, aging can affect cellular composition in different ways, such as changing stem cell proliferation or differentiation processes, altering cell identity, or inducing cell death. We assessed whether and how aging impacts cellular composition across the whole fly. Note that our measurements are based on nuclei composition. Since the nuclei are extracted from the whole head and body with minimal sampling bias, the ratio for each cell type in our sequencing data should largely reflect their composition in vivo.

To perform reliable analyses, we focused on cell types that have more than 500 nuclei in total (112 cell types after filtering). We then calculated the composition changes by comparing three older ages (30d, 50d, 70d) to young flies (5d). The top 5 increased cell types are cardia cells (proventriculus from the gut), fat body cells, male accessory gland main cells, EBs, and adult salivary gland cells (**Fig. 2E**). Comparing two consecutive ages showed similar results (**fig. S12**). Age-related increases of proventriculus cells, male accessory gland main cells, and EBs have been reported previously (*19*–*21*), confirming the quality of our data and analysis. Fat body cells are one of the most abundant cell types in *Drosophila*. They are polyploid, filled with lipid droplets, and tightly attached to the abdominal cuticle. These features make it difficult to isolate them or to compare their compositions using traditional methods. To our knowledge, an increase of fat body nuclei in old flies has not been reported. We were able to validate this observation (**Fig. 2F**).

Fat body cells are postmitotic cells, and no adult stem cells or progenitors have been reported for regeneration (*22*). To examine why fat body nuclei were increased in old flies, we first checked the number of nuclei within single cells using a fat body-specific GAL4 driving cell membrane GFP. Many aged fat body cells exhibited an increase in cell size and contained multiple nuclei per cell (**Fig. 2G**). The multinucleated phenotype can potentially be caused by cell membrane fusion as reported in other cell types (*23*), but it cannot explain the increase in the number of nuclei (**Fig. 2, E and F**). It has been reported that polyploid enterocytes from the fly gut can undergo nuclear cleavage without mitosis (a process termed amitosis) (*24*). To test this possibility, we performed immunohistochemistry to detect the nuclear lamina protein, Lamin C (LamC), and a mitosis marker, Phospho-Histone H3 (pH3). We did not detect any mitotic events from more than 60 flies across different ages, but we did observe many cases where two nuclei were localized very close to each other and they were negative for the mitotic marker (**Fig. 2H**). 3D reconstruction of confocal images confirmed that these nuclei were present in the same cell without a separating cell membrane (**Movies S1-S3**). Such events were captured across different ages (**fig. S13**). Together, these data suggest that fat body cells undergo nuclear division without cytokinesis across different ages, leading to multinucleated cells and an increase in the number of nuclei in older flies.

Among the top 5 decreased cell types are three types of muscles – indirect flight muscle, visceral muscle, and other muscle cells (mostly skeletal muscle) (**Fig. 2E**). Loss of muscle mass and strength, termed sarcopenia, is a conserved aging phenotype across different mammals, including humans (*25*). Our in vivo staining data confirmed the age-related degeneration of indirect flight muscles as well as a significant increase of an apoptosis marker Caspase3 in old flies (**Fig. 2I and fig. S14**), consistent with a previous study (*26*). Germline cells also showed a significant decrease (**Fig. 2E, fig. S12**), presumably contributing to the decline of fecundity in old flies. The majority of the cell types from the head, mostly neurons, showed minimal cellular composition changes (**Fig. 2E and fig. S12**).

### Differentially Expressed Genes (DEGs)

Altered gene expression is another consequence of aging (*27*). To assess such changes, we performed differentially expressed gene (DEG) analysis between young and old flies and ranked cell types based on the number of DEGs (**Fig. 3A and fig. S15A**). Again, we focused on 112 cell types with more than 500 cells for a reliable analysis. In the body, the cell type with the highest number of DEGs was the fat body, while the most affected cell type in the head was the outer photoreceptor (**Fig. 3A**).

**Figure 3.**
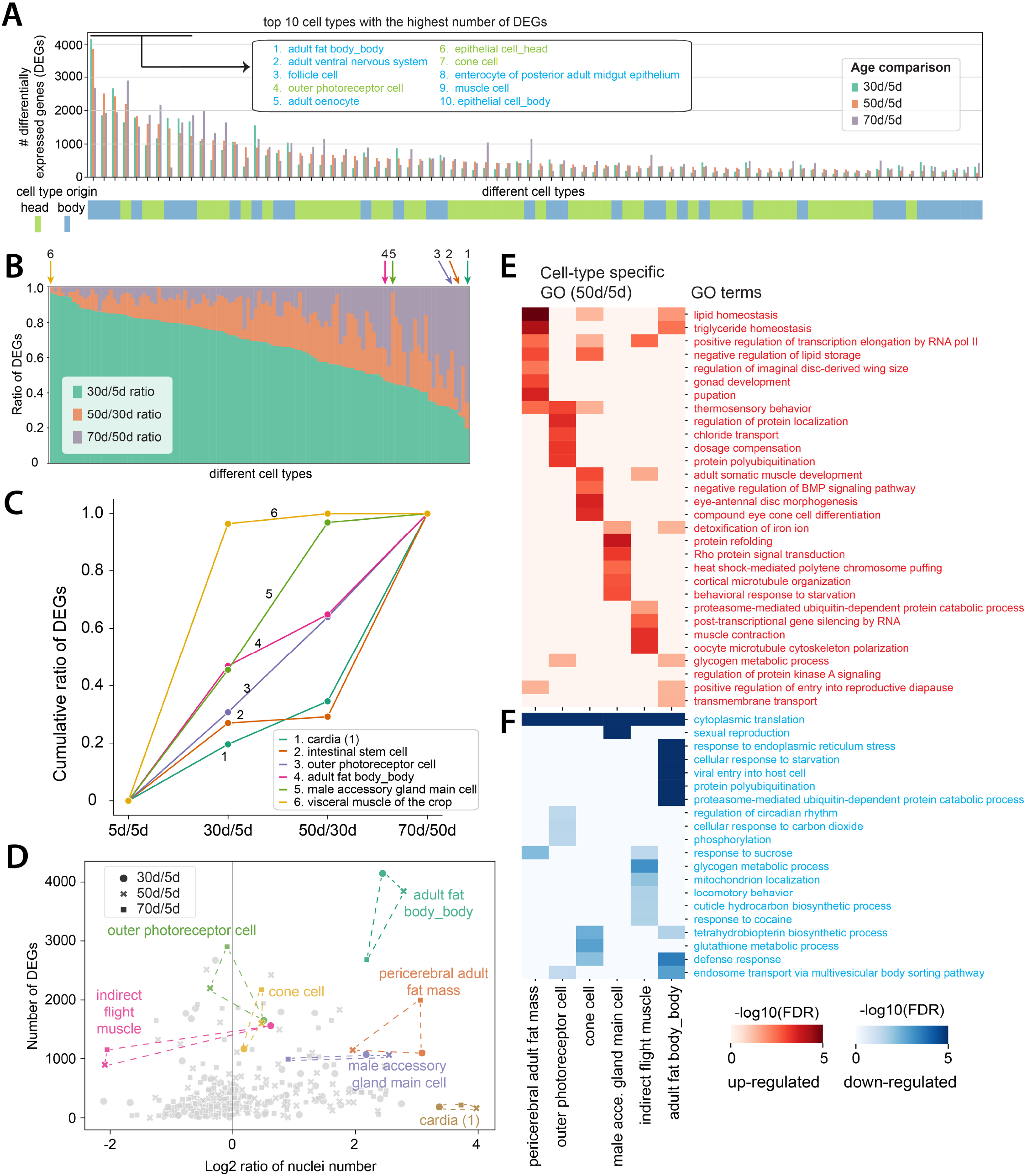
Differentially expressed genes (DEGs) **A)** Number of DEGs from different cell types. Each age group is compared with the 5d population. Each line shows the number of DEGs from the indicated age comparison. Cell types are ranked by DEG numbers from high to low (50d vs 5d). The top 10 cell types are indicated. **B)** Ratio of DEGs from each age comparison. Arrows point out representative cell types that are further compared in **Fig. 3C**. **C)** Cumulative ratios of DEGs from cell types indicated in **Fig. 3B**. **D)** Combination of DEG number and change of nuclear number illustrate different aging patterns. **E-F)** Top 5 cell type-specific GOs from selected cell types. **E)** GOs enriched in the selected cell types based on up-regulated DEGs. **F)** GOs enriched in the selected cell types based on down-regulated DEGs.

To explore the dynamics of cell type-specific changes, we further examined the time window during which cell types change the most by computing DEG numbers between two neighboring ages (**fig. S15B**) and normalizing their ratios (**Fig. 3B and fig. S16**). This analysis revealed several interesting insights (**Fig 3, B and C**). Specifically, about 80% of the cell types showed major changes (> 50% of DEGs) during the first time window, suggesting that 30d old flies have captured a large portion of age-related gene changes. Some cell types, like the male accessory gland, showed minimal changes in the last time window, indicating these cell types reach their maximum transcriptomic changes around 50d. However, 5.3% of cell types showed dramatic changes (> 50% of DEGs) in the last time window, such as intestinal stem cells and cardia cells, suggesting they age at a slower rate during the first 50d. Other cell types showed a steady change, such as outer photoreceptors and fat body cells. Hence, this analysis indicates that different cell types age at different rates and exhibit unique patterns of gene expression changes.

We compared DEGs from AFCA with those from the aging fly brain study (8) and found they are well correlated (**fig. S17**).

By integrating cellular composition changes and DEG analysis, we determined which cell types were affected by those two parameters (**Fig. 3D**). Significantly affected cell types (‘outliers’) fell into four categories: i) cells showing changes for both, such as fat body cells from the body, pericerebral adult fat mass (fat cells from the head), and male accessory gland main cells; ii) cells showing high DEGs but minimal composition changes, such as outer photoreceptors and cone cells, consistent with a previous study reporting that the age-related fly visual decline is not due to the loss of photoreceptors (*28*); iii) cells showing decreased nuclear number and a moderate number of DEGs, such as indirect flight muscles; iv) cells showing increased nuclear number but minimal DEGs, such as cardia.

Next, we performed sex-related analysis. We first observed that female marker yolk protein genes (Yp1, Yp2, and Yp3) showed a significant decrease during aging in most female cell types, while the male markers (roX1 and roX2) maintained high expression levels in most male cells (**fig. S18, A and B**). In addition, some genes, like Lsd-2 and CG45050, showed different trends between males and females with age (**fig. S18C**), suggesting aging impacts male and female cells differently. It has been shown previously that different cell types exhibit different DEGs between males and females at a young age (*7*). We next checked how these numbers change during aging. Generally, if one cell type showed a high or low number of DEGs at a young age, it maintained the high or low number during aging (**fig. S19A**). However, some cell types showed age-specific sex differences. For example, three cell types from the head, pericerebral adult fat mass, skeletal muscle, and hemocyte, all showed few DEGs in young flies, but many DEGs in the old (**fig. S19, B and C**). In contrast, three glial populations showed a significant decrease of DEGs with age. We also checked how sex affects the DEG number and cell composition and found that these two features highly correlate between male and female flies (**fig. S19, D-G**).

### Analysis of Gene Pathways

Next, we asked which genes and pathways are enriched in DEGs. Gene ontology (GO) analysis was performed for both up- and down-regulated genes from the 50d/5d dataset (**fig. S20**). Most GO terms were cell type-specific (**fig. S20A**): only 20% GO terms were shared by >5 cell types for down-regulated genes and 40% for up-regulated genes. Interestingly, we found that one GO term from down-regulated genes, called ‘cytoplasmic translation’, was shared by almost all cell types (**fig. S20C**). Cytoplasmic translation refers to the ribosome-mediated process for protein synthesis. Many transcripts of genes encoding ribosomal proteins (RPs) were decreased across cell types, consistent with previous studies (*29*). There were no globally shared GOs among the up-regulated genes (**fig. S20B**). Instead, they were restricted to specific groups of cells, like signal transduction seen in neuronal types and protein phosphorylation enriched in different non-neuronal cells.

We next focused on GOs enriched within a few cell types to understand the cell type-specific regulations (**Fig. 3, E and F**). Fat body cells from body and head (pericerebral adult fat mass) shared metabolic-related GO terms from up-regulated genes, such as lipid homeostasis and triglyceride homeostasis (**Fig. 3E**), reflecting common metabolic changes in these tissues. For indirect flight muscles, a reduction of locomotor behavior was observed and is likely to be caused by muscle degeneration (**Fig. 3F**) (*26*). Also, reproduction-related genes were strongly decreased in male accessory gland main cells (**Fig. 3F**), consistent with the decline in reproductive ability in males (*30*). In summary, we observed that genes involved in “cytoplasmic translation” are commonly decreased during aging in many cell types, whereas most other GO terms showed cell type-specific patterns.

### Aging Clock to Predict the Biological Age

To predict the biological age of an animal or human, a number of different aging clock models have been recently developed using epigenetic markers and transcriptomic data (*8, 31, 32*). Here, we investigated whether our snRNA-seq data can be used to develop aging clocks. To perform a more accurate prediction, we focused on cell types that have more than 200 cells at each age point (64 cell types). For each cell type, we trained a regression model (*33*) to predict age. We measured predictive performance using the coefficient of determination or R^2^ (**Fig. 4A**). The average performance across all cell types was high (average R^2^=0.79 for body and 0.84 for head; **fig. S21**).

**Figure 4.**
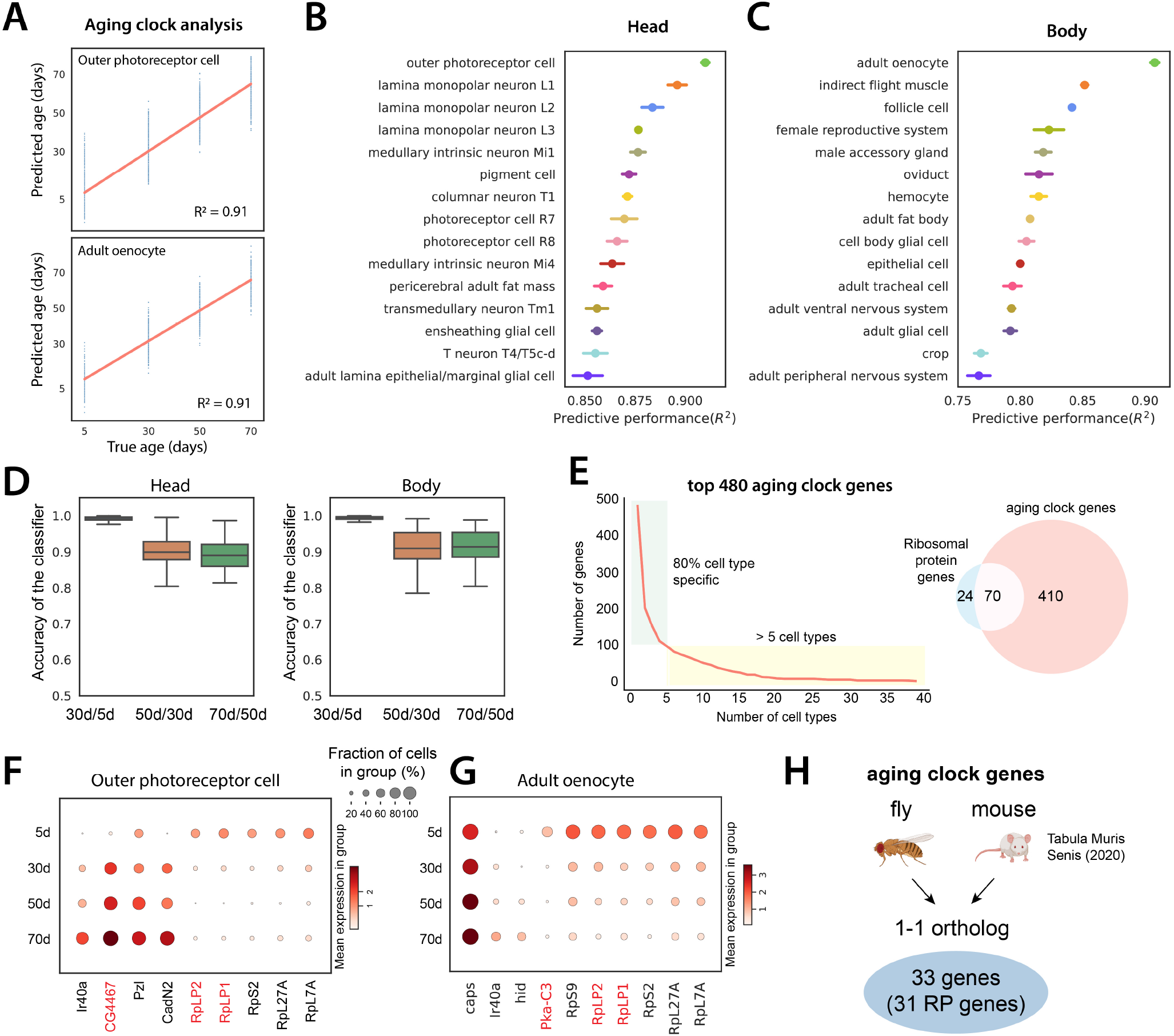
Aging clock analysis. **A)** Example of aging clocks for outer photoreceptor cells and adult oenocytes. Redline is the fitted regression line. Blue dots represent individual predictions where each dot corresponds to one cell. We measure performance as the proportion of the variance for an age variable that’s explained by transcriptome (R^2^). **B, C)** Predictive performance of cell type-specific aging clocks for head and body cell types. 15 cell types with the highest scores are shown. Error bars are estimated as a SD over 5 runs. **D)** Accuracy of logistic regression models trained to distinguish transcriptome between two consecutive time points. Boxplots show the distribution across head cell types (left) and body cell types (right). **E)** Number of aging clock genes as a function of the number of cell types (left). 80% of genes appear in less than 5 cell types. Out of 480 genes identified as aging clock genes, 70 encode RP genes (right). **F-G)** Examples of aging clock genes for outer photoreceptor cell F) and adult oenocyte G). Cell type-specific genes that appear in less than 5 cell types are shown in black color, while genes that appear in at least 5 cell types are marked in red color. **H)** Aging clock genes identified in flies and mice. 33 genes are 1-1 orthologs between the two species, and 31 of these are RP genes.

Outer photoreceptor cells and oenocytes showed the highest predictive scores in head and body, respectively (**Fig. 4, B and C**). *Drosophila* oenocytes perform liver-like functions including lipid storage and metabolomics functions, similar to fat body cells, and also produce important cuticular pheromones (*34*). We confirmed that high scores were not caused by higher nuclear numbers (f**ig. S22A**). We further asked whether there is a difference in predictive performance between different time windows and found that the largest transcriptomic differences are present between the first two ages (**Fig. 4D**).

Next, we focused on identifying aging clock genes that are used to predict the age. First, we found that most aging clock genes are used in a cell type-specific manner (**Fig. 4E**). We found that 70 out of 94 fly ribosomal protein (RP) genes were identified as aging clocking genes, showing age-related reductions in different cell types (**Fig. 4, E-G**). This data is consistent with our previous GO analysis where ‘cytoplasmic translation’ is reduced in almost all cell types (**fig. S20C**). Decreased protein translation, which can be caused by the reduction of RPs, is a prevalent feature of aging (*35*). To know what possible transcription factors regulate the expression of RPs, we utilized the regulon information from FCA (*7, 36*) and identified several transcription factors regulating RP genes (**fig. S23**). Our data suggest that the reduction of ribosomal expression contributes to the age-related decrease of protein synthesis. To further investigate the relationship of aging clock genes across species, we identified aging clock genes from Mouse Aging Cell Atlas (*27*). Among 33 overlapping aging clock genes from the fly and mouse, 31 genes encode RPs (**Fig. 4H and fig. S24 and S25**).

### Comprehensive Aging Features

Through the above analyses, we noticed that different cell types are sensitive to different aging features. To gain a better understanding of cell type-specific aging, we investigated more aging features, including expressed gene number or transcript number (measured by UMI) changes, and decline of cell identity (**Fig. 5A**).

**Figure 5.**
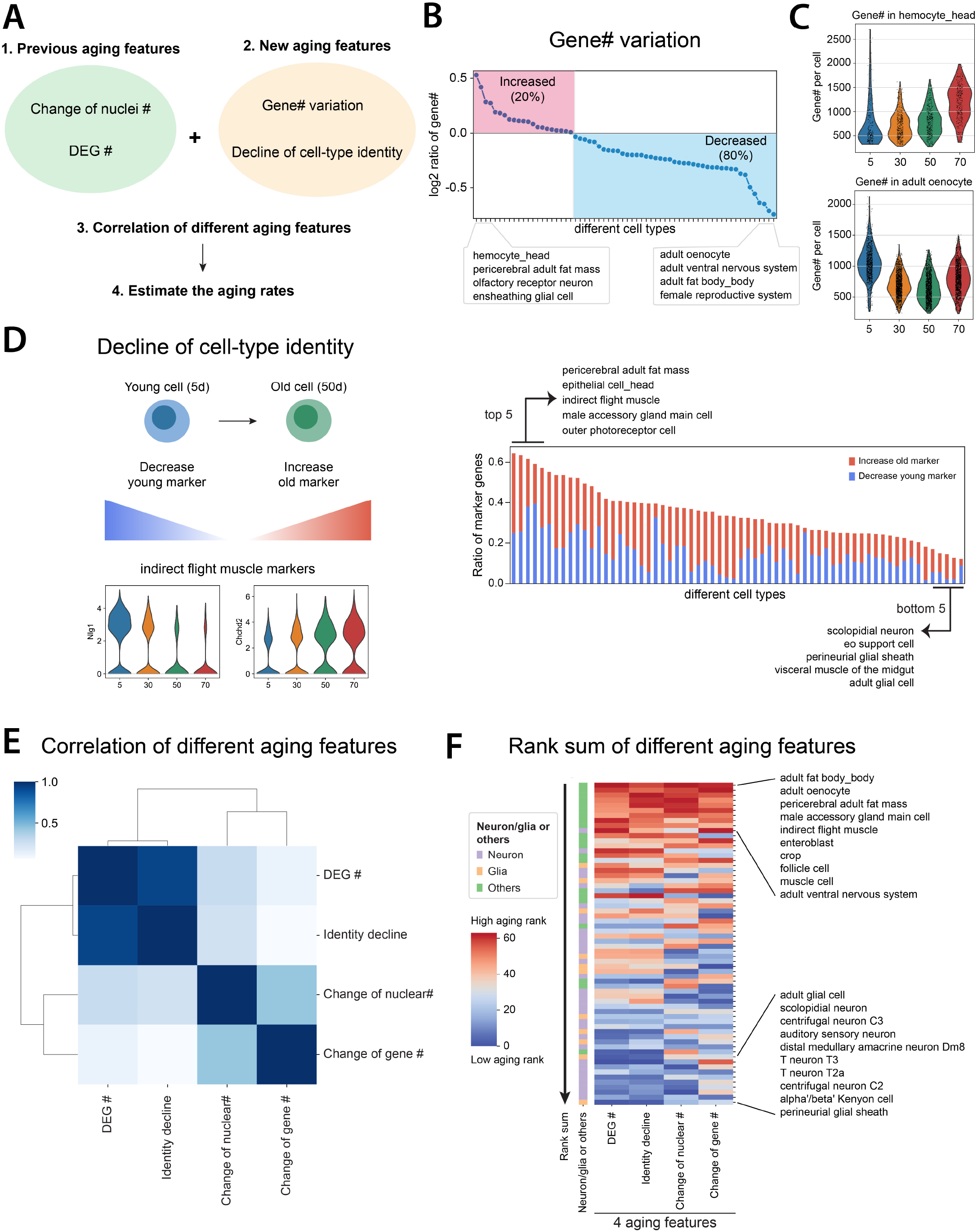
Systematic comparisons of different aging features. **A)** Flowchart of comparing different aging features. **B)** Expressed gene numbers per cell from each cell type are compared between 50d and 5d flies. The red block shows cell types with increased expressed gene numbers, while the blue block includes cell types with decreased ones. **C)** Two cell types, hemocyte from the head and oenocyte, have the highest increase and decrease of expressed gene numbers per cell. **D)** Decline of cell identity during aging. The left panel illustrates two different mechanisms of decreasing cell identity. The right panel shows the ratio of marker genes decreasing cell identity. Each line in the right panel represents one cell type. **E)** Spearman’s correlation of different aging features. **F)** Rank sums of different aging features. The heatmap shows the overall rank sum scores from different cell types. High aging ranks are shown in red, while low aging ranks are shown in blue. Neuron-or glia-related cell types are indicated beside the heatmap.

Similar to the aging clock analysis, we only focused on cell types with at least 200 cells from each age. Gene and UMI numbers were previously found to decrease in the old fly brain (*8*). Consistently, we observed that the CNS neurons decrease the gene and UMI numbers during aging (**fig. S26A**). To understand whether such a reduction is a general aging feature or not, we examined all other cell types. The overall trend of gene and UMI numbers was largely consistent (**fig. S26B**). We found that around 80% of cell types decreased expressed gene numbers, but 20% increased the expressed gene number, suggesting that aging affects expressed gene numbers in a cell type-specific manner (**Fig. 5B)**. Head hemocytes and pericerebral adult fat mass increased most in expressed gene numbers, while oenocytes, ventral nervous system, and fat body decreased the most (**Fig. 5C and fig. S26C**). We confirmed this was not caused by sequencing depth (**fig. S26D**). Even though head and body fat body cells both increased the nuclear number (**Fig. 3D**), their expressed gene numbers showed opposite trends.

Loss of cell-type identity has been shown to occur during aging for certain cell types (*12, 37*). However, how cell identity is changed across the entire organism during aging remains uncharacterized. To assess whether established cellular gene expression programs that define cell identity are changed during aging, we developed a measurement to combine two ratios – loss of original markers and gain of new markers – by comparing old to young populations (**Fig. 5D**, left panel). We then ranked cell types by their cell identity decline score (**Fig. 5D**, right panel; **fig. S27**). The gene, *Nlg1*, which was used as a marker gene to annotate the indirect flight muscle, showed a dramatic decrease with age, similar to a number of other young marker genes (**Fig. 5D and fig. S6A**). Meanwhile, many other genes’ transcripts, like *Chchd2*, began to appear in this cell type with aging. Other than the indirect flight muscle, pericerebral adult fat mass and epithelial cell from the head were also found to largely decrease cell-type identity (**fig. S28**).

### Correlation and Ranks of Aging Features

We examined four different aging features: cell composition changes, DEGs, change of expressed gene numbers, and cell identity decline. To understand the overall correlation of them, we ranked each feature from the least age-related change to the most. After integrating four aging features, their correlations were compared using Spearman’s correlation and clustered using the correlation scores (**Fig. 5E**). Among those features, DEG number and decline of cell identity were highly correlated, suggesting cell types with high numbers of DEGs would usually fluctuate in the expression of marker genes. On the other hand, changes of nuclear number, and expressed gene number were more correlated with each other.

Next, we summed different feature ranks and sorted cell types by the total rank sums, a higher rank indicating a more “aged” cell type (**Fig. 5F and fig. S29**). Interestingly, the top three cell types include three adipose cell types, oenocytes, fat body cells, and pericerebral adult fat mass, suggesting that those cells age faster than other cell types. Following them are male accessory gland main cells, indirect flight muscle, and enteroblast. Generally, neurons and glia from the nervous system age slower compared to other cell types (**Fig. 5F**). In summary, our analysis provided the first exhaustive analysis of different aging features and revealed the aging rates of different cell types across the entire organism.

## Discussion

The cell atlas approach is emerging as a powerful tool to systematically study aging in different organisms, including worms (*38, 39*), mice (*27*), and humans (*40*). A recent study performed cross-species analysis with scRNA-seq data from three species, including *Drosophila melanogaster* (*41*). Our AFCA provides a complementary dataset and thorough analyses for studying aging features across the whole organism.

One interesting observation is the significant increase of fat body nuclei during aging. *Drosophila* fat body is a liver-like tissue that stores fat and serves as a detoxifying and immune-responsive organ. Adult fat body cells are postmitotic polyploid cells without a stem cell or progenitor population (*22*). How do they increase their nuclear number? Our observations suggest that these polyploid cells increase their number of nuclei by nuclear cleavage without cytokinesis, forming multinucleated cells. To complete karyokinesis, the nuclear envelope needs to be reassembled or reorganized. Fat body cells have been shown to decrease their nuclear envelope integrity due to the loss of Lamin-B during aging (*42*), suggesting that the loss of nuclear envelope integrity may be associated with the multinucleated phenomenon. On the other hand, multinucleated cells have also been observed in the fly male accessory gland (*43, 44*), and subperineurial glial cells (*45*), suggesting multinucleated cells play specific roles in *Drosophila*. Other than *Drosophila*, multinucleated cells have also been observed in other species, including mushrooms (*46*), plants (*47*), and human and mouse liver cells (*48, 49*). Understanding the formation and regulation of multinucleated cells in the aging organism may provide new insights into an evolutionarily conserved phenomenon, as well as into the potential roles of multinucleated cells in age-related diseases.

We focused on four different aging features. Although these features cover several key aspects of age-related changes, the picture is incomplete. Additional aging measurements may reveal more specific aging patterns. For example, we investigated the change of alternative polyadenylation (APA) patterns, which can reflect short or long 3’ UTR usage for different isoforms (*50*), and found that neuronal extended 3’ isoforms were progressively depleted during aging. This phenotype is more obvious at 70d and more pronounced in females than in males (**fig. S30**). These data imply a global change in post-transcriptional regulation in aging neurons. Thus, future analysis may help elucidate additional aging patterns.

One major goal of this study is to characterize how different cell types age across the organism. Our analysis using different aging features provides several interesting insights. First, different cell types have very distinct aging patterns. For example, the ventral nervous system showed high ranks for 3 aging features but a low rank for the change of nuclear number, while scolopidia neurons showed low ranks for 3 aging features but high changes in expressed gene number (**Fig. 5F**). This observation is not unexpected considering that each cell type carries a unique function. Second, we observed a divergence in the contribution of individual cell types to a tissue’s aging. For example, in the female reproductive system, follicle cells were ranked very high (8th of 64 cell types), but germline cells were ranked near the bottom (41st of 64 cell types) (**fig. S29A**). This indicates that age-related declines of female fertility may be due to the aging of follicle cells. Third, the top-ranked cell types include all adipose cells. This is surprising and we do not fully understand the underlying mechanisms. It may be linked to the fact that these cell types play multiple critical roles in different physiological conditions, such as lipid storage and metabolism, immune responses, and interorgan communication with muscles and gut (*22, 51, 52*).

## Materials and methods summary

The study involved collecting fly head and body samples from wild-type F1 flies from the cross between female W^1118^ and male Oregon R (OreR). Samples from different ages were dissected and stored at -80°C after flash-freezing with liquid nitrogen. The snRNA-seq was prepared using the FCA protocol (*7*), and each age group had 12 samples with 6 females and 6 males. The libraries were sequenced using NovaSeq 6000 (Illumina).

FASTQ files were filtered for index-hooping reads using 10x Genomics index-hopping-filter software. The Cell Ranger (version 4.0.0) index was built using the *Drosophila melanogaster* genome (FlyBase r6.31) and the pre-mRNA GTF established by the FCA. Cell Ranger Count was used to estimate the nuclei number and gene expression from each nucleus. Nuclei from different sexes and ages of flies were integrated using Harmony, and cell-type annotations from FCA samples were transferred to AFCA samples using a cluster-centered or machine-learning-based method, followed by manual corrections.

For cell composition, age-specific ratios were obtained by dividing the number of nuclei in each cell type from one specific age by the total number of nuclei in the corresponding age. The relative ratios were then compared between the young (5d) and old populations. Wilcoxon Rank Sum tests were used to compare gene expression between different ages or sexes, and genes with FDR lower than 0.05 were considered to be differentially expressed. The differential expression of the top 200 marker genes from 5d and 50d samples was used to estimate declines of cell-type identity. Aging rates of each cell type were calculated by integrating different aging features and ranking the sums of aging ranks from high to low.

## Acknowledgments

We thank Stein Aerts and Jasper Janssens for helping preprocess the data and their constructive advice on annotation, Norbert Perrimon for suggestions during the initiation of this project, Laura Buttitta for comments on polyploid phenotypes, Andrey Parkhitko for discussion on metabolic pathways. We thank Ao-Lin Hsu (NYCU) for supporting C.Y.L., Claude Desplan for supporting Y.C.C., Chuangye Qi, Madeline Burns, and all other lab members for comments and feedback along the project. We also thank FCA Consortium and the fly community for their enthusiastic support. **Funding:** M.B. acknowledges the EPFL support. T.J. was supported by the Baylor College of Medicine Cancer and Cell Biology program grant T32 (GM136560). H.J.B. is supported by NIH/NIA, R01 AG07326, the Huffington foundation, and the endowment of the Chair of the Neurological Research Institute. L.L. is an investigator of the Howard Hughes Medical Institute and supported by NIH (R01-DC005982). L.L. and S.Q.R. are supported by Wu Tsai Neuroscience Institute (Neuro-omics program). This work was supported by CZ Biohub (S.R.Q.). S.L. was supported by a training award from the NYSTEM contract #C32559GG and the Center for Stem Cell Biology at MSKCC. Work in the E.C.L. group was supported by NIH/NINDS (R01-NS083833) and NIH MSK Core Grant P30-CA008748. Y.C.C. was supported by the MacCracken Program at New York University, by a NYSTEM institutional training grant (Contract #C322560GG), and by Scholarship to Study Abroad from the Ministry of Education, Taiwan. C.Y.L. is supported by the NSTC grant from Taiwan (110-2311-B-A49A-501-MY3). H.L. is a CPRIT Scholar in Cancer Research (RR200063), and supported by NIH (R00AG062746), the Longevity Impetus Grant, and Ted Nash Long Life Foundation. **Author contributions:** Conceptualization: T.C.L., L.L., H.J., S.R.Q., H.L. Sample preparation: N.K., X.T.C., H.L. Sequencing: S.S.K., Y.Q., R.C.J., H.L. Computational analysis: T.C.L., M.B., J.L., T.J., D.W. Data portal and AFCA website: J.C., T.C.L., S.D., S.H.D., A.P. APA analysis: S.L., Y.C.C., E.C.L. Data validation: Y.J.P., H.J.B., T.J., Y.Q., N.A., C.Y.L Writing: T.C.L., M.B., H.L. Review and Editing: All authors. Supervision: H.J., S.R.Q., H.L. Funding Acquisition: H.J., S.R.Q., H.L. **Competing interests:** H.J., N.K., and X.T.C. are employees of Genentech, Inc. The other authors declare no competing interests. **Data and materials availability:** All data are available for querying at https://hongjielilab.org/afca. Raw FASTQ files and processed h5ad files, including expression matrices and cell-type annotations, can be downloaded from NCBI GEO: GSE218661. Count matrices and analysis codes for generating the figures are available from Zenodo (*53*). FCA data were downloaded from https://www.flycellatlas.org/.

## Supplementary Materials

Materials and Methods

Supplementary Text

Figs. S1 to S30

References (*54*-*59*)

Movies S1 to S3

