## Supplemental Figures for "Aging Fly Cell Atlas Identifies Exhaustive Aging Features at Cellular Resolution"

#### Materials and Methods

##### Fly sample and lifespan

Fly head and body samples were from F1 of two wild type fly stains: female $w^{1118}$ and male Oregon R (*OreR*). Flies were maintained on standard cornmeal medium at 25 ℃ with 12-hour light–dark cycle. For the lifespan experiment, progenies were collected 3 days after the first fly hatched. Flies were allowed to mate for 3 days. Then male and female flies were separated into different cages (120 flies per cage) for lifespan or sample collection at 25°C. Fly food was changed every 2 days. Dead flies were counted every 2 days. Lifespan curves were generated in Excel.

##### Single-nucleus RNA sequencing

Fly heads and bodies dissected at different ages were put into 1.5ml RNAase-free Eppendorf tubes, flash-frozen in liquid nitrogen, and then stored at –80 ℃. We put 100 heads or 50 bodies in each tube. Single-nucleus suspensions were prepared following the protocol described in the Fly Cell Atlas study (*7*). Next, we collect nuclei by FACS. We used the SONY SH800 FACS sorter for collecting nuclei. Nuclei were stained by Hoechst-33342 (1:1000; >5min). Since polyploidy is common for many fly tissues, we observed different populations of nuclei according to DNA content (Hoechst signal). In order to include all cell populations with different nuclear sizes, we have included all nuclear populations from the FACS. Nuclei were collected into a 1.5ml tube with 200ul 1x PBS with 0.5% BSA as the receiving buffer (RNase inhibitor added). For each 10x Genomics run, 100k–400k nuclei were collected. Nuclei were spun down for 10 min at 1000g at 4 ℃ and then resuspended using 40ul or desired amount of 1x PBS with 0.5% BSA (RNase inhibitor added). 2ul nucleus suspension was used for counting the nuclei with hemocytometers to calculate the concentration. When loading to the 10x controller, we always target 10k nuclei for each channel. We observed that loading 1.5 folds more nuclei as recommended by the protocol allowed us to recover about 10k cells after sequencing. For example, if the concentration is 1500 nuclei per ul for one sample, we treat it as 1000 nuclei per ul when loading to the 10x controller.

Next, we performed snRNA-seq using the 10x Genomics system with 3' v3.1 kits with the following settings. All PCR reactions were performed using the Biorad C1000 Touch Thermal cycler with 96-Deep Well Reaction Module. 13 cycles were used for cDNA amplification and 16 cycles were used for sample index PCR. As per 10x protocol, 1:10 dilutions of amplified cDNA and final libraries were evaluated on a bioanalyzer. Each library was diluted to 4 nM, and equal volumes of 18 libraries were pooled for each NovaSeq S4 sequencing run. Pools were sequenced using 100 cycle run kits and the Single Index configuration. Read 1, Index 1 (i7), and Read 2 are 28 bp, 8 bp and 91 bp respectively. A PhiX control library was spiked in at 0.2 to 1% concentration. Libraries were sequenced on the NovaSeq 6000 Sequencing System (Illumina).

For each age, we sequenced the head for 12 runs (6 for male and 6 for female) and body for 12 runs (6 for male and 6 for female). To collect enough nuclei for 6 runs, we used 100 heads and 50 bodies. For three ages, we sequenced 24*3 = 72 runs. We have recently published a protocol describing the single-nucleus RNAseq in *Drosophila* and all the above steps have been detailed (*54*).

#### Sequencing read alignment

The raw FASTQ files were first filtered for the index-hopping reads by using the index-hopping-filter developed by the 10x Genomics (version 1.0.1). Before mapping reads, the *Drosophila melanogaster* genome (FlyBase r6.31) was indexed with a pre-mRNA GTF, which was established by FCA (*7*), using the Cell Ranger (version 4.0.0). FASTQ files removing the index-hopping reads were mapped to the *Drosophila melanogaster* genome (r6.31) using Cell Ranger Count (version 4.0.0) to generate the count matrix.

#### AFCA annotation

In the FCA dataset, there are 251 cell types annotated, many of which were characterized in dissected tissues. For cell types from the head, besides the whole head sequencing, there are only two dissected tissues: antennae and proboscis. So, most cell types from the aging head can be annotated based on the FCA whole head data. For cell types from the body, most cell type annotations are from tissues (only 33 annotated cell types from the FCA whole-body data). To annotate aging body data, we need to transfer annotations from the body and individual tissues (e.g., gut, wing, leg, testis, ovary, fat body, etc). Thus, the aging head and body data were annotated using slightly different strategies and we characterized them separately. For both, we first integrated FCA and AFCA datasets and corrected age and sex differences using Harmony (*55*). Next, we characterized cell types using two different approaches: co-clustering and supervised machine learning (ML)-based classification. Then, we validated those two approaches using cell type-specific markers and performed manual correction when necessary.

For the head data, FCA and AFCA head cells were co-clustered and 150 principal components (PCs) were selected for clustering. We followed the co-cluster-based approaches utilized in FCA brain-head integration (*7*) and annotations from the FCA cells were transferred to the clusters that contained at least 15% of cells with the same annotation. Alternatively, if more than 15% of the annotation-specific cells were enriched in the clusters, those clusters were assigned to the corresponding annotations. Different Leiden resolutions were applied for the transfer and the one with the maximum number of transferred annotations was chosen. If annotations from the two approaches were not consistent, cluster-dominant annotations would be assigned to the clusters. Besides cluster-based transfer, we also included a supervised ML method. We utilized the Logistic Regression classifier to transfer the FCA annotations to AFCA cells. FCA cells were used as the training dataset, while AFCA samples were considered as test data. By comparing the annotations from these two approaches, annotations were manually confirmed by examining the expression of cell type-specific genes to prevent the loss of transfer due to aging. Also, annotations were unified and denoised at the cluster level. In total, 91 cell types were annotated, including 17 newly identified neuron types.

To annotate AFCA body cells, FCA and AFCA body cells were first co-clustered using 50 PCs and 33 annotations from the FCA body were transferred to the AFCA body cells using approaches similar to the head transfer. To include annotations from different body tissues, we co-clustered cells from the FCA body, AFCA body, and all FCA tissues to identify clusters with tissue-specific annotations (**Fig. S4**). The cluster-based method enabled us to quickly identify tissue-specific annotations from integrated body and tissue cells. Clusters with tissue-specific annotations were chosen for further sub-clustering and tissue annotations were transferred similarly using cluster- and supervised ML-based approaches. Due to the similarity between some gut and malpighian tubule cell types, especially for the intestinal stem cell and renal stem cell (**Fig. S6C and S7**), we combined annotations from these two tissues together for the transfer. Some cell types, like muscle cells, were present in multiple tissues and we considered subsetting those cells based on the expression of cell-type markers, instead of selecting the annotation-enriched clusters (**Fig. S5**). After subsetting marker-positive cells, FCA annotations were similarly transferred. After transferring all tissue annotations, 72 cell types were annotated in the AFCA body.

**Gut cell type case study**

The analysis of the gut cells in the AFCA data was performed using SCANPY and scFates python packages (*56, 57*). The intestinal stem cell lineage was isolated from the data and includes intestinal stem cells, enteroblasts, adult differentiating enterocytes, enteroendocrine cells, anterior enterocytes, and posterior enterocytes. The pseudotime was generated by combining the young (FCA age 5d sample) and old (AFCA age 50d and 70d samples). To generate this plot, the partition-based graph abstraction (PAGA) function was applied to infer the connectivity of clusters then the ForceAtlas2 algorithm was implemented to spatially overlay the cells onto the connectivity plot generated by PAGA (*16, 17*). To compare the cell compositions of each age, the cells of different ages were plotted separately next to the pseudotime. The percentages of cells comprising the intestinal stem cell lineage were extracted from the data and then plotted in **Fig 2D**.

The changes in gene expression were generated using the scFates python package (*57*). Cells were isolated from each age and then run separately in the scFates pipeline. Briefly, scFates uses the ElPiGraph algorithm to learn the topography of the data. Then, the pseudotime was constructed from the graph inferred by the ElPiGraph algorithm. The genes differentially expressed along the trajectory were determined using a cubic spline regression model and then these results were compared to an unrestrained model. The Benjamini-Hochberg correction was then applied to adjust for multiple comparisons. A significantly altered gene along the trajectory between each age, Rbfox1, is shown in **Fig S11**.

**Fat body and indirect flight muscle staining**

To label the fat body tissue with GFP, cg-GAL4 (Bloomington Drosophila Stock Center (BDSC), #7011) virgins were crossed with UAS-unc84GFP (from Dr. Gilbert L. Henry) or UAS-CD8GFP (from Dr. Liqun Luo) males to obtain cg-GAL4>unc84GFP or cg-GAL4>CD8GFP flies. For indirect flight muscle labeling, Act88F-GAL4 (BDSC #38461) virgins were crossed with UAS-unc84GFP to obtain Act88F-GAL4>unc84GFP flies. Flies were cultured at 25°C to desired ages.

For fat body dissection, fly abdomen filets were prepared in 1X PBS using Vannas Spring Scissors (FST, 3mm Cutting Edge, 15000-00) and Dumont #55 forceps (FST, 11255-20). For indirect flight muscle dissection, flies in 1.5ml Eppendorfs tube were snap-freezed in liquid nitrogen and placed on the ice rack. After removing the head, leg, and wing, the abdomen was clamped with forceps and the thorax was cut in the sagittal section by a razor. Tissues were fixed with 4% paraformaldehyde for 20 min at room temperature (RT), washed with 1X PBS, blocked in 5% NGS blocking buffer (1XPBS, 0.3% Triton X-100, 5% NGS) for 2 h at RT, incubated with primary antibodies (in 5% NGS blocking buffer) overnight at 4°C, and then washed 5 times with 0.3% PBST (1XPBS, 0.3% Triton X-100) before incubation with secondary antibodies for 2 h at RT. Tissues were thoroughly rinsed in PBST, stained with DAPI(1:1000) for 15 min, washed with 1XPBS and mounted with SlowFade™ Gold Antifade Mountant (Thermo Fisher, S36936).

Mouse anti-LamC (1:100, Developmental Studies Hybridoma Bank (DSHB), LC28.26), chicken anti-GFP (1:1000, Aves Labs), rabbit anti-pH3 (1:1000, Cell Signaling Technology, #9701), rabbit anti-cleaved-Caspase3 (1:100, Cell Signaling Technology, #9661) were used as primary antibodies. For secondary antibodies, we used Cy™3 AffiniPure Donkey Anti-Mouse IgG (H+L) (1:250, Jackson ImmunoResearch, AB_2340813), Alexa Fluor® 488 AffiniPure Donkey Anti-Chicken IgY (IgG) (H+L) (1:250, Jackson ImmunoResearch, AB_2340375), Alexa Fluor™ 647 Donkey anti-Rabbit IgG (H+L) Highly Cross-Adsorbed Secondary Antibody (1:250, Invitrogen, A31573). Alexa Fluor 647 Phalloidin (1:250, Invitrogen, A22287) was used to stain F-actin in muscle.

Images were obtained with Leica STELLARIS 5 confocal microscope. Images were obtained as Z series with the same interval. Z series images were merged by ImageJ (Image-Stacks-Z projection-Max Intensity), and then the mean fluorescence intensity was measured (Analyze-Measure-Mean gray value) or the nuclear number was counted by Point Tool. Quantification graphs are generated by GraphPad Prism. P-values from unpaired t-test. Error bar, SD.

**DEG analysis**

To identify genes that are differentially expressed in the aged population, we applied the Wilcoxon Rank Sum test (rank_genes_groups() in SCANPY) to compare the gene expression between the aged (30d, 50d, and 70d) and the young population (5d). Genes with FDR lower than 0.05 would be considered to be differentially expressed.

**Change of cell composition**

Cell numbers of each cell type were counted and separated by age. To get the ratio of one cell type from one age, the cell numbers of a specific cell type and age were further divided by the total cell number of the corresponding age. The cell-type ratios from the aged population (30d, 50d, 70d) were further divided by ratios from the young population (5d) to get the relative ratios in the log2 scale.

**GO analysis**

In each cell type, aging DEGs, including up- or down-regulated genes, were applied for the Gene Ontology (GO) analysis using GOATOOLS (version 1.2.3) (*58*). The gene association file (version FB2022_04) was downloaded from FlyBase. Redundant GO terms were removed by REVIGO

(*59*). GO terms from Biological Process (BP) were mainly used for our analysis (**Fig. 3E and 3F**). GOs restricted within 5 cell types were considered as cell-type specific GOs (**Fig. 3E, 3F, and Fig. S20A**).

**Aging clock analysis**

For aging clock analysis, we focused on 64 head and body cell types that have at least 200 cells at each time point. For each cell type satisfying the condition, we then built aging clock models – regression models that predict age from the transcriptome. In particular, for each cell type we trained the elastic net model (*33*) by using cells' transcriptome as explanatory (independent) variables and age as the response (dependent) variable. We randomly split the dataset into a train and test dataset where we used 70% of cells to train the model, and 30% of cells to validate the model. We compared the predictions to the true age and measured performance of the model as $R^2$, *i.e.*, proportion of the variance for a dependent variable that's explained by an independent variable or variables. We repeated the analysis over five random seeds, resulting in different train and test data splits. To find genes that correlate with age, we first filtered all genes with zero coefficient in the regression model. For the remaining genes, we calculated the Pearson correlation between its value and age and retained those genes for which the correlation coefficient was higher than 0.3. This analysis resulted in 480 genes. We repeated the same analysis for the Mouse Aging Cell Atlas (Tabula Muris Senis) (*27*)**.**

To compare models between two consecutive stages, we trained three logistic regression models for each cell type to distinguish between (i) 5 days and 30 days old fly, (ii) 30 days and 50 days old fly, and (iii) 50 days and 70 days old fly. We split the data into train and test dataset by using 70% data for training the model and 30% data for evaluation by preserving the percentage of samples for each class (stratified sampling). We evaluated classification performance using accuracy.

**Identification of possible transcription factor(s) regulating the RP genes**

Head and body regulons identified by SCENIC were obtained from FCA (*7, 36*). TF regulons containing more than 5 RP genes were considered to be the potential transcription factors. RP genes were removed from the regulon of each transcription factor and the average expression of RP and regulon genes was calculated in each cell type. The average expression of RP and regulon genes was clustered to find regulons showing gene expressions similar to the RP genes.

**Cell-identity analysis**

Change in cell identity was based on the expression changes of marker genes between different ages. Cell-type markers from young (5d) and old animals (50d) were identified separately using the Wilcoxon Rank Sum test (rank_genes_groups() in SCANPY). The top 200 genes that are differentially expressed between cell types were defined as marker genes for each cell type. Expression of marker genes was further compared between young and old populations using the Wilcoxon Rank Sum test to identify differentially expressed markers (FDR < 0.05). Young marker genes with a significant reduction of expression would be considered to decrease the cell-type identity. On the other hand, old marker genes with a significant increase in expression were also considered to decrease the identity. Together, we used the average of these two ratios to indicate the change in cell identity.

**AFCA website**

The AFCA Website was developed with the Shiny package (v1.6.0) in R language (v4.0.5). We included three major datasets: Head dataset, Body dataset, and Head and Body dataset, where they contain the snRNA-Seq analytical results from head tissues, body tissues, and combined respectively. For the Cell type and Gene Expression tabs, the data processing and visualization were performed with the Tidyverse (v1.3.0) and ggplot2 (v3.3.3) packages in R. For the customized analysis tab, its features and functions were powered by the ShinyCell (v2.1.0) package.

**The rank sum of aging features**

We measured the absolute differences of each aging feature between the young (5d) and old (50d) population and ranked the cell-type differences. High ranks represented high aging differences, while low ranks indicated low differences between young and old populations. To have a better estimation, we focused on cell types with more than 200 nuclei from each age. Correlations of different aging features were compared using Spearman's correlation. To know which cell types showed high aging differences supported by different features, we summed the ranks of five different aging features and ranked the sums. High-rank sums of different aging features represented high aging differences and vice versa.

**Analysis of alternative polyadenylation (APA) analysis**

Our overall analysis strategy was based on our recent scAPA study of the FCA dataset (*50*). We used LABRATsc (https://github.com/TaliaferroLab/LABRAT) to quantify APA from the scRNA-seq data with current FlyBase *Drosophila melanogaster* gene annotation (version r6.45). Two modes were used for either single-cell level (cellbycell) or cell-type specific (subsampleClusters) quantification using '--mode' parameter. We used read coverage thresholds of at least 3 or 100 counts per gene for single-cell level or cell-type specific quantification respectively to minimize noise and false positives. LABRAT calculates ψ values reflecting 3' isoform usage, with "0" indicating exclusive usage of the most proximal pA site, while "1" indicates exclusive usage of the most distal pA site. Ψ values for each annotated cell-type, and for individual cells, were used for downstream analysis.

**Figs. S1 to S30**

Figure S1

**A** data processing steps

**Preprocessing**
- Removing index-hopping reads
- Removing ambient RNAs
- Removing doublets
- Removing low-quality cells

**Normalization**
- Normalize to the library size
- Log transformation

**Dimensionality reduction**
- PCA
- Batch correction using harmony
- Compute the neighborhood graph
- Embed the graph using t-SNE or UMAP

**Clustering**
- Leiden-based clustering
- DEG identification using Wilcoxon rank-sum test

**Annotation and data sharing**
- AFCA annotations
- Upload raw fastq files to NCBI
- AFCA website for data visulization

**B** data availability

AFCA

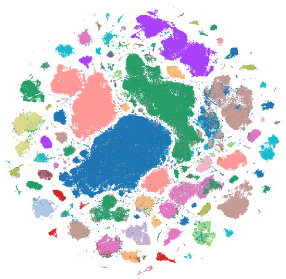

**raw data** → **NCBI GEO: GSE218661**

**codes** → **https://doi.org/10.5281/zenodo.7853649**

**visualization** → **CELLxGENE**

**visualization & analysis** → **AFCA website (See Fig. S2)**

**https://hongjielilab.org/afca/**

**Fig. S1. General pipeline of snRNA-seq analysis and data availability.**

A) Data processing steps, including preprocessing, normalization, dimensionality reduction, clustering, annotation, and data sharing. B) Data available from different sources, including NCBI, GitHub, CELLxGENE, and AFCA website.

### Figure S2

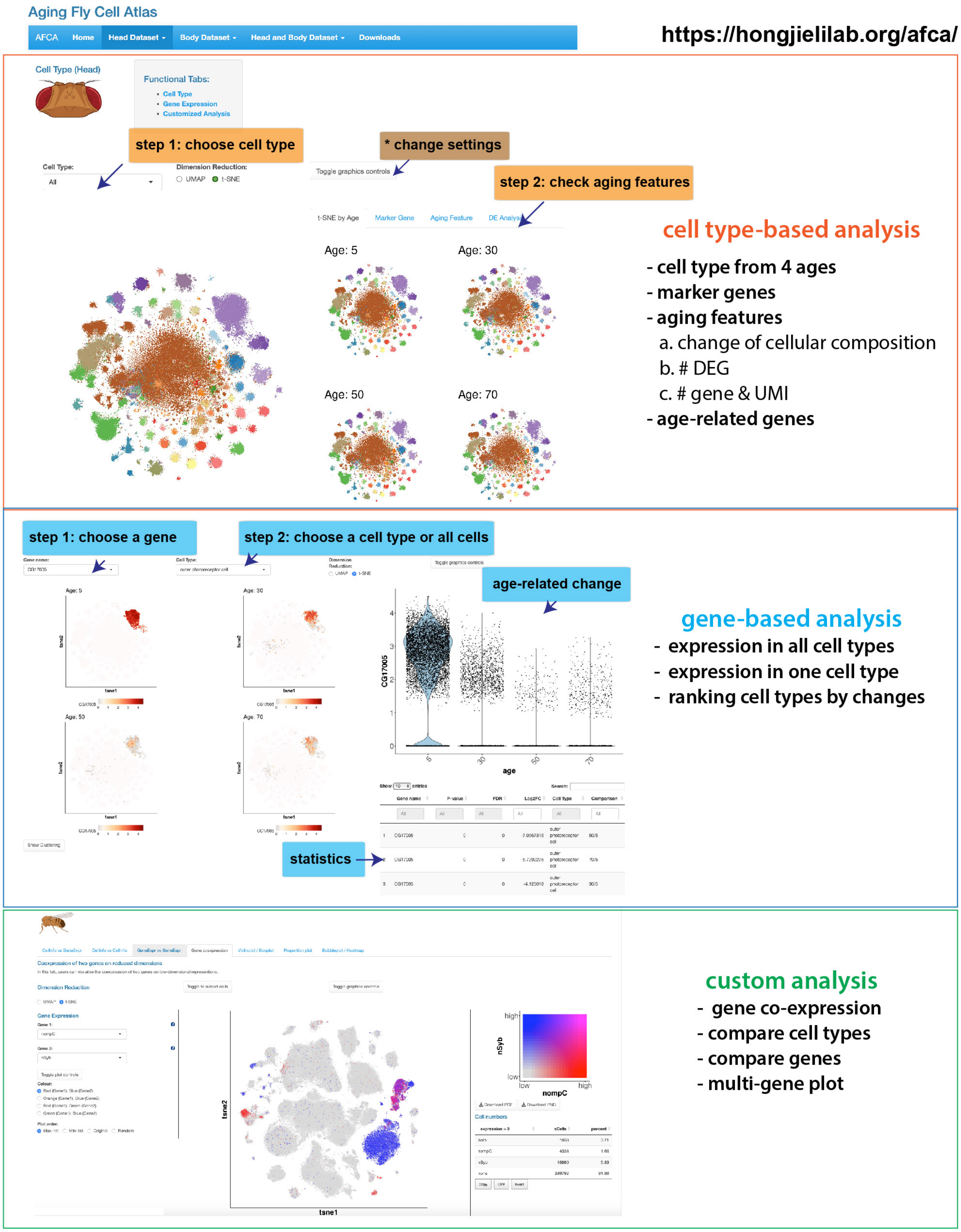
Aging Fly Cell Atlas
AFCA
Home
Head Dataset
Body Dataset
Head and Body Dataset
Downloads
https://hongjielilab.org/afca/
Cell Type (Head)
Functional Tabs:
Cell Type
Gene Expression
Customized Analysis
step 1: choose cell type
* change settings
step 2: check aging features
Cell Type:
All
Dimension Reduction:
UMAP
t-SNE
Toggle graphics controls
t-SNE by Age
Marker Gene
Aging Feature
Age: 5
Age: 30
Age: 50
Age: 70
cell type-based analysis
- cell type from 4 ages
- marker genes
- aging features
a. change of cellular composition
b. # DEG
c. # gene & UMI
- age-related genes
step 1: choose a gene
step 2: choose a cell type or all cells
age-related change
CG17005
age
statistics
gene-based analysis
- expression in all cell types
- expression in one cell type
- ranking cell types by changes
Coexpression of two genes on reduced dimensions
Dimension Reduction
Gene Expression
tsne1
tsne2
nSyb
nompC
high
low
Cell numbers
custom analysis
- gene co-expression
- compare cell types
- compare genes
- multi-gene plot

**Fig. S2. AFCA web portal for surveying the age-related changes.**
AFCA web portal provides cell type-based, gene-based, and customized analysis.

### Figure S3

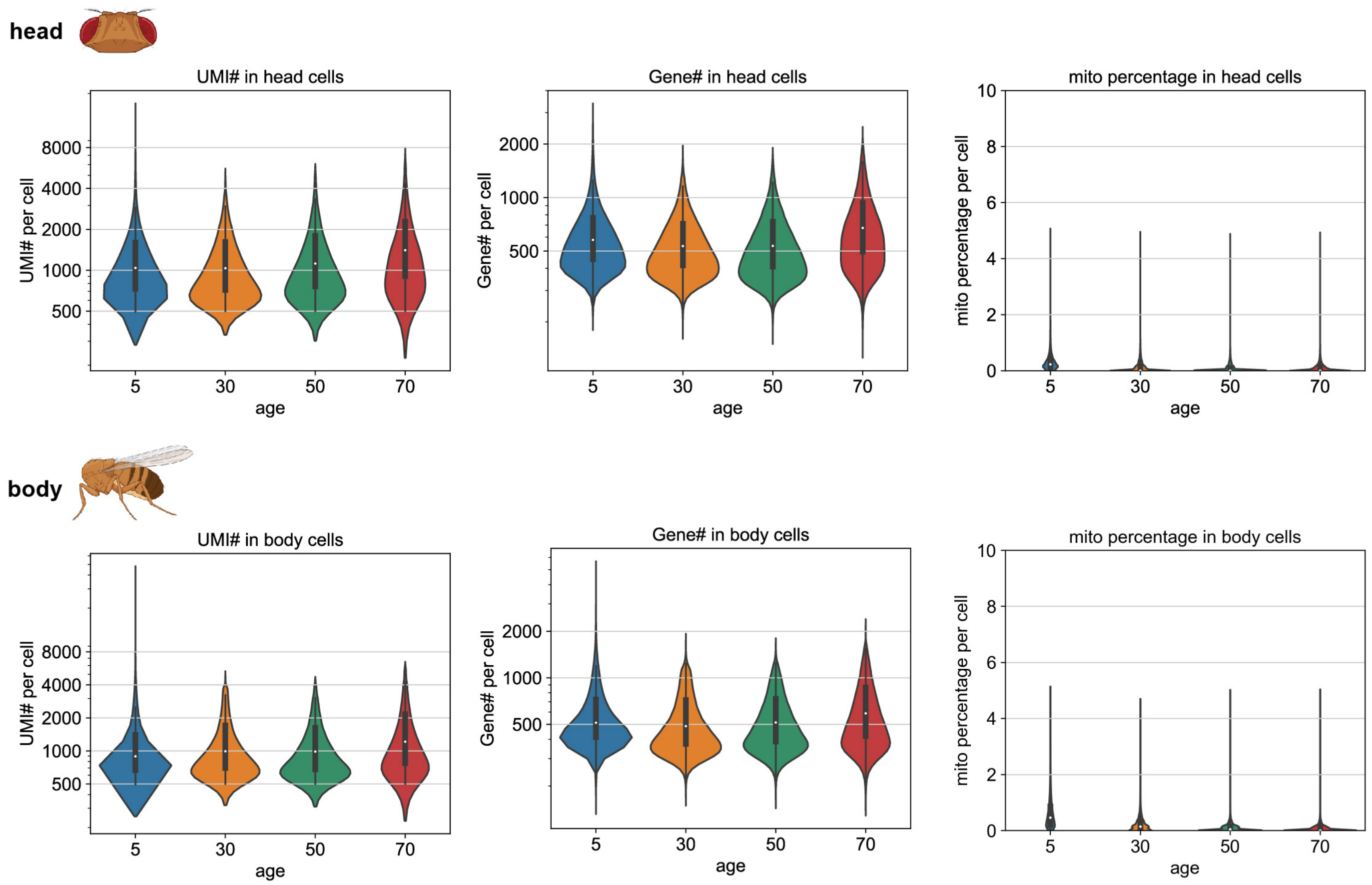
head
UMI# in head cells
UMI# per cell
8000
4000
2000
1000
500
5
30
50
70
age
Gene# in head cells
Gene# per cell
2000
1000
500
mito percentage in head cells
mito percentage per cell
10
8
6
4
2
0
body
UMI# in body cells
Gene# in body cells
mito percentage in body cells

**Fig. S3. General quality of snRNA-seq results.**
UMI number, expressed gene number, and mitochondria transcript ratio from the AFCA data.

### Figure S4

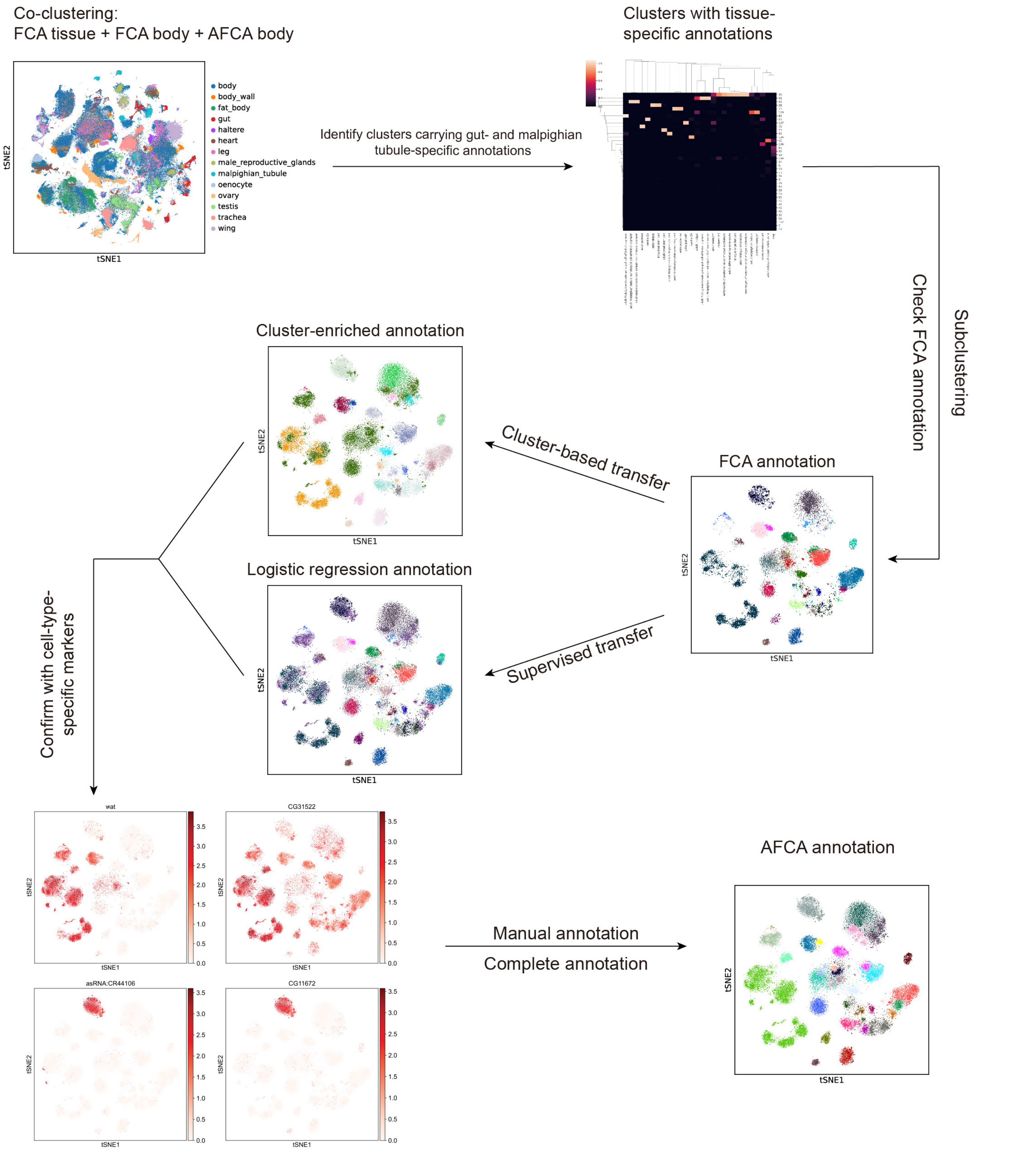

Co-clustering:
FCA tissue + FCA body + AFCA body
body
body_wall
fat_body
gut
haltere
heart
leg
male_reproductive_glands
malpighian_tubule
oenocyte
ovary
testis
trachea
wing
tSNE1
tSNE2
Identify clusters carrying gut- and malpighian tubule-specific annotations
Clusters with tissue-specific annotations
Check FCA annotation
Subclustering
FCA annotation
Cluster-based transfer
Supervised transfer
Cluster-enriched annotation
Logistic regression annotation
Confirm with cell-type-specific markers
wat
CG31522
asRNA:CR44106
CG11672
Manual annotation
Complete annotation
AFCA annotation

**Fig. S4. Illustration of transferring the FCA annotation to the AFCA cells.**

Flowchart of transferring the annotations from FCA to AFCA. FCA and AFCA data are first co-clustered. Co-clustered data can be further extracted for tissue-specific cell types, e.g. gut- and malpighian tubule-specific cell types. After subclustering targeted cells, FCA annotations are transferred to AFCA cells using cluster-based or supervised machine-learning approaches. Results from two transfer methods are compared and ambiguous transfers are further checked for the expression of the corresponding marker genes. After confirming the expression of the marker genes, annotations are corrected and finalized.

### Figure S5

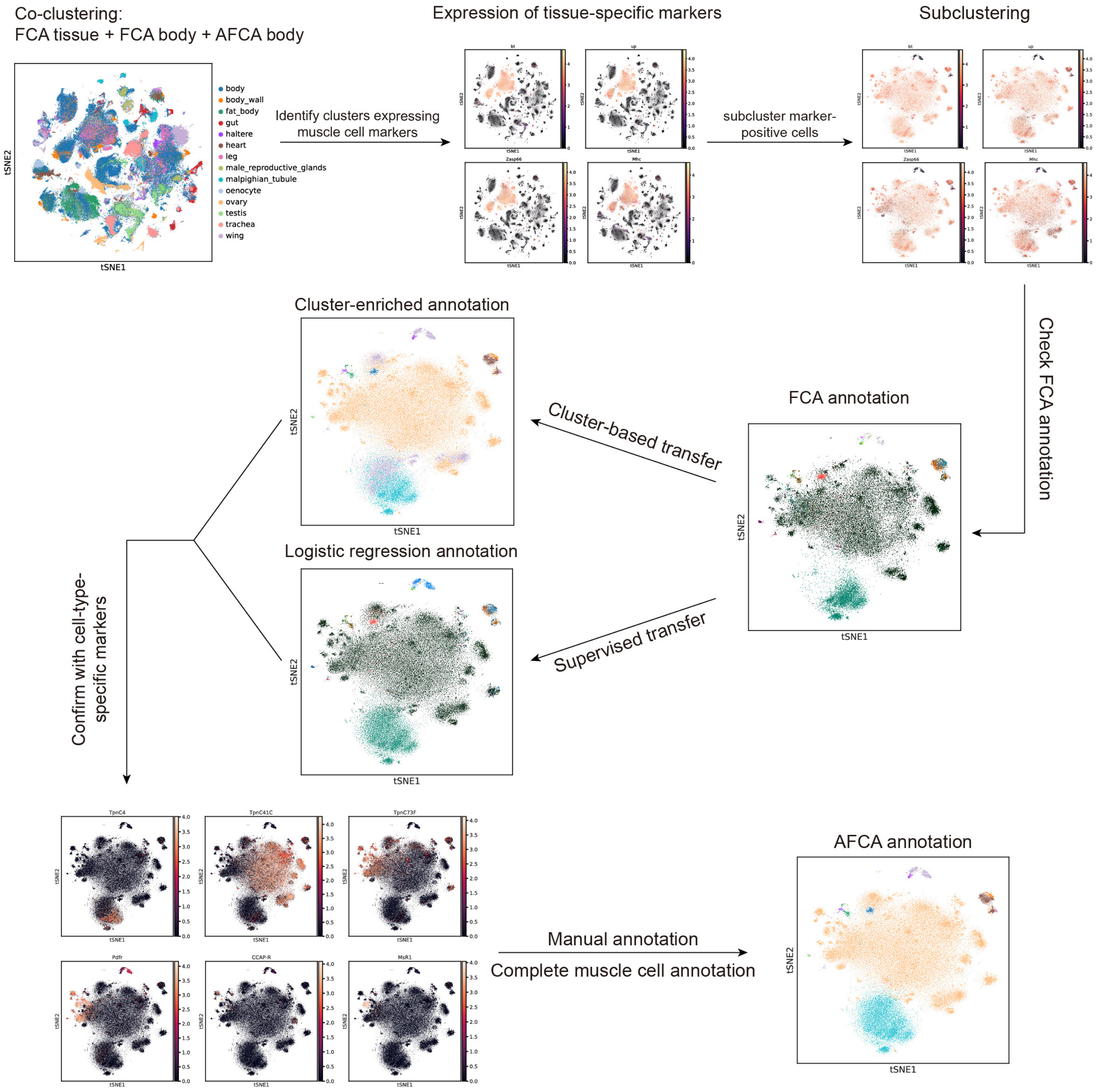
Co-clustering:
FCA tissue + FCA body + AFCA body
body
body_wall
fat_body
gut
haltere
heart
leg
male_reproductive_glands
malpighian_tubule
oenocyte
ovary
testis
trachea
wing
Identify clusters expressing muscle cell markers
Expression of tissue-specific markers
bt
up
Zasp66
Mhc
subcluster marker-positive cells
Subclustering
Check FCA annotation
FCA annotation
Cluster-based transfer
Supervised transfer
Cluster-enriched annotation
Logistic regression annotation
Confirm with cell-type-specific markers
TpnC4
TpnC41C
TpnC73F
Pdfr
CCAP-R
MsR1
Manual annotation
Complete muscle cell annotation
AFCA annotation
tSNE1
tSNE2

**Fig. S5. Illustration of transferring the FCA annotation to the AFCA muscle cells.** Flowchart of transferring the muscle-type annotations from FCA to AFCA. Some cell types, like muscle cells, can be found in multiple tissues and we consider first extracting those cells based on the expression of general markers of muscle cells. After subclustering the corresponding cells, all remaining analyses are similar to approaches used in **Fig. S4**.

### Figure S6

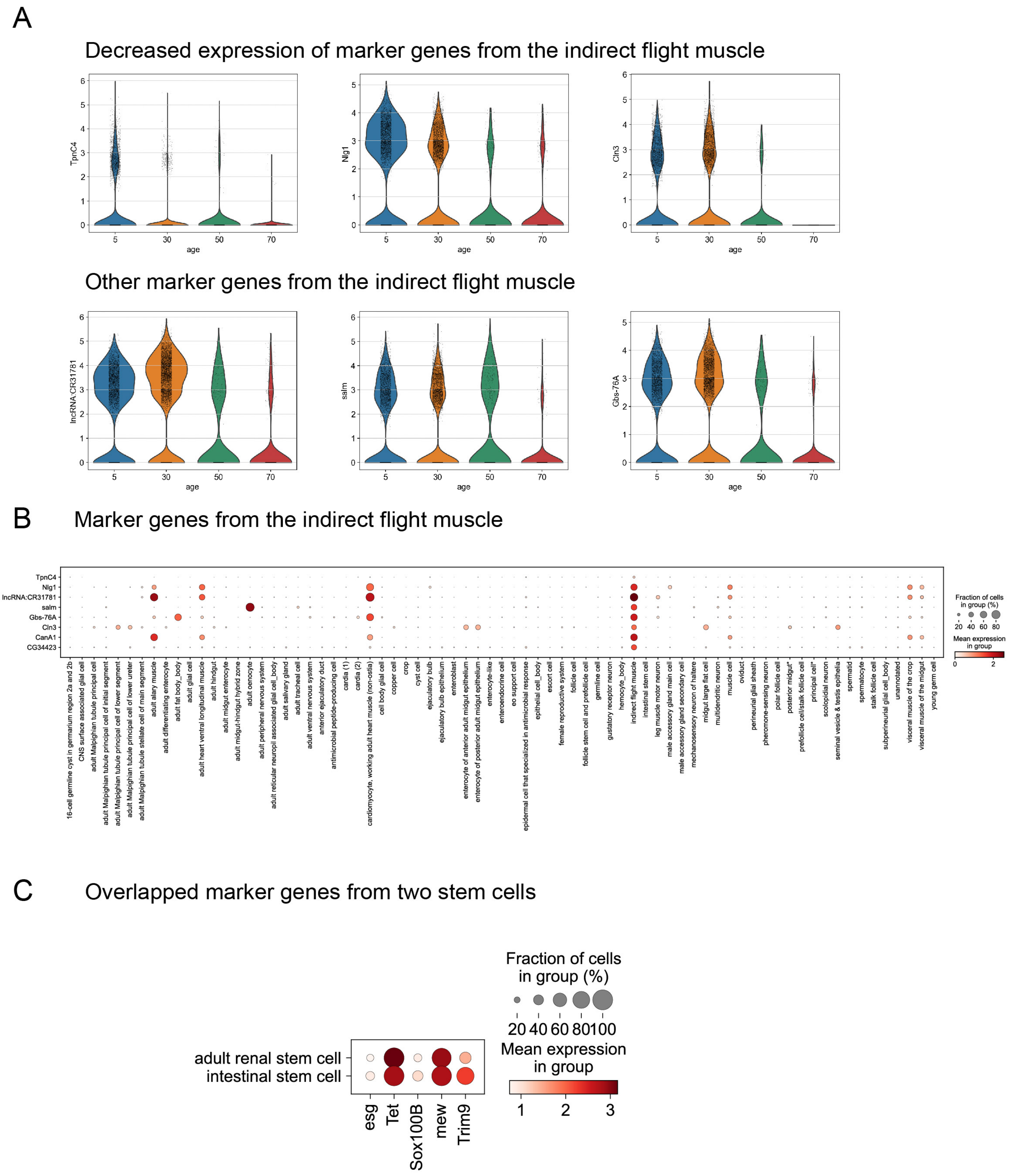
A
Decreased expression of marker genes from the indirect flight muscle
TpnC4
Nlg1
Cln3
age
5
30
50
70
Other marker genes from the indirect flight muscle
lncRNA:CR31781
salm
Gbs-76A
B
Marker genes from the indirect flight muscle
Fraction of cells in group (%)
Mean expression in group
C
Overlapped marker genes from two stem cells
adult renal stem cell
intestinal stem cell
esg
Tet
Sox100B
mew
Trim9
Fraction of cells in group (%)
20 40 60 80 100
Mean expression in group
1 2 3

**Fig. S6. Ambiguous annotations from the aging samples.**

A, B) Marker genes of the indirect flight muscle. A) Loss of marker genes in the indirect flight muscle. B) Markers specific for the indirect flight muscle. C) Overlapped marker genes from renal stem cells and intestinal stem cells.

### Figure S7

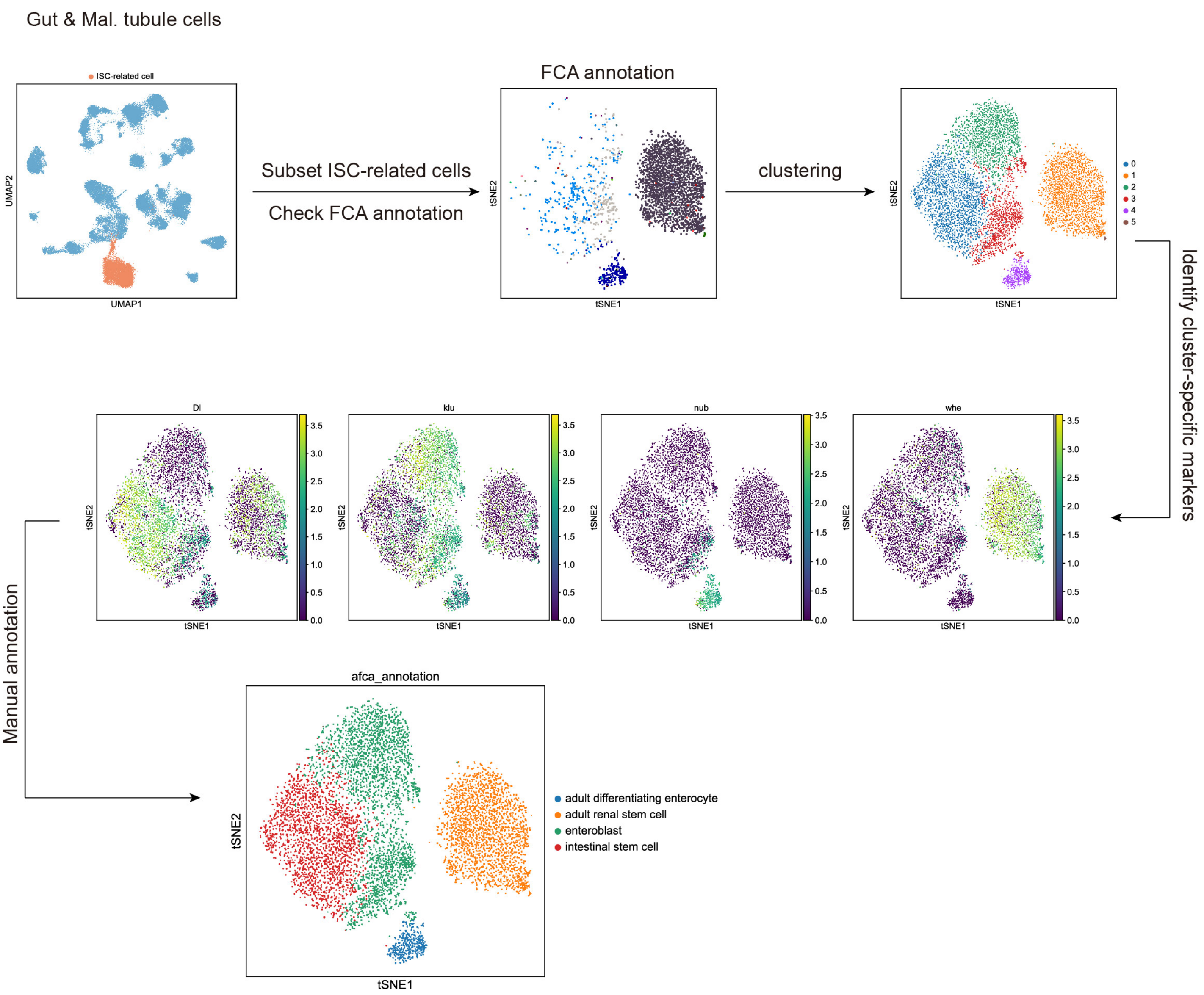
Gut & Mal. tubule cells
ISC-related cell
UMAP2
UMAP1
Subset ISC-related cells
Check FCA annotation
FCA annotation
tSNE2
tSNE1
clustering
0
1
2
3
4
5
Identify cluster-specific markers
Dl
klu
nub
whe
Manual annotation
afca_annotation
adult differentiating enterocyte
adult renal stem cell
enteroblast
intestinal stem cell

**Fig. S7. Illustration of manually annotating ISCs, EBs, and renal stem cells.**

Flowchart of manual annotation of the stem cell population. ISCs, EBs, and differentiating enterocyte cells are selected from the gut and malpighian tubule dataset shown in **Fig. S4**. Marker genes enriched in each cluster are identified and confirmed.

### Figure S8

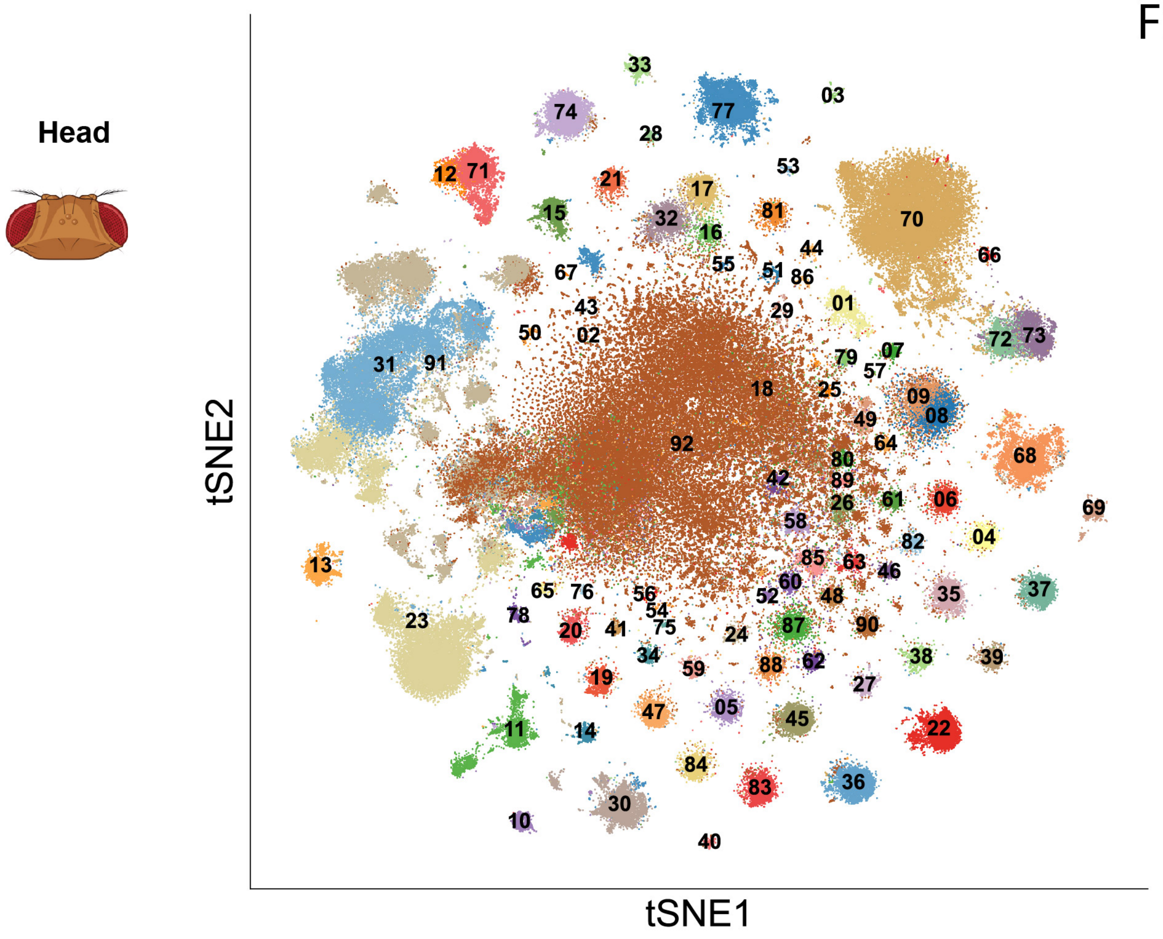


- 01.Johnston organ neuron
- 02.Kenyon cell
- 03.Poxn neuron
- 04.T neuron T2
- 05.T neuron T2a
- 06.T neuron T3
- 07.T neuron T4/T5
- 08.T neuron T4/T5a-b
- 09.T neuron T4/T5c-d
- 10.adult brain cell body glial cell
- 11.adult brain perineurial glial cell
- 12.adult fat body_head
- 13.adult lamina epithelial/marginal glial cell
- 14.adult optic chiasma glial cell
- 15.adult reticular neuropil associated glial cell_head
- 16.alpha'/beta' Kenyon cell
- 17.alpha/beta Kenyon cell
- 18.antennal lobe projection neuron
- 19.auditory sensory neuron
- 20.centrifugal neuron C2
- 21.centrifugal neuron C3
- 22.columnar neuron T1
- 23.cone cell
- 24.distal medullary amacrine neuron Dm10
- 25.distal medullary amacrine neuron Dm12
- 26.distal medullary amacrine neuron Dm3
- 27.distal medullary amacrine neuron Dm8
- 28.distal medullary amacrine neuron Dm9
- 29.dopaminergic PAM neuron
- 30.ensheathing glial cell
- 31.epithelial cell_head
- 32.gamma Kenyon cell
- 33.hemocyte_head
- 34.lamina intrinsic amacrine neuron Lai
- 35.lamina monopolar neuron L1
- 36.lamina monopolar neuron L2
- 37.lamina monopolar neuron L3
- 38.lamina monopolar neuron L4
- 39.lamina monopolar neuron L5
- 40.lamina wide-field 1 neuron
- 41.lamina wide-field 2 neuron
- 42.lobula columnar neuron LC10
- 43.lobula columnar neuron LC12
- 44.lobula columnar neuron LC17
- 45.medullary intrinsic neuron Mi1
- 46.medullary intrinsic neuron Mi15
- 47.medullary intrinsic neuron Mi4
- 48.medullary intrinsic neuron Mi9
- 49.neuron 1
- 50.neuron 10
- 51.neuron 11
- 52.neuron 12
- 53.neuron 13
- 54.neuron 14
- 55.neuron 15
- 56.neuron 16
- 57.neuron 17
- 58.neuron 2
- 59.neuron 3
- 60.neuron 4
- 61.neuron 5
- 62.neuron 6
- 63.neuron 7
- 64.neuron 8
- 65.neuron 9
- 66.ocellus retinula cell
- 67.octopaminergic/tyraminergic neuron
- 68.olfactory receptor neuron
- 69.optic-lobe-associated cortex glial cell
- 70.outer photoreceptor cell
- 71.pericerebral adult fat mass
- 72.photoreceptor cell R7
- 73.photoreceptor cell R8
- 74.pigment cell
- 75.proximal medullary amacrine neuron Pm2
- 76.proximal medullary amacrine neuron Pm4
- 77.skeletal muscle of head
- 78.subperineurial glial cell_head
- 79.transmedullary Y neuron TmY14
- 80.transmedullary Y neuron TmY4
- 81.transmedullary Y neuron TmY5a
- 82.transmedullary Y neuron TmY8
- 83.transmedullary neuron Tm1
- 84.transmedullary neuron Tm2
- 85.transmedullary neuron Tm20
- 86.transmedullary neuron Tm29
- 87.transmedullary neuron Tm3a
- 88.transmedullary neuron Tm4
- 89.transmedullary neuron Tm5c
- 90.transmedullary neuron Tm9
- 91.unannotated
- 92.uncharacterized CNS neuron

**Fig. S8. Detailed cell types annotated in the head sample shown by tSNE.**
91 head cell types are shown in the tSNE plot.

### Figure S9

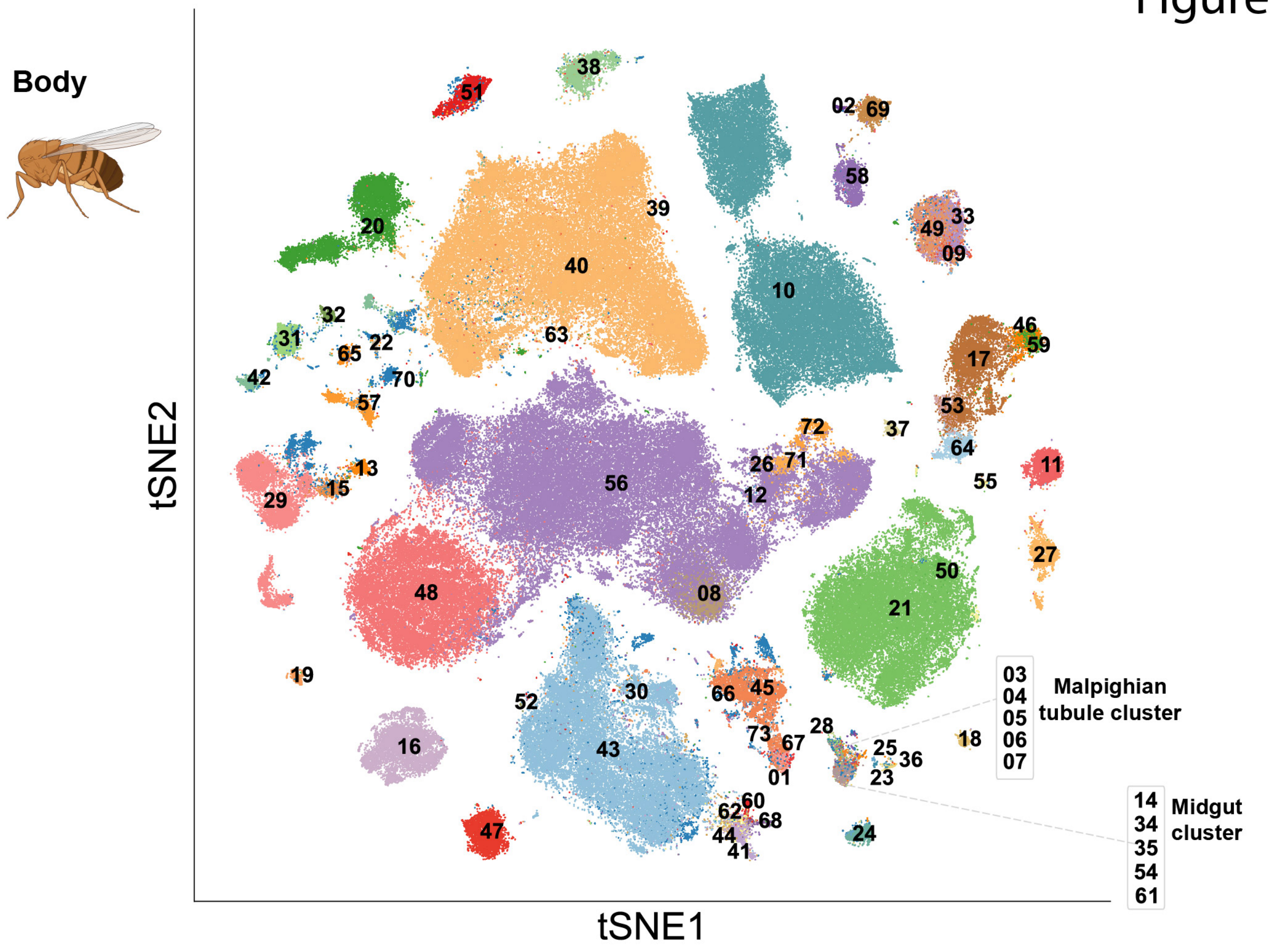


- 01.16-cell germline cyst in germarium region 2a and 2b
- 02.CNS surface associated glial cell
- 03.adult Malpighian tubule principal cell
- 04.adult Malpighian tubule principal cell of initial segment
- 05.adult Malpighian tubule principal cell of lower segment
- 06.adult Malpighian tubule principal cell of lower ureter
- 07.adult Malpighian tubule stellate cell of main segment
- 08.adult alary muscle
- 09.adult differentiating enterocyte
- 10.adult fat body_body
- 11.adult glial cell
- 12.adult heart ventral longitudinal muscle
- 13.adult hindgut
- 14.adult midgut enterocyte
- 15.adult midgut-hindgut hybrid zone
- 16.adult oenocyte
- 17.adult peripheral nervous system
- 18.adult reticular neuropil associated glial cell_body
- 19.adult salivary gland
- 20.adult tracheal cell
- 21.adult ventral nervous system
- 22.anterior ejaculatory duct
- 23.antimicrobial peptide-producing cell
- 24.cardia (1)
- 25.cardia (2)
- 26.cardiomyocyte, working adult heart muscle (non-ostia)
- 27.cell body glial cell
- 28.copper cell
- 29.crop
- 30.cyst cell
- 31.ejaculatory bulb
- 32.ejaculatory bulb epithelium
- 33.enteroblast
- 34.enterocyte of anterior adult midgut epithelium
- 35.enterocyte of posterior adult midgut epithelium
- 36.enterocyte-like
- 37.enteroendocrine cell
- 38.eo support cell
- 39.epidermal cell that specialized in antimicrobial response
- 40.epithelial cell_body
- 41.escort cell
- 42.female reproductive system
- 43.follicle cell
- 44.follicle stem cell and prefollicle cell
- 45.germline cell
- 46.gustatory receptor neuron
- 47.hemocyte_body
- 48.indirect flight muscle
- 49.intestinal stem cell
- 50.leg muscle motor neuron
- 51.male accessory gland main cell
- 52.male accessory gland secondary cell
- 53.mechanosensory neuron of haltere
- 54.midgut large flat cell
- 55.multidendritic neuron
- 56.muscle cell
- 57.oviduct
- 58.perineurial glial sheath
- 59.pheromone-sensing neuron
- 60.polar follicle cell
- 61.posterior midgut*
- 62.prefollicle cell/stalk follicle cell
- 63.principal cell*
- 64.scolopidial neuron
- 65.seminal vesicle & testis epithelia
- 66.spermatid
- 67.spermatocyte
- 68.stalk follicle cell
- 69.subperineurial glial cell_body
- 70.unannotated
- 71.visceral muscle of the crop
- 72.visceral muscle of the midgut
- 73.young germ cell

**Fig. S9. Detailed cell types annotated in the body sample shown by tSNE.**
72 body cell types are shown in the tSNE plot.

### Figure S10

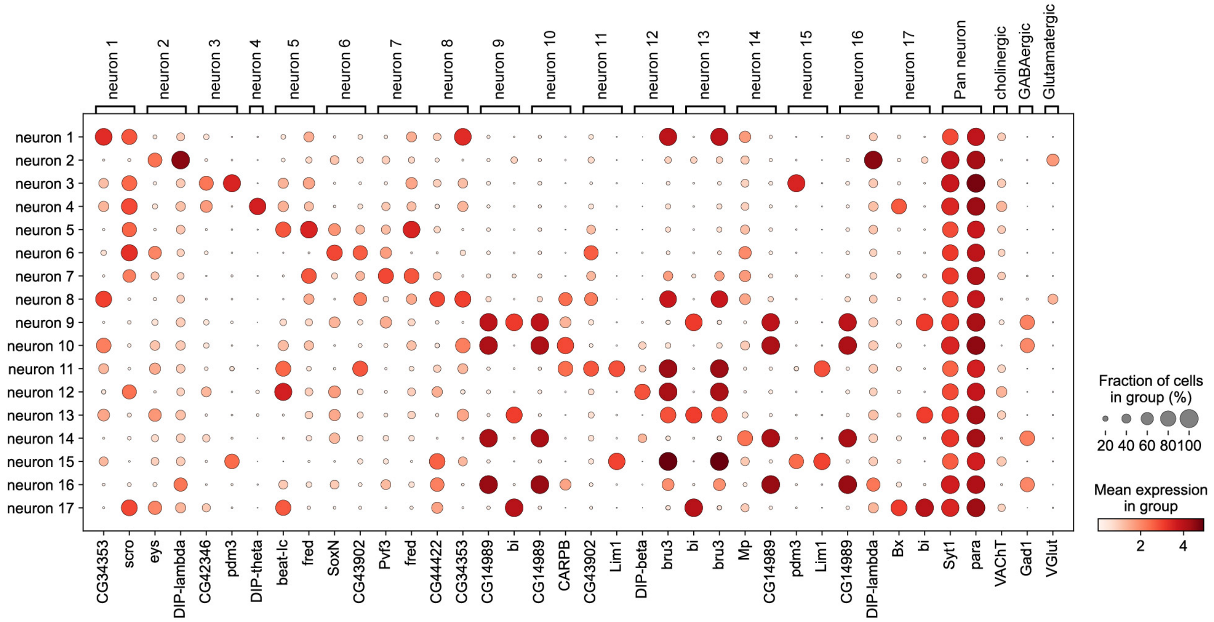

neuron 1
neuron 2
neuron 3
neuron 4
neuron 5
neuron 6
neuron 7
neuron 8
neuron 9
neuron 10
neuron 11
neuron 12
neuron 13
neuron 14
neuron 15
neuron 16
neuron 17
Pan neuron
cholinergic
GABAergic
Glutamatergic
CG34353
scro
eys
DIP-lambda
CG42346
pdm3
DIP-theta
beat-Ic
fred
SoxN
CG43902
Pvf3
fred
CG44422
CG34353
CG14989
bi
CG14989
CARPB
CG43902
Lim1
DIP-beta
bru3
bi
bru3
Mp
CG14989
pdm3
Lim1
CG14989
DIP-lambda
Bx
bi
Syt1
para
VAChT
Gad1
VGlut
Fraction of cells in group (%)
20 40 60 80 100
Mean expression in group
2 4

**Fig. S10. Marker genes identified from the newly identified neuron clusters.**

Markers of 17 newly identified neuron types are shown. Expression of general neuronal markers, Syt1, and para, indicates neuron specificity. Expression of VAChT, Gad1, and VGlut indicates they are cholinergic, GABAergic, or Glutamatergic neurons.

### Figure S11

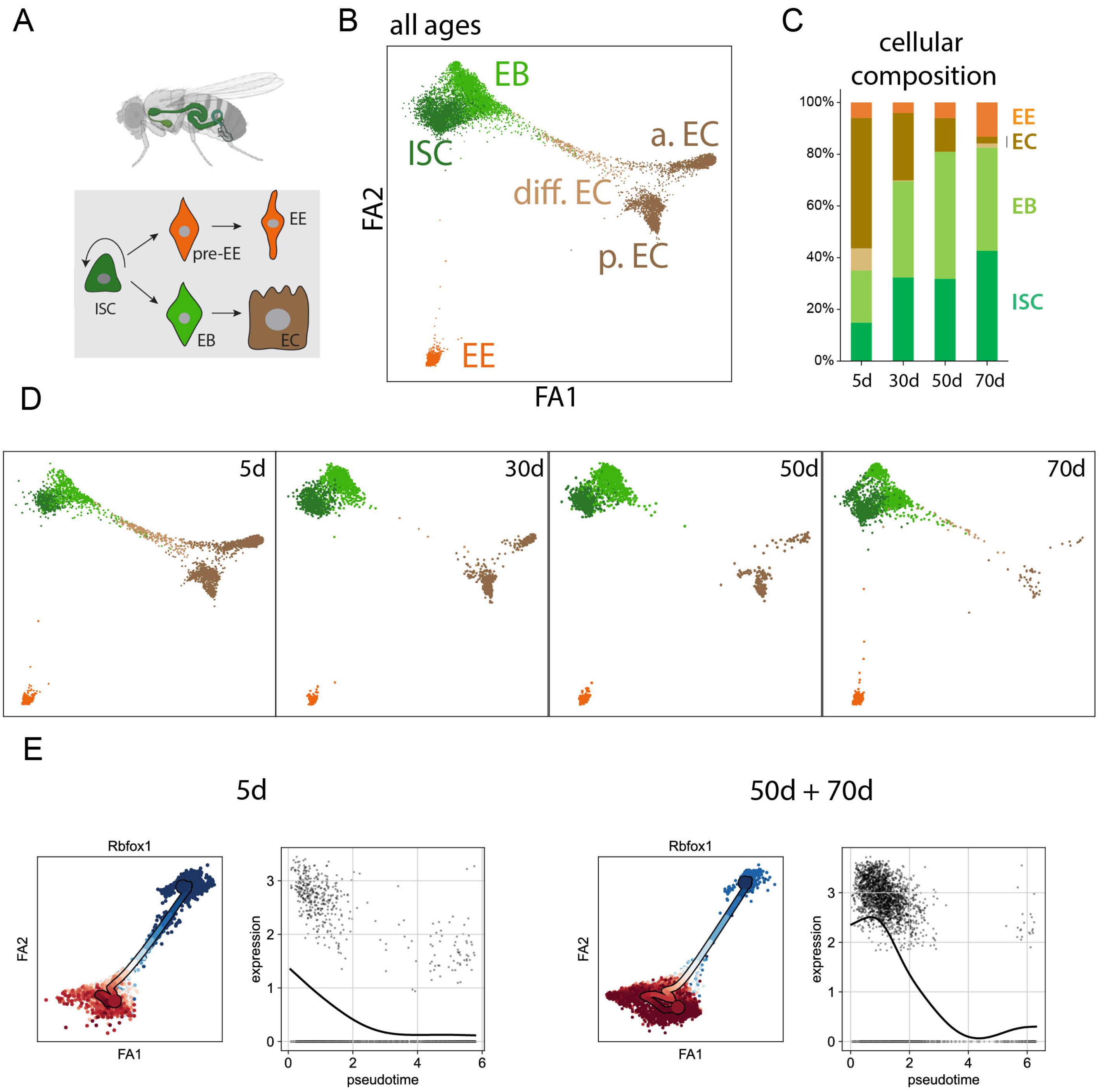

A
EE
pre-EE
ISC
EB
EC
B
all ages
EB
ISC
a. EC
diff. EC
p. EC
EE
FA2
FA1
C
cellular composition
100%
80%
60%
40%
20%
0%
5d
30d
50d
70d
EE
EC
EB
ISC
D
5d
30d
50d
70d
E
5d
50d + 70d
Rbfox1
FA2
FA1
expression
pseudotime

**Fig. S11 Pseudotime inference and cellular composition of ISC and ISC-differentiated cell types.**

A) Illustration of ISC differentiation process. B) Cell types shown in different colors. C) Changes of cellular composition in the ISC lineage. D) Age-specific profile of different cell types. E) Rbfox1 gene shows different patterns along the young and old ISC lineage. ISC, intestinal stem cell; EB, enteroblast; EC, enterocyte; EE, enteroendocrine cell; a. EC, anterior EC; p. EC, posterior EC; diff. EC, differentiating EC.

### Figure S12

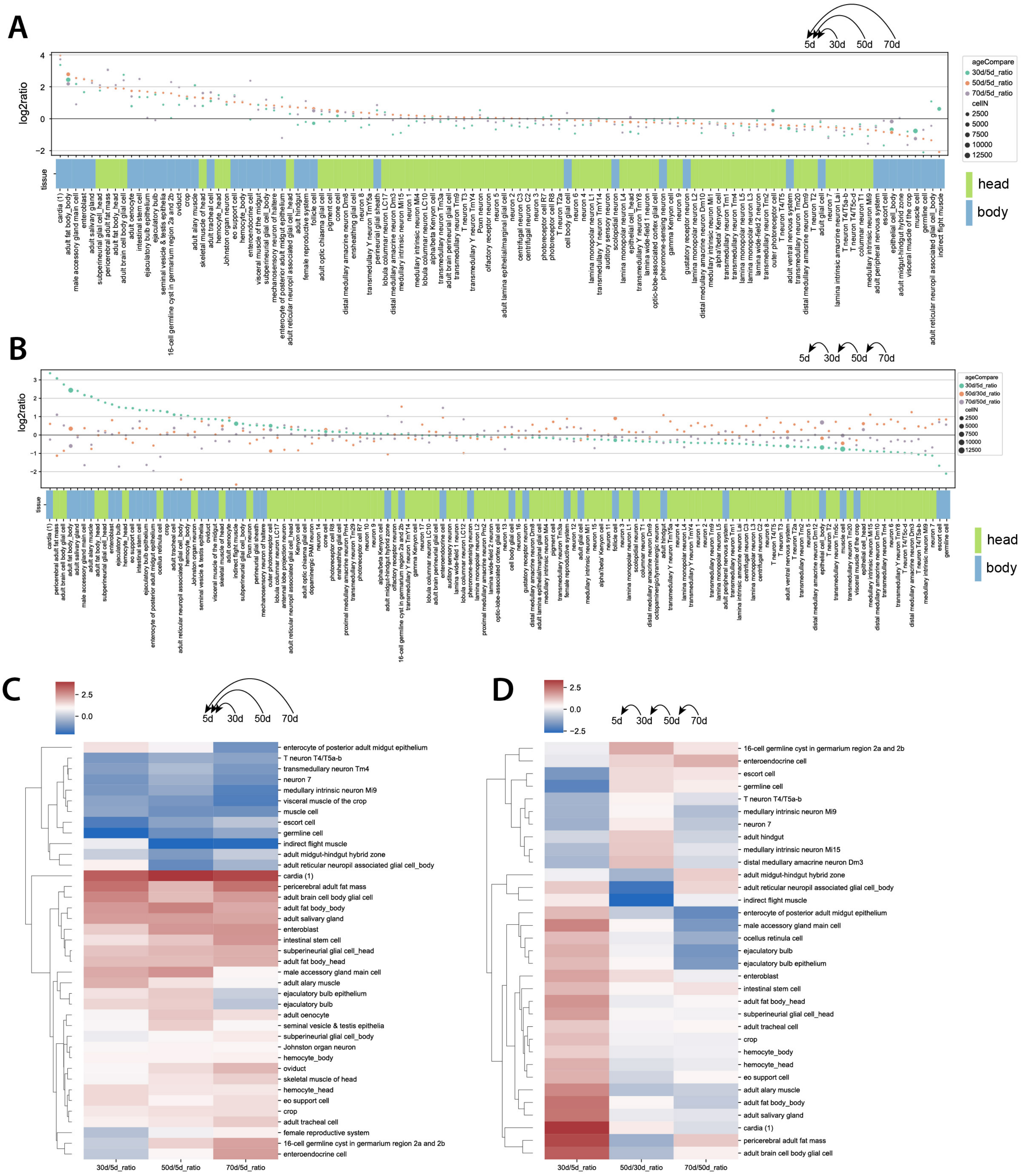
A
5d 30d 50d 70d
log2ratio
tissue
ageCompare
30d/5d_ratio
50d/5d_ratio
70d/5d_ratio
cellN
2500
5000
7500
10000
12500
head
body
B
5d 30d 50d 70d
log2ratio
tissue
ageCompare
30d/5d_ratio
50d/30d_ratio
70d/50d_ratio
cellN
2500
5000
7500
10000
12500
head
body
C
2.5
0.0
5d 30d 50d 70d
enterocyte of posterior adult midgut epithelium
T neuron T4/T5a-b
transmedullary neuron Tm4
neuron 7
medullary intrinsic neuron Mi9
visceral muscle of the crop
muscle cell
escort cell
germline cell
indirect flight muscle
adult midgut-hindgut hybrid zone
adult reticular neuropil associated glial cell_body
cardia (1)
pericerebral adult fat mass
adult brain cell body glial cell
adult fat body_body
adult salivary gland
enteroblast
intestinal stem cell
subperineurial glial cell_head
adult fat body_head
male accessory gland main cell
adult alary muscle
ejaculatory bulb epithelium
ejaculatory bulb
adult oenocyte
seminal vesicle & testis epithelia
subperineurial glial cell_body
Johnston organ neuron
hemocyte_body
oviduct
skeletal muscle of head
hemocyte_head
eo support cell
crop
adult tracheal cell
female reproductive system
16-cell germline cyst in germarium region 2a and 2b
enteroendocrine cell
30d/5d_ratio
50d/5d_ratio
70d/5d_ratio
D
2.5
0.0
−2.5
5d 30d 50d 70d
16-cell germline cyst in germarium region 2a and 2b
enteroendocrine cell
escort cell
germline cell
T neuron T4/T5a-b
medullary intrinsic neuron Mi9
neuron 7
adult hindgut
medullary intrinsic neuron Mi15
distal medullary amacrine neuron Dm3
adult midgut-hindgut hybrid zone
adult reticular neuropil associated glial cell_body
indirect flight muscle
enterocyte of posterior adult midgut epithelium
male accessory gland main cell
ocellus retinula cell
ejaculatory bulb
ejaculatory bulb epithelium
enteroblast
intestinal stem cell
adult fat body_head
subperineurial glial cell_head
adult tracheal cell
crop
hemocyte_body
hemocyte_head
eo support cell
adult alary muscle
adult fat body_body
adult salivary gland
cardia (1)
pericerebral adult fat mass
adult brain cell body glial cell
30d/5d_ratio
50d/30d_ratio
70d/50d_ratio

**Fig. S12 Changes of cell-type composition during aging.**

A) Changes in cellular composition by comparing the nuclear ratio of the old sample (30d, 50d, and 70d) to the ratio of the young one (5d). B) Changes of cellular composition by comparing the nuclear ratio between two consecutive ages. C) Cell types with more than 2-fold changes of cellular composition by comparing the old samples to the young ones. D) Cell types with more than 2-fold changes of cellular composition by comparing the sample to the previous age.

### Figure S13

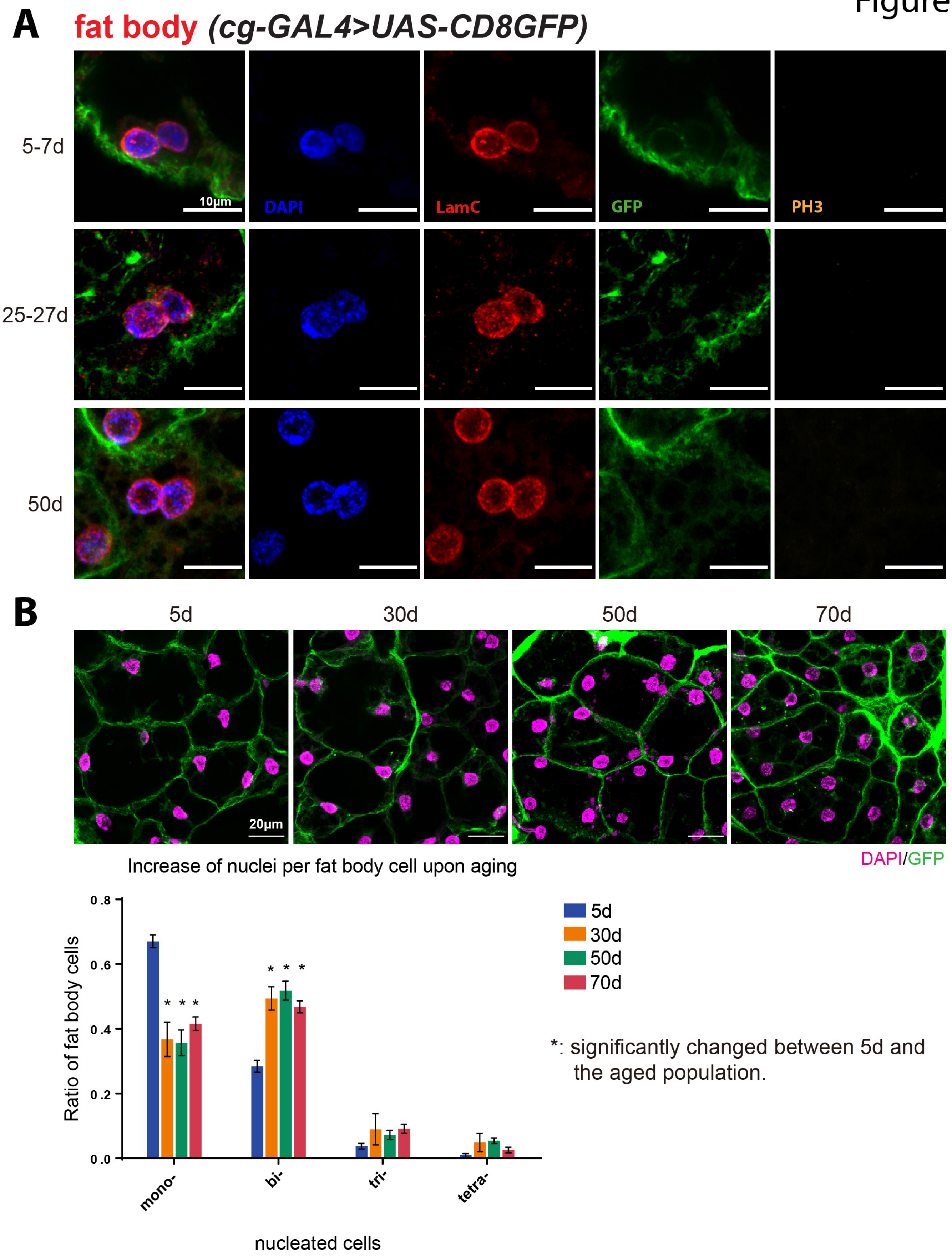
A
fat body (cg-GAL4>UAS-CD8GFP)
5-7d
25-27d
50d
10μm
DAPI
LamC
GFP
PH3
B
5d
30d
50d
70d
20μm
DAPI/GFP
Increase of nuclei per fat body cell upon aging
Ratio of fat body cells
0.8
0.6
0.4
0.2
0.0
mono-
bi-
tri-
tetra-
nucleated cells
5d
30d
50d
70d
*: significantly changed between 5d and the aged population.

**Fig. S13. Examples of dividing nuclei without the mitosis marker.**

A) Dividing nuclei from flies with different ages showing the absence of the mitosis marker (pH3). Fat body nuclei are collected from different ages, including 5-7d, 25-27d, and 50d. Dividing nuclei are stained by DAPI and LamC. The fat body membranes are labeled by *cg-GAL4 > UAS-CD8GFP*. B) The multinucleation of fat body cells in aged flies The representative images of fat bodies of different ages are shown. The multinucleation of the fat body was examined for at least 324 cells (5d:359, 30d:324, 50d:369, and 70d:494 cells) in 7-10 animals of each age. The two-way ANOVA and Tukey's multiple comparisons test were performed. The asterisks show the $P<0.0001$.

Figure S14

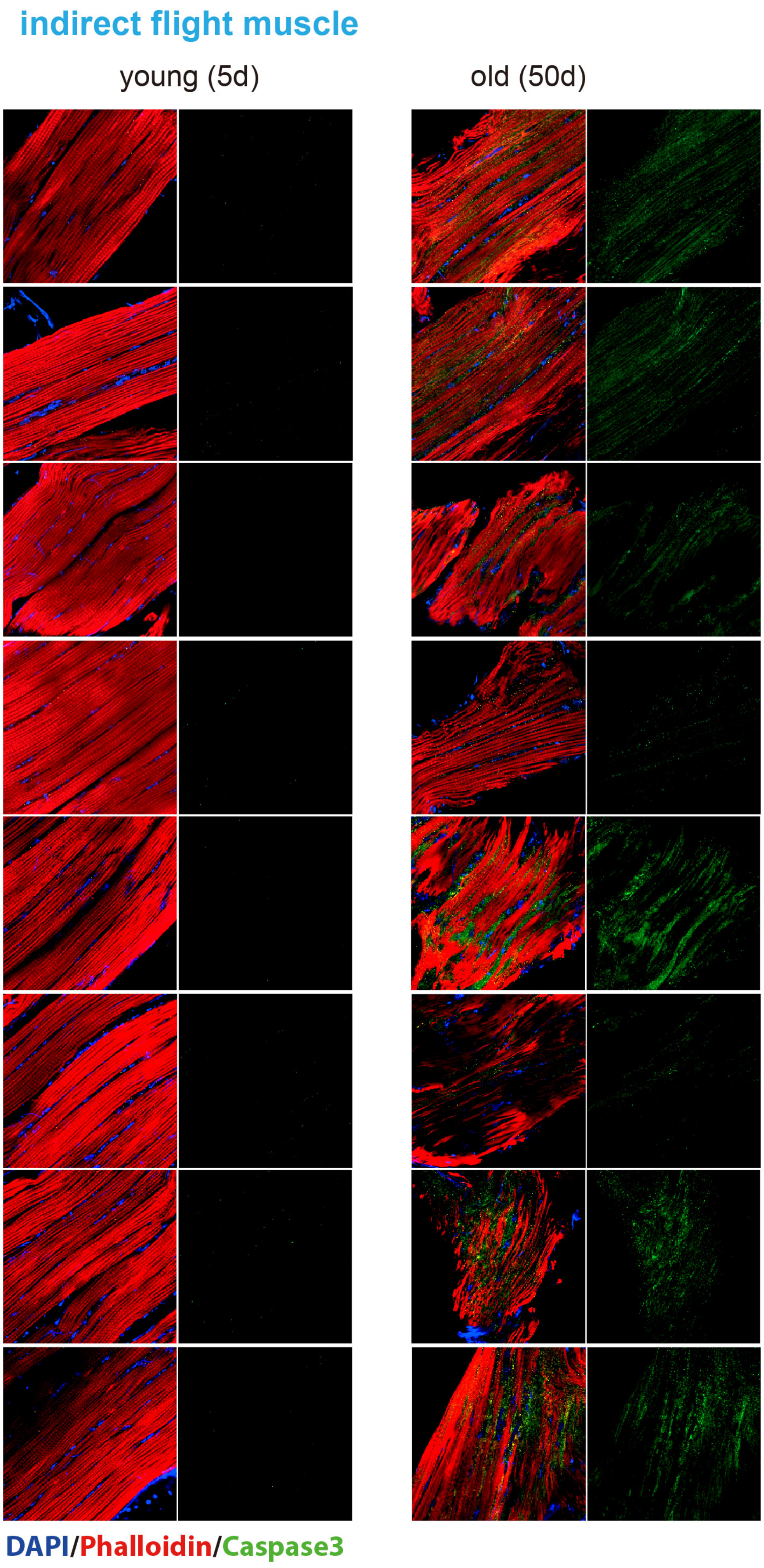
indirect flight muscle
young (5d)
old (50d)
DAPI/Phalloidin/Caspase3

**Fig. S14 Apoptosis signals form the aged indirect flight muscle.**

Examples of indirect flight muscle stained by the apoptosis marker in the young and old flies. The indirect flight muscles are stained for the apoptosis events using a cleaved-Caspase3 antibody. Nuclei are stained by DAPI. Actin filaments in the muscle cells are labeled by Phalloidin.

### Figure S15

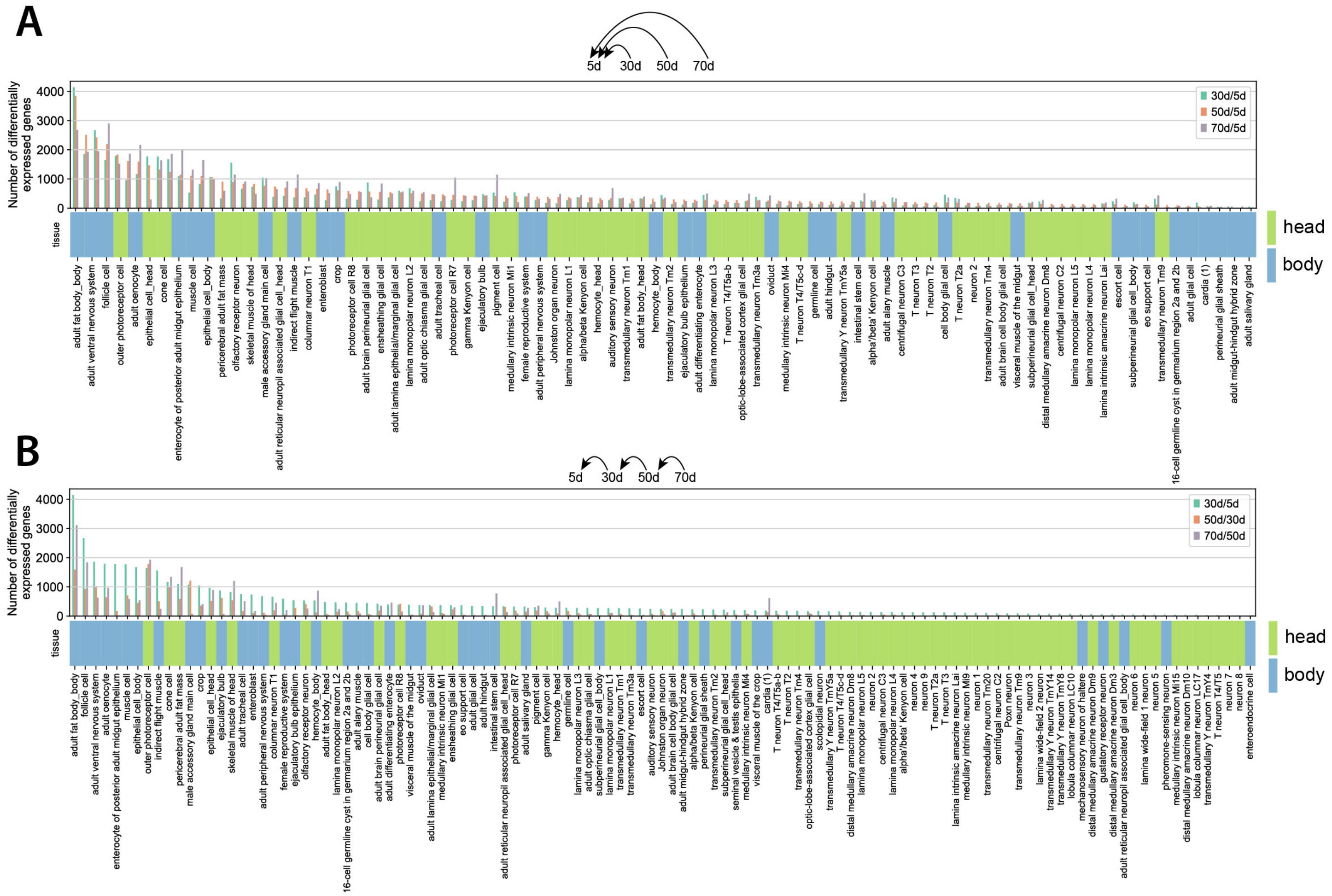

A
5d
30d
50d
70d
Number of differentially expressed genes
30d/5d
50d/5d
70d/5d
tissue
head
body
B
5d
30d
50d
70d
Number of differentially expressed genes
30d/5d
50d/30d
70d/50d
tissue
head
body

**Fig. S15 DEG numbers detected in each cell type.**
A) DEG numbers from different cell types. The aged groups (30d, 50d, and 70d) are compared with the young population (5d). B) Two consecutive ages are compared to identify DEGs in each cell type.

### Figure S16

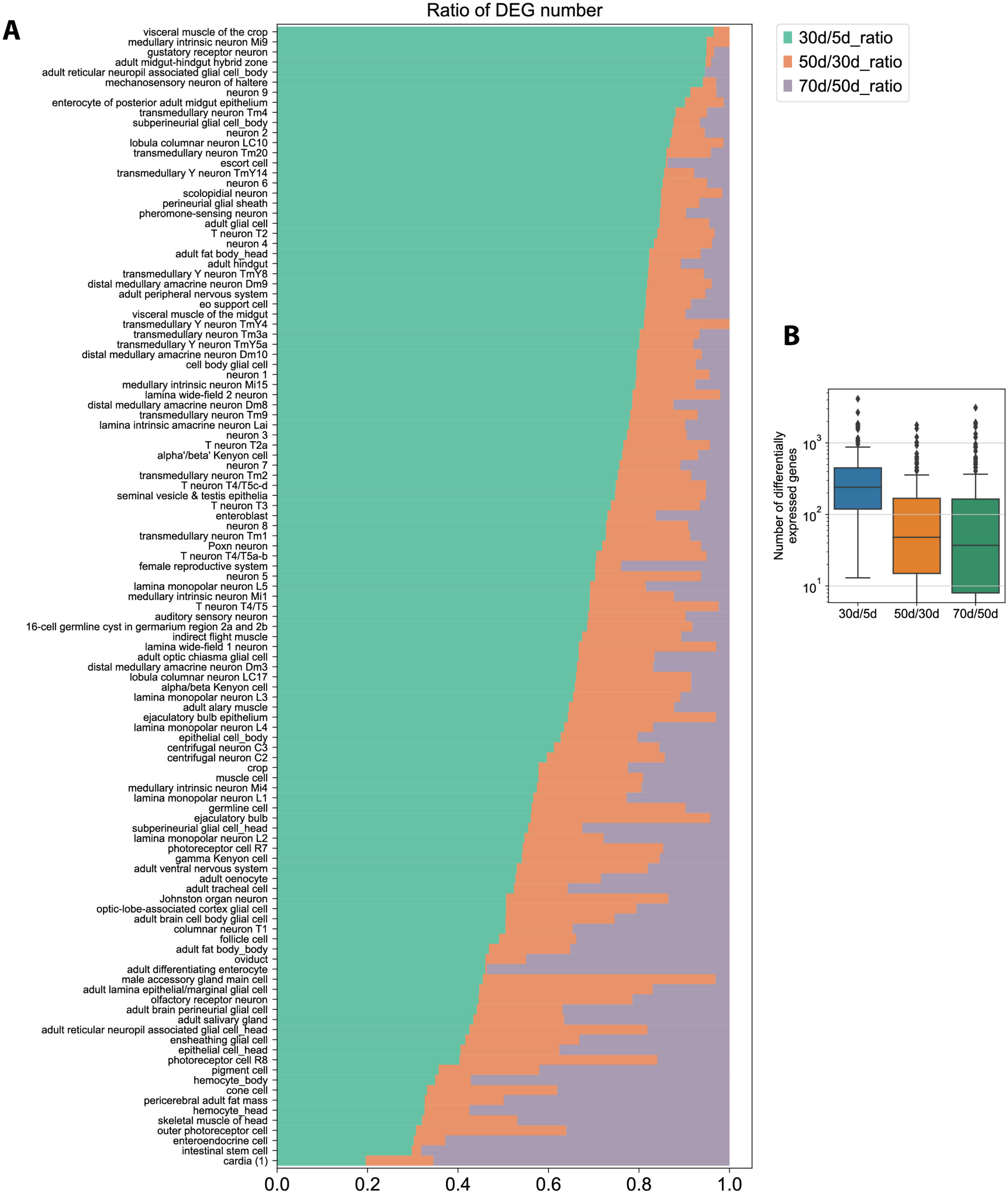
A
Ratio of DEG number
visceral muscle of the crop
medullary intrinsic neuron Mi9
gustatory receptor neuron
adult midgut-hindgut hybrid zone
adult reticular neuropil associated glial cell_body
mechanosensory neuron of haltere
neuron 9
enterocyte of posterior adult midgut epithelium
transmedullary neuron Tm4
subperineurial glial cell_body
neuron 2
lobula columnar neuron LC10
transmedullary neuron Tm20
escort cell
transmedullary Y neuron TmY14
neuron 6
scolopidial neuron
perineurial glial sheath
pheromone-sensing neuron
adult glial cell
T neuron T2
neuron 4
adult fat body_head
adult hindgut
transmedullary Y neuron TmY8
distal medullary amacrine neuron Dm9
adult peripheral nervous system
eo support cell
visceral muscle of the midgut
transmedullary Y neuron TmY4
transmedullary neuron Tm3a
transmedullary Y neuron TmY5a
distal medullary amacrine neuron Dm10
cell body glial cell
neuron 1
medullary intrinsic neuron Mi15
lamina wide-field 2 neuron
distal medullary amacrine neuron Dm8
transmedullary neuron Tm9
lamina intrinsic amacrine neuron Lai
neuron 3
T neuron T2a
alpha'/beta' Kenyon cell
neuron 7
transmedullary neuron Tm2
T neuron T4/T5c-d
seminal vesicle & testis epithelia
T neuron T3
enteroblast
neuron 8
transmedullary neuron Tm1
Poxn neuron
T neuron T4/T5a-b
female reproductive system
neuron 5
lamina monopolar neuron L5
medullary intrinsic neuron Mi1
T neuron T4/T5
auditory sensory neuron
16-cell germline cyst in germarium region 2a and 2b
indirect flight muscle
lamina wide-field 1 neuron
adult optic chiasma glial cell
distal medullary amacrine neuron Dm3
lobula columnar neuron LC17
alpha/beta Kenyon cell
lamina monopolar neuron L3
adult alary muscle
ejaculatory bulb epithelium
lamina monopolar neuron L4
epithelial cell_body
centrifugal neuron C3
centrifugal neuron C2
crop
muscle cell
medullary intrinsic neuron Mi4
lamina monopolar neuron L1
germline cell
ejaculatory bulb
subperineurial glial cell_head
lamina monopolar neuron L2
photoreceptor cell R7
gamma Kenyon cell
adult ventral nervous system
adult oenocyte
adult tracheal cell
Johnston organ neuron
optic-lobe-associated cortex glial cell
adult brain cell body glial cell
columnar neuron T1
follicle cell
adult fat body_body
oviduct
adult differentiating enterocyte
male accessory gland main cell
adult lamina epithelial/marginal glial cell
olfactory receptor neuron
adult brain perineurial glial cell
adult salivary gland
adult reticular neuropil associated glial cell_head
ensheathing glial cell
epithelial cell_head
photoreceptor cell R8
pigment cell
hemocyte_body
cone cell
pericerebral adult fat mass
hemocyte_head
skeletal muscle of head
outer photoreceptor cell
enteroendocrine cell
intestinal stem cell
cardia (1)
0.0
0.2
0.4
0.6
0.8
1.0
30d/5d_ratio
50d/30d_ratio
70d/50d_ratio
B
Number of differentially expressed genes
$10^3$
$10^2$
$10^1$
30d/5d
50d/30d
70d/50d

**Fig. S16 Detailed ratio of DEG number across different age periods.**
A) The ratios of DEG numbers in different cell types. Each age period is shown in a different color.
B) Number of differentially expressed genes detected in the different age periods.

### Figure S17

**A** **DEG comparsions between AFCA and Davie et al., 2018**

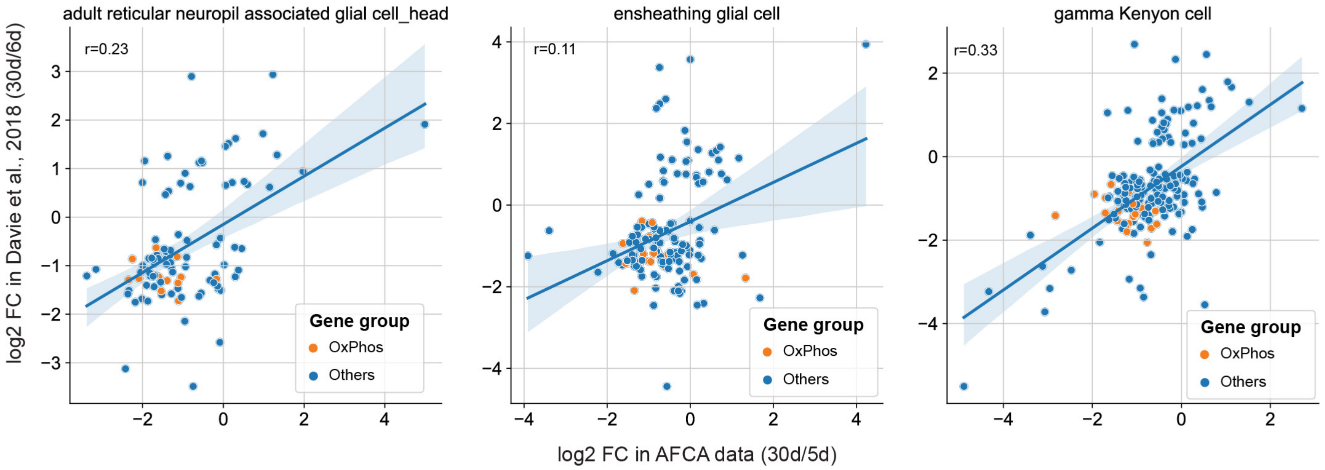


**B** **ensheathing glial cell**

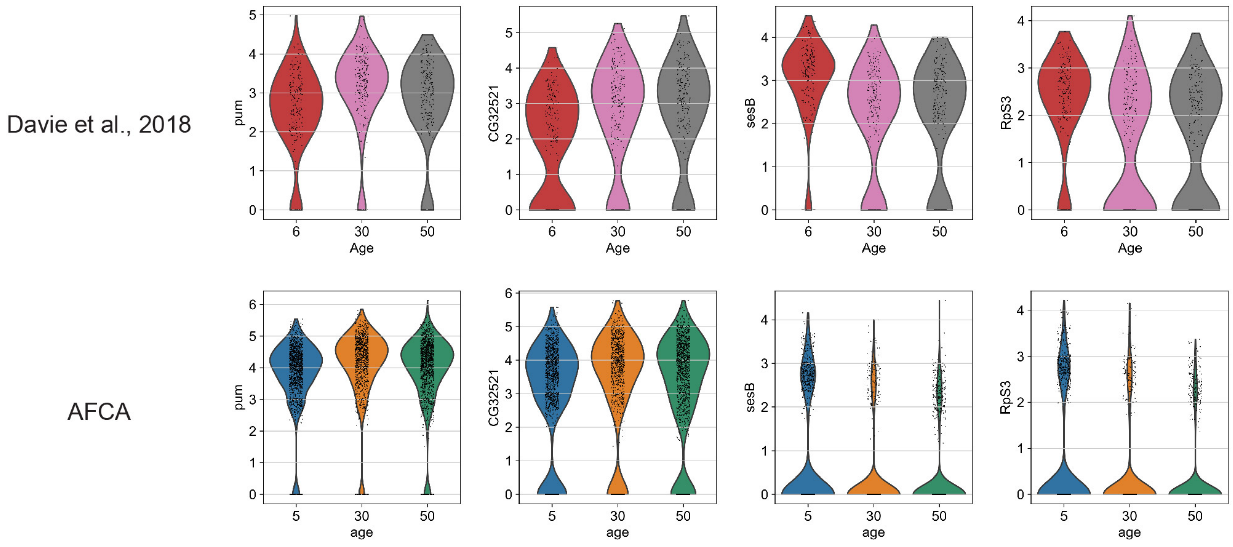


**C**

#### DEG ratio in Davie et al., 2018

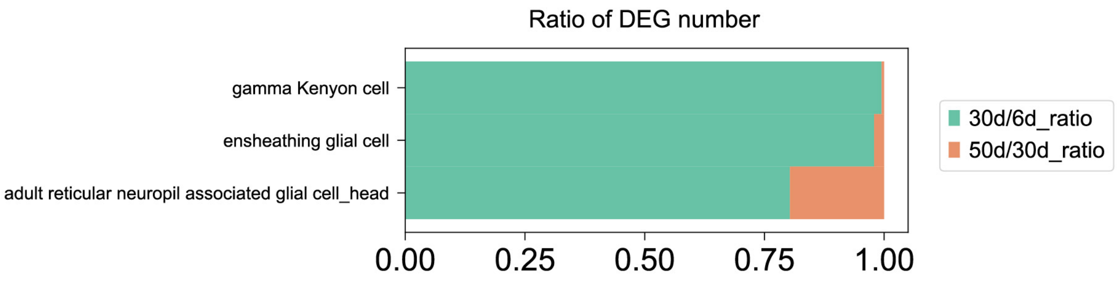

**Fig. S17. Comparisons between AFCA and Davie et al., 2018 data.**

A) Three cell types with the highest cell number from Davie et al., 2018 are compared to the corresponding cell types in the AFCA. X-axis indicates the log2 fold changes of gene expression between 30d and 5d samples in AFCA, while Y-axis represents the log2 changes between 30d and 6d samples in Davie et al., 2018. Orange dots are genes related to oxidative phosphorylation. Pearson correlations are performed to measure the correlation of log2 fold changes between two datasets. B) Aging DEGs in ensheathing glial cells are similarly regulated in AFCA and Davie et al. 2018 samples. C) Three cell types show a higher DEG ratio in the 30d/6d age interval compared to the 50d/30d one.

### Figure S18

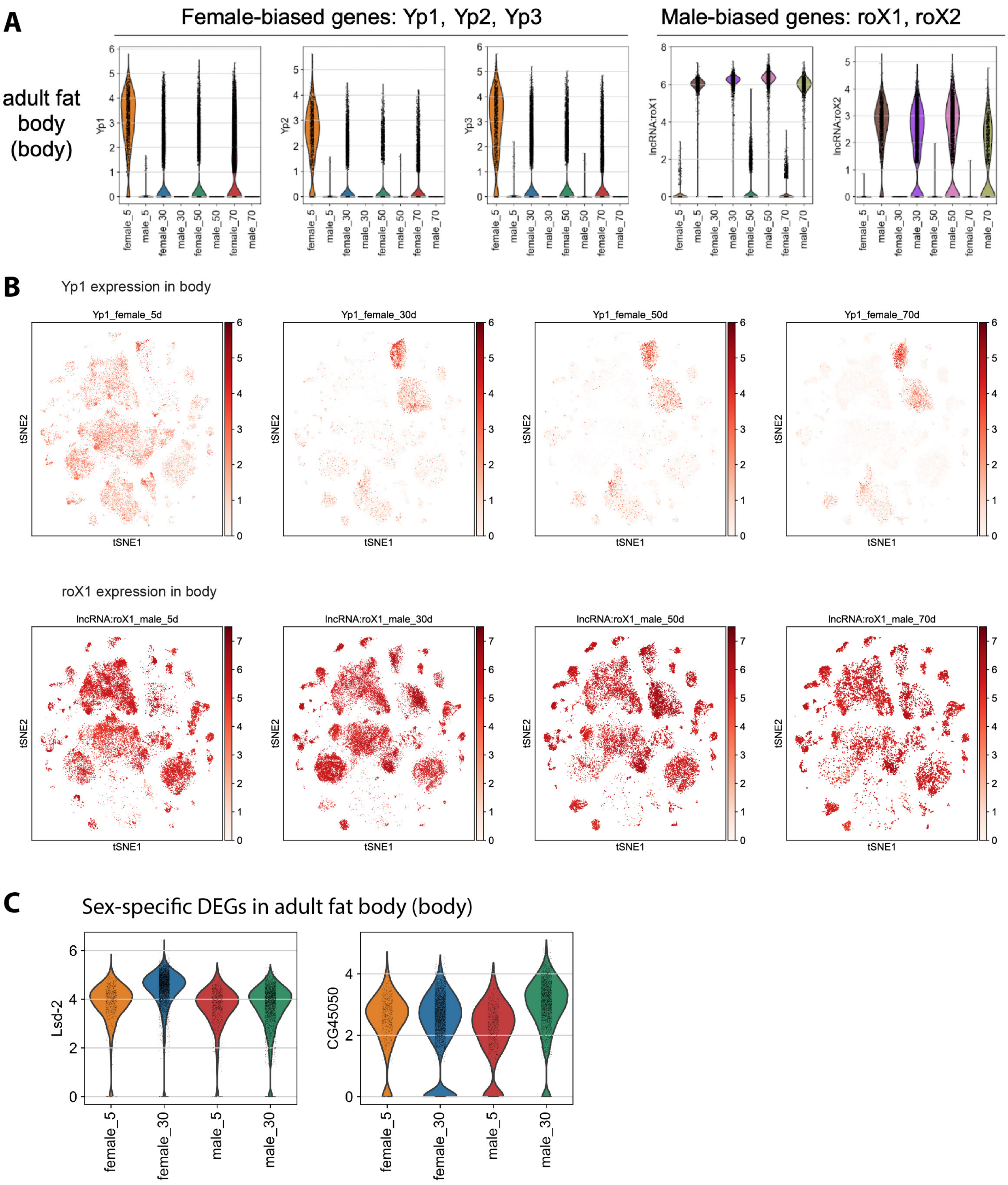
A
Female-biased genes: Yp1, Yp2, Yp3
Male-biased genes: roX1, roX2
adult fat body (body)
Yp1
Yp2
Yp3
lncRNA:roX1
lncRNA:roX2
female_5
male_5
female_30
male_30
female_50
male_50
female_70
male_70
B
Yp1 expression in body
Yp1_female_5d
Yp1_female_30d
Yp1_female_50d
Yp1_female_70d
tSNE1
tSNE2
roX1 expression in body
lncRNA:roX1_male_5d
lncRNA:roX1_male_30d
lncRNA:roX1_male_50d
lncRNA:roX1_male_70d
C
Sex-specific DEGs in adult fat body (body)
Lsd-2
CG45050

**Fig. S18 Sex-specific expression of genes.**

A) Expression of female- and male-biased genes from the fat body cells during aging. B) Expression of Yp1 and roX1 in body cells shown by tSNE. C) two example genes showing different change trends between males and females.

### Figure S19

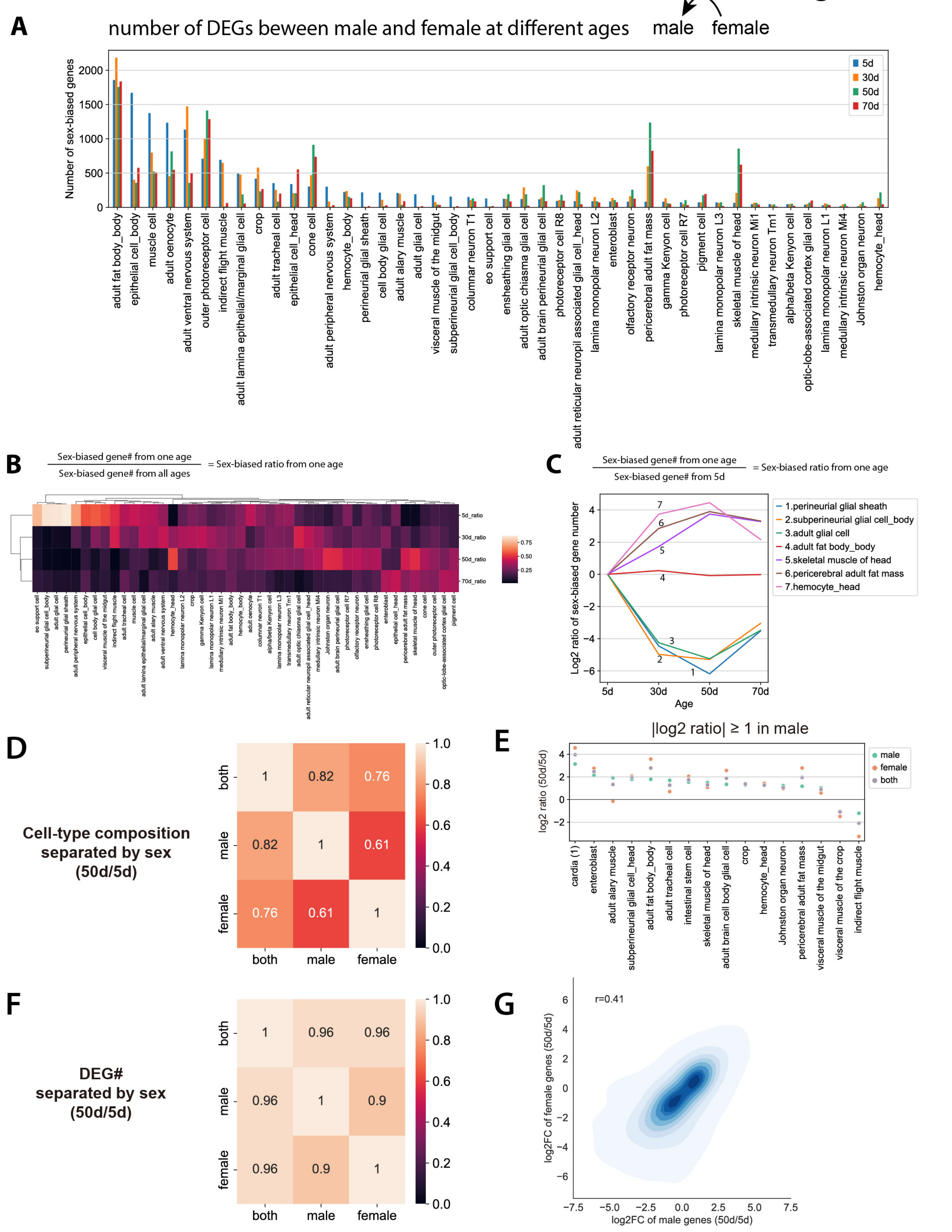

A
number of DEGs beween male and female at different ages
male
female
Number of sex-biased genes
5d
30d
50d
70d
B
Sex-biased gene# from one age
Sex-biased gene# from all ages
= Sex-biased ratio from one age
5d_ratio
30d_ratio
50d_ratio
70d_ratio
C
Sex-biased gene# from one age
Sex-biased gene# from 5d
= Sex-biased ratio from one age
Log2 ratio of sex-biased gene number
Age
1.perineurial glial sheath
2.subperineurial glial cell_body
3.adult glial cell
4.adult fat body_body
5.skeletal muscle of head
6.pericerebral adult fat mass
7.hemocyte_head
D
Cell-type composition separated by sex (50d/5d)
both
male
female
E
|log2 ratio| ≥ 1 in male
log2 ratio (50d/5d)
male
female
both
F
DEG# separated by sex (50d/5d)
G
r=0.41
log2FC of female genes (50d/5d)
log2FC of male genes (50d/5d)

**Fig. S19 Sex-biased genes from each age.**

A) The number of sex-specific genes from each age. B) Sex-biased ratios from different ages. C) Illustration of sex-biased ratios from several cell types. D) Spearman's rank correlation of the change of cell-type composition between males and females. (E) Log2 ratio of cell-type composition by comparing 50d to 5d samples in different sexes of flies. (F) Spearman's rank correlation of DEG number from each cell type by comparing 50d to 5d samples in male or female flies. (G) Comparison of log2 fold changes of gene expression between male and female. The Pearson correlation coefficient of male and female log2 fold changes is 0.41.

### Figure S20

**A**

Up-regulated

Down-regulated

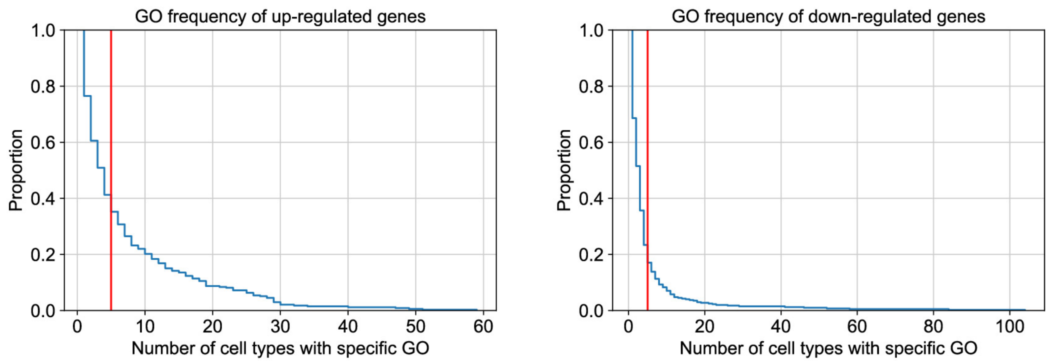


**B**

GOs with the lowest FDR using the up-regulated genes

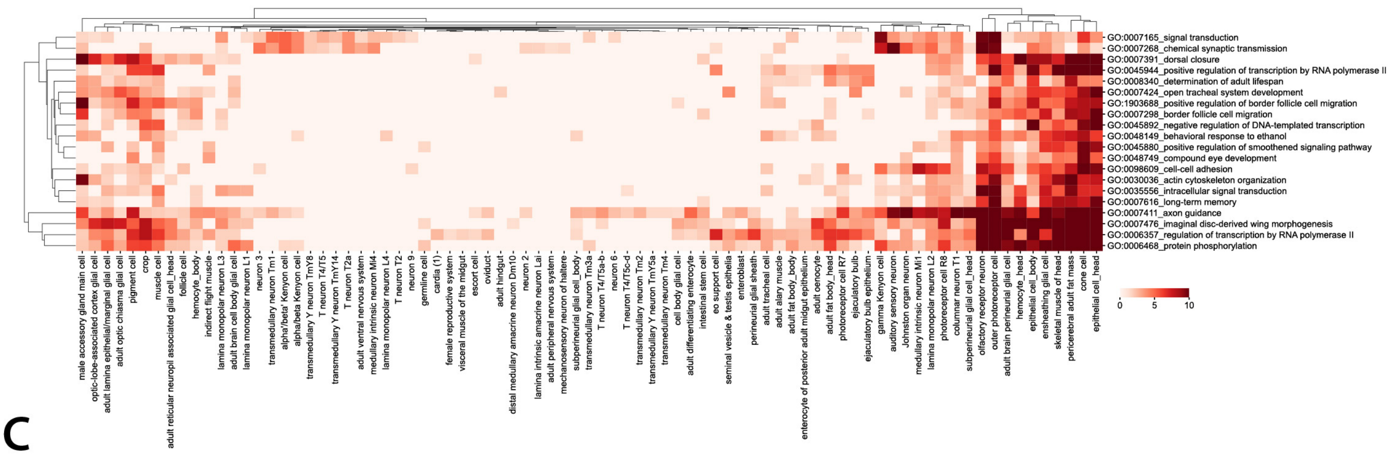


**C**

GOs with the lowest FDR using the down-regulated genes

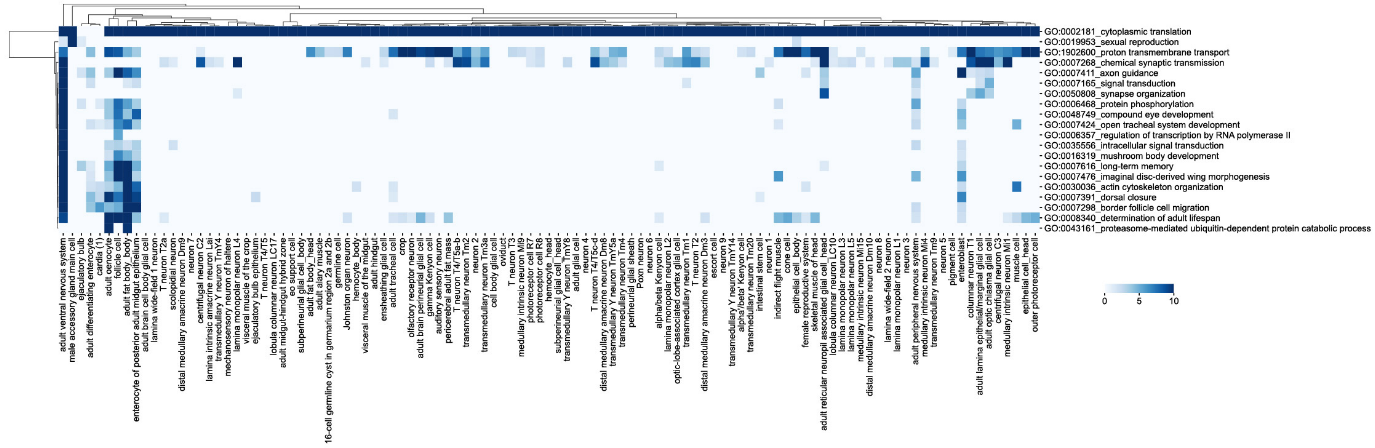

**Fig. S20 GO frequency and the most significant GOs from the up- or down-regulated genes.** A-B) Up- and down-regulated DEGs are identified by comparing 50d flies to 5d flies. A) GO frequencies from up- or down-regulated genes. GO frequency indicates how many cell types have the corresponding GO. The red line indicates five cell types with the corresponding GO. B) GOs with the lowest FDR from the up-regulated genes. C) GOs with the lowest FDR from the down-regulated genes.

Figure S21

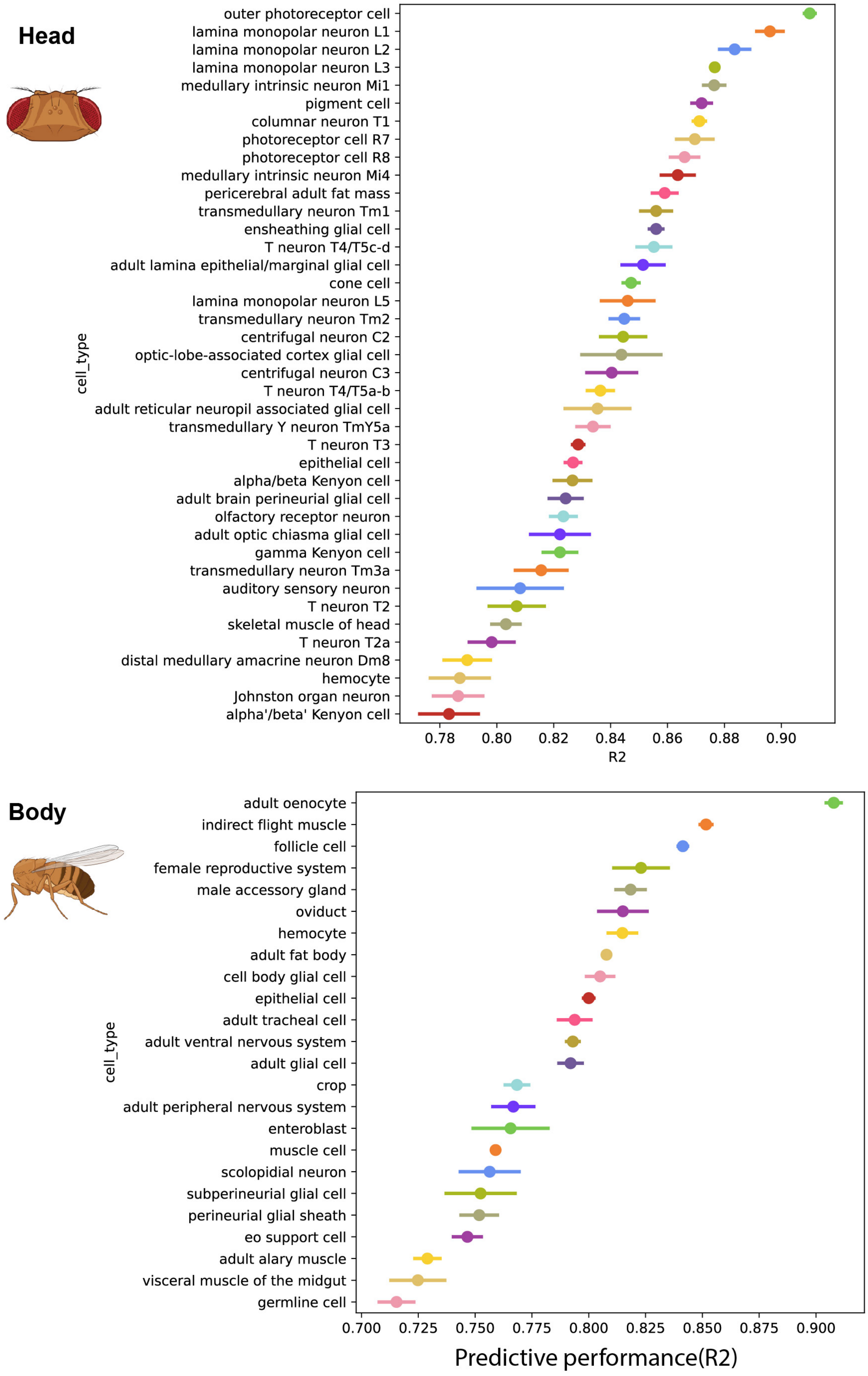

Head
outer photoreceptor cell
lamina monopolar neuron L1
lamina monopolar neuron L2
lamina monopolar neuron L3
medullary intrinsic neuron Mi1
pigment cell
columnar neuron T1
photoreceptor cell R7
photoreceptor cell R8
medullary intrinsic neuron Mi4
pericerebral adult fat mass
transmedullary neuron Tm1
ensheathing glial cell
T neuron T4/T5c-d
adult lamina epithelial/marginal glial cell
cone cell
lamina monopolar neuron L5
transmedullary neuron Tm2
centrifugal neuron C2
optic-lobe-associated cortex glial cell
centrifugal neuron C3
T neuron T4/T5a-b
adult reticular neuropil associated glial cell
transmedullary Y neuron TmY5a
T neuron T3
epithelial cell
alpha/beta Kenyon cell
adult brain perineurial glial cell
olfactory receptor neuron
adult optic chiasma glial cell
gamma Kenyon cell
transmedullary neuron Tm3a
auditory sensory neuron
T neuron T2
skeletal muscle of head
T neuron T2a
distal medullary amacrine neuron Dm8
hemocyte
Johnston organ neuron
alpha'/beta' Kenyon cell
cell_type
0.78
0.80
0.82
0.84
0.86
0.88
0.90
R2
Body
adult oenocyte
indirect flight muscle
follicle cell
female reproductive system
male accessory gland
oviduct
hemocyte
adult fat body
cell body glial cell
epithelial cell
adult tracheal cell
adult ventral nervous system
adult glial cell
crop
adult peripheral nervous system
enteroblast
muscle cell
scolopidial neuron
subperineurial glial cell
perineurial glial sheath
eo support cell
adult alary muscle
visceral muscle of the midgut
germline cell
cell_type
0.700
0.725
0.750
0.775
0.800
0.825
0.850
0.875
0.900
Predictive performance(R2)

**Fig. S21 Predictive performance of aging clocks.**
Predictive performances of aging clocks from each cell type. Performances of head and body cell types are shown separately.

### Figure S22

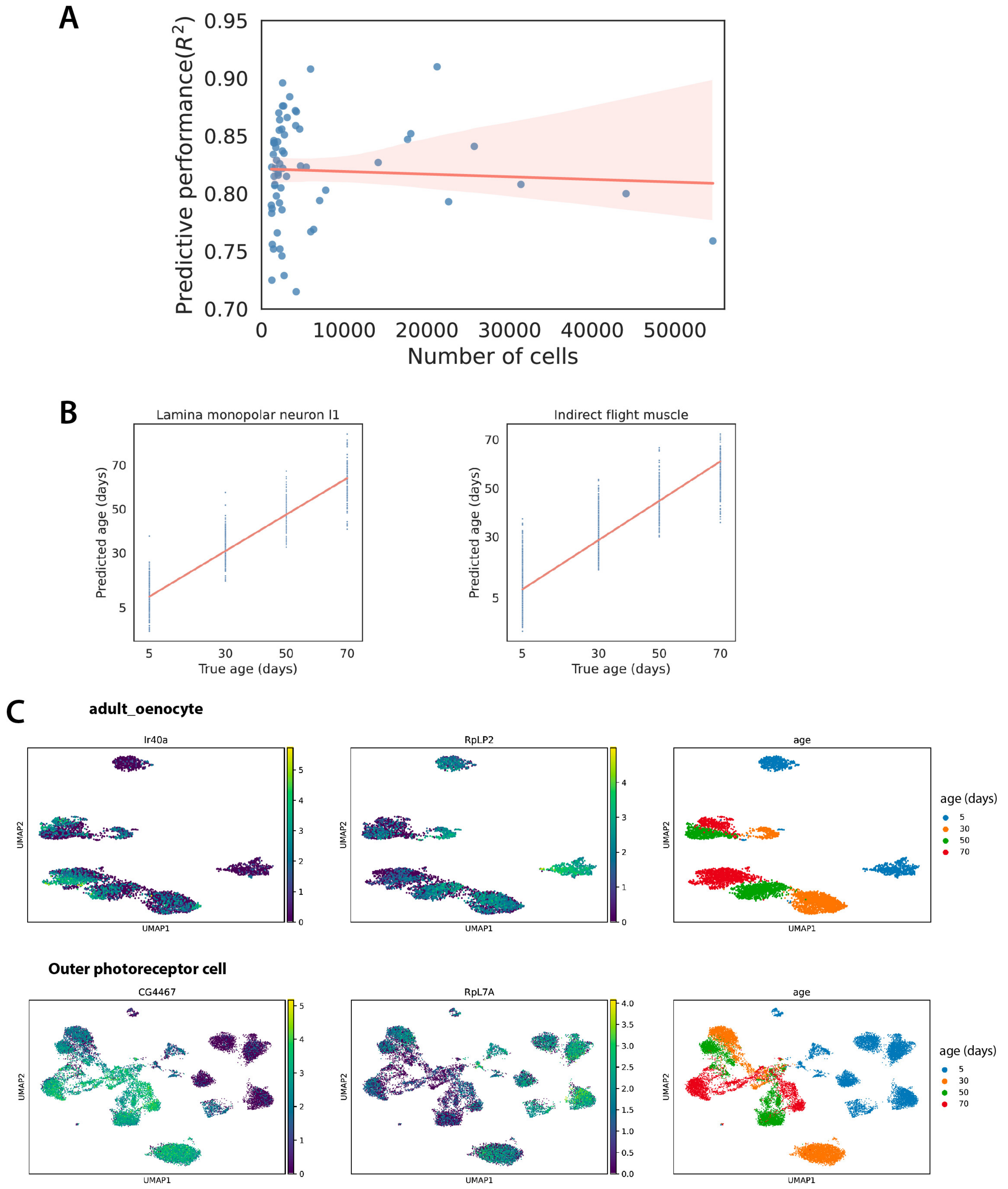
A
Predictive performance($R^2$)
0.95
0.90
0.85
0.80
0.75
0.70
0
10000
20000
30000
40000
50000
Number of cells
B
Lamina monopolar neuron l1
Predicted age (days)
True age (days)
5
30
50
70
Indirect flight muscle
Predicted age (days)
True age (days)
5
30
50
70
C
adult_oenocyte
Ir40a
RpLP2
age
UMAP1
UMAP2
age (days)
5
30
50
70
Outer photoreceptor cell
CG4467
RpL7A
age
UMAP1
UMAP2
age (days)
5
30
50
70

**Fig. S22 Aging clock profiles.**

A) The cell numbers from each cell type are not correlated to the predictive performance. B) Examples of two cell types with good prediction of true age. C) Expression of aging clock genes in two different cell types.

### Figure S23

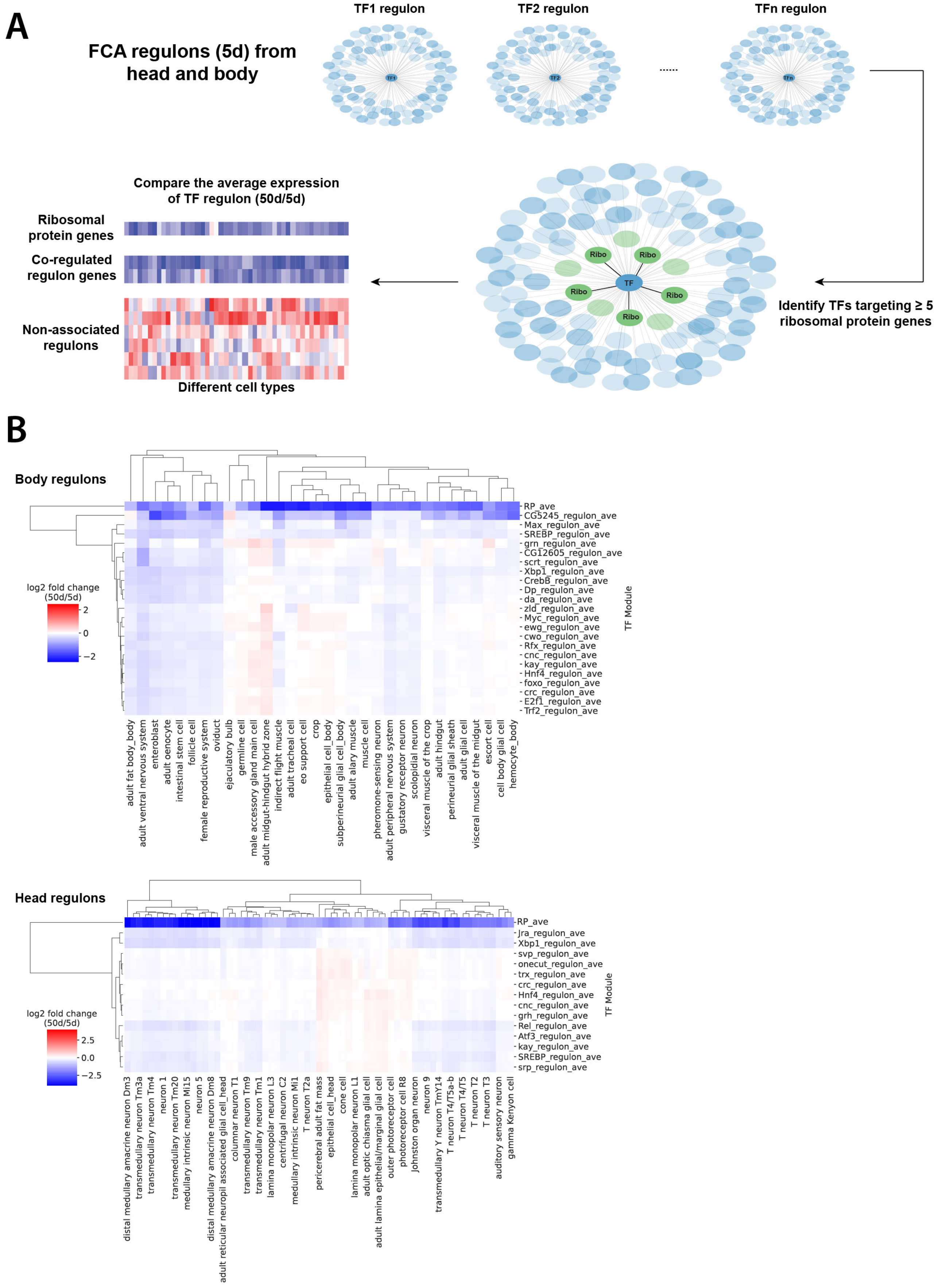
A
FCA regulons (5d) from head and body
TF1 regulon
TF2 regulon
TFn regulon
......
Identify TFs targeting ≥ 5 ribosomal protein genes
TF
Ribo
Compare the average expression of TF regulon (50d/5d)
Ribosomal protein genes
Co-regulated regulon genes
Non-associated regulons
Different cell types
B
Body regulons
log2 fold change (50d/5d)
2
0
−2
RP_ave
CG5245_regulon_ave
Max_regulon_ave
SREBP_regulon_ave
grn_regulon_ave
CG12605_regulon_ave
scrt_regulon_ave
Xbp1_regulon_ave
CrebB_regulon_ave
Dp_regulon_ave
da_regulon_ave
zld_regulon_ave
Myc_regulon_ave
ewg_regulon_ave
cwo_regulon_ave
Rfx_regulon_ave
cnc_regulon_ave
kay_regulon_ave
Hnf4_regulon_ave
foxo_regulon_ave
crc_regulon_ave
E2f1_regulon_ave
Trf2_regulon_ave
TF Module
adult fat body_body
adult ventral nervous system
enteroblast
adult oenocyte
intestinal stem cell
follicle cell
female reproductive system
oviduct
ejaculatory bulb
germline cell
male accessory gland main cell
adult midgut-hindgut hybrid zone
indirect flight muscle
adult tracheal cell
eo support cell
crop
epithelial cell_body
subperineurial glial cell_body
adult alary muscle
muscle cell
pheromone-sensing neuron
adult peripheral nervous system
gustatory receptor neuron
scolopidial neuron
visceral muscle of the crop
adult hindgut
perineurial glial sheath
adult glial cell
visceral muscle of the midgut
escort cell
cell body glial cell
hemocyte_body
Head regulons
log2 fold change (50d/5d)
2.5
0.0
−2.5
RP_ave
Jra_regulon_ave
Xbp1_regulon_ave
svp_regulon_ave
onecut_regulon_ave
trx_regulon_ave
crc_regulon_ave
Hnf4_regulon_ave
cnc_regulon_ave
grh_regulon_ave
Rel_regulon_ave
Atf3_regulon_ave
kay_regulon_ave
SREBP_regulon_ave
srp_regulon_ave
TF Module
distal medullary amacrine neuron Dm3
transmedullary neuron Tm3a
transmedullary neuron Tm4
neuron 1
transmedullary neuron Tm20
medullary intrinsic neuron Mi15
neuron 5
distal medullary amacrine neuron Dm8
adult reticular neuropil associated glial cell_head
columnar neuron T1
transmedullary neuron Tm9
transmedullary neuron Tm1
lamina monopolar neuron L3
centrifugal neuron C2
medullary intrinsic neuron Mi1
T neuron T2a
pericerebral adult fat mass
epithelial cell_head
cone cell
lamina monopolar neuron L1
adult optic chiasma glial cell
adult lamina epithelial/marginal glial cell
outer photoreceptor cell
photoreceptor cell R8
Johnston organ neuron
neuron 9
transmedullary Y neuron TmY14
T neuron T4/T5a-b
T neuron T4/T5
T neuron T2
T neuron T3
auditory sensory neuron
gamma Kenyon cell

**Fig. S23 Identification of possible TFs regulating the expression of ribosomal protein genes.**

A) The flowchart of identification of possible TFs of ribosomal protein genes. B) The averaged expressions of RP gene-related regulons.

### Figure S24

| fly_gene | mouse_ortholog | # cell types in fly | # cell types in mouse |
|---|---|---|---|
| RpL13A | Rpl13a | 8 | 24 |
| RpS25 | Rps25 | 13 | 18 |
| RpS29 | Rps29 | 5 | 24 |
| RpS26 | Rps26 | 10 | 10 |
| RpS21 | Rps21 | 3 | 15 |
| RpS15 | Rps15 | 15 | 2 |
| RpS23 | Rps23 | 6 | 11 |
| RpL27A | Rpl27a | 15 | 1 |
| RpS7 | Rps7 | 13 | 2 |
| RpL21 | Rpl21 | 11 | 2 |
| RpL35A | Rpl35a | 3 | 10 |
| RpL14 | Rpl14 | 11 | 1 |
| RpS12 | Rps12 | 7 | 5 |
| RpS8 | Rps8 | 11 | 1 |
| RpL23 | Rpl23 | 8 | 3 |
| RpL38 | Rpl38 | 1 | 10 |
| RpL10Ab | Rpl10a | 9 | 1 |
| RpS16 | Rps16 | 6 | 3 |
| RpS3 | Rps3 | 3 | 6 |
| sta | Rpsa | 7 | 1 |
| RpL13 | Rpl13 | 3 | 4 |
| RpL31 | Rpl31 | 4 | 3 |
| RpS17 | Rps17 | 2 | 5 |
| RpLP0 | Rplp0 | 4 | 2 |
| RpS20 | Rps20 | 5 | 1 |
| RpS24 | Rps24 | 4 | 1 |
| RpS30 | Fau | 3 | 2 |
| ATPsynbeta | Atp5b | 2 | 2 |
| RpL11 | Rpl11 | 3 | 1 |
| RpS11 | Rps11 | 3 | 1 |
| RpS6 | Rps6 | 3 | 1 |
| Hsc70-3 | Hspa5 | 1 | 2 |
| RpL24 | Rpl24 | 1 | 1 |

**Fig. S24 Aging clock genes present in both fly and mouse.**

1-1 orthologs of 33 aging clock genes conserved between fly and mouse. The number of cell types shown in each species indicates that the aging clock genes are detected as aging clock genes in the corresponding number of cell types.

### Figure S25

**Fly ribosomal protein gene** | **Mouse ribosomal protein gene**

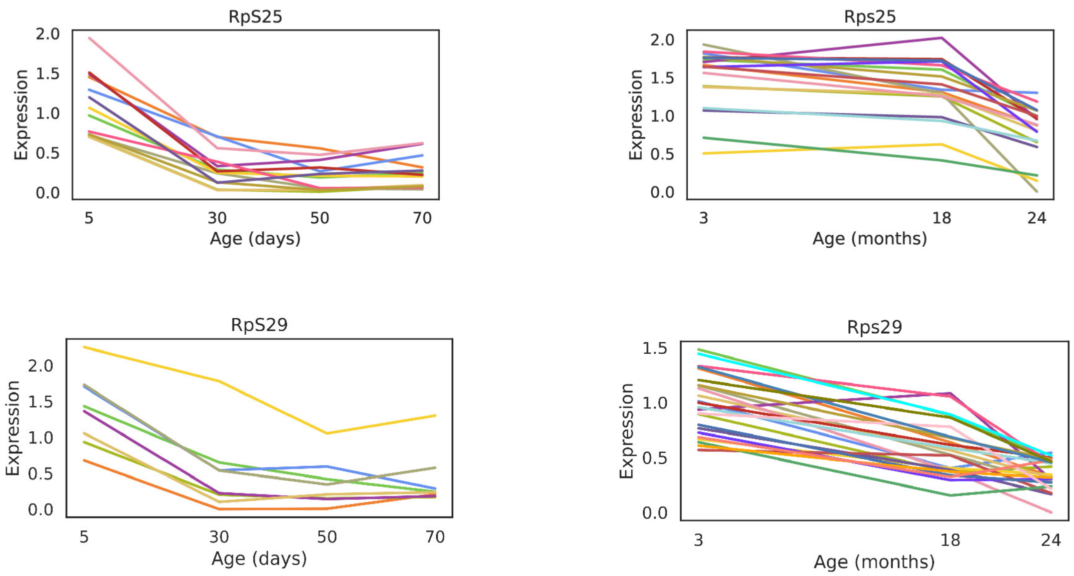

**Fig. S25 Examples of cross-species comparison of aging clock genes.**
Expression of two orthologous aging clock genes.

### Figure S26

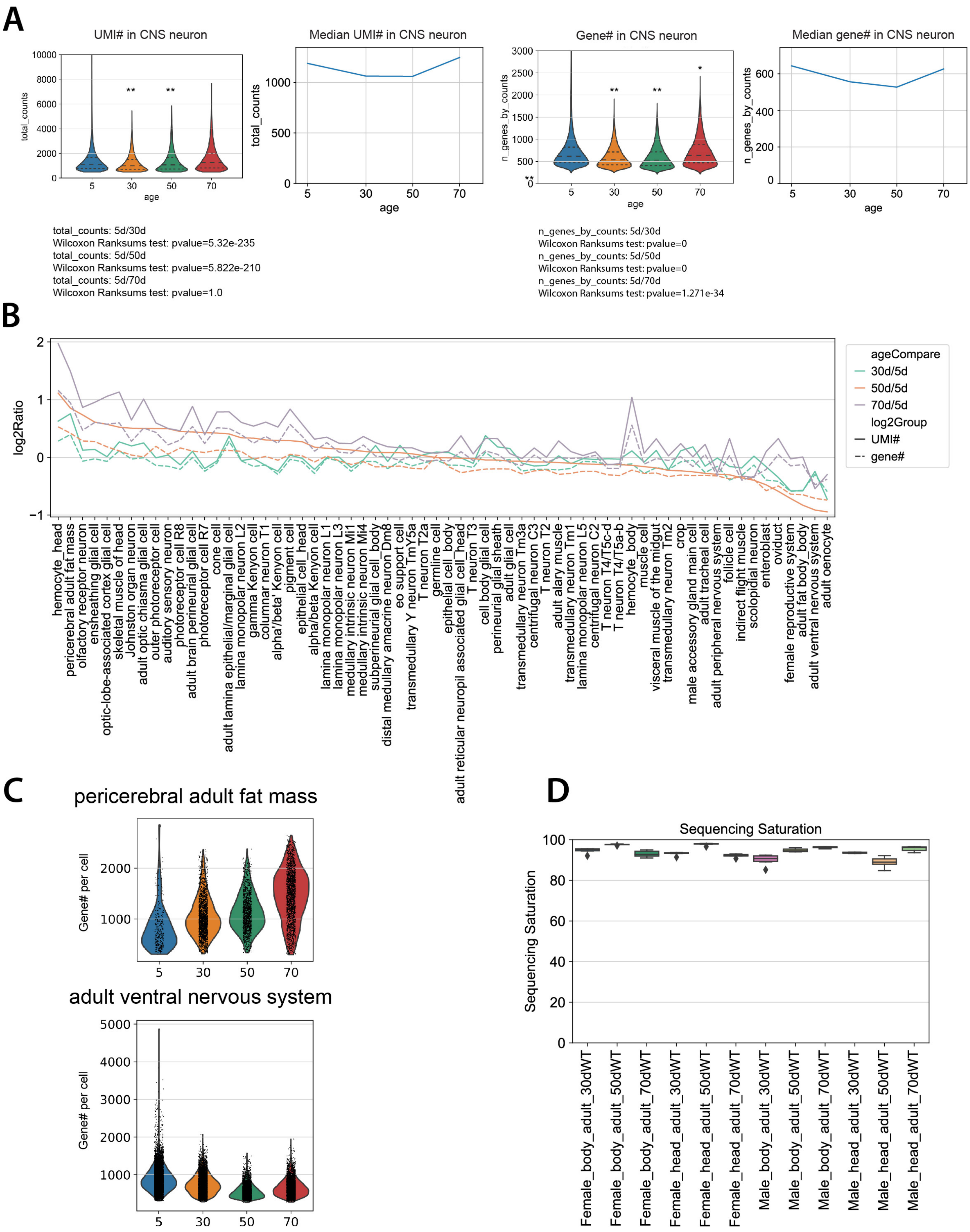
A
UMI# in CNS neuron
total_counts
age
**
**
Median UMI# in CNS neuron
total_counts
age
Gene# in CNS neuron
n_genes_by_counts
age
**
**
*
Median gene# in CNS neuron
n_genes_by_counts
age
total_counts: 5d/30d
Wilcoxon Ranksums test: pvalue=5.32e-235
total_counts: 5d/50d
Wilcoxon Ranksums test: pvalue=5.822e-210
total_counts: 5d/70d
Wilcoxon Ranksums test: pvalue=1.0
n_genes_by_counts: 5d/30d
Wilcoxon Ranksums test: pvalue=0
n_genes_by_counts: 5d/50d
Wilcoxon Ranksums test: pvalue=0
n_genes_by_counts: 5d/70d
Wilcoxon Ranksums test: pvalue=1.271e-34
B
log2Ratio
ageCompare
30d/5d
50d/5d
70d/5d
log2Group
UMI#
gene#
hemocyte_head
pericerebral adult fat mass
olfactory receptor neuron
ensheathing glial cell
optic-lobe-associated cortex glial cell
skeletal muscle of head
Johnston organ neuron
adult optic chiasma glial cell
outer photoreceptor cell
auditory sensory neuron
photoreceptor cell R8
adult brain perineurial glial cell
photoreceptor cell R7
cone cell
adult lamina epithelial/marginal glial cell
lamina monopolar neuron L2
gamma Kenyon cell
columnar neuron T1
alpha'/beta' Kenyon cell
pigment cell
epithelial cell_head
alpha/beta Kenyon cell
lamina monopolar neuron L1
lamina monopolar neuron L3
medullary intrinsic neuron Mi1
medullary intrinsic neuron Mi4
subperineurial glial cell_body
distal medullary amacrine neuron Dm8
eo support cell
transmedullary Y neuron TmY5a
T neuron T2a
germline cell
epithelial cell_body
adult reticular neuropil associated glial cell_head
T neuron T3
cell body glial cell
perineurial glial sheath
adult glial cell
transmedullary neuron Tm3a
centrifugal neuron C3
T neuron T2
adult alary muscle
transmedullary neuron Tm1
lamina monopolar neuron L5
centrifugal neuron C2
T neuron T4/T5c-d
T neuron T4/T5a-b
hemocyte_body
muscle cell
visceral muscle of the midgut
transmedullary neuron Tm2
crop
male accessory gland main cell
adult tracheal cell
adult peripheral nervous system
follicle cell
indirect flight muscle
scolopidial neuron
enteroblast
oviduct
female reproductive system
adult fat body_body
adult ventral nervous system
adult oenocyte
C
pericerebral adult fat mass
Gene# per cell
adult ventral nervous system
Gene# per cell
D
Sequencing Saturation
Sequencing Saturation
Female_body_adult_30dWT
Female_body_adult_50dWT
Female_body_adult_70dWT
Female_head_adult_30dWT
Female_head_adult_50dWT
Female_head_adult_70dWT
Male_body_adult_30dWT
Male_body_adult_50dWT
Male_body_adult_70dWT
Male_head_adult_30dWT
Male_head_adult_50dWT
Male_head_adult_70dWT

**Fig. S26 Variance of UMI and expressed gene numbers during aging.**

A) Numbers of expressed genes and UMIs per cell are decreased in the aged CNS neurons (30d and 50d). B) Log2 ratio of expressed gene or UMI number per cell during aging. C) Changes of expressed gene number in pericerebral adult fat mass and adult fat body from the body. D) Sequencing saturation of different batches of samples.

### Figure S27

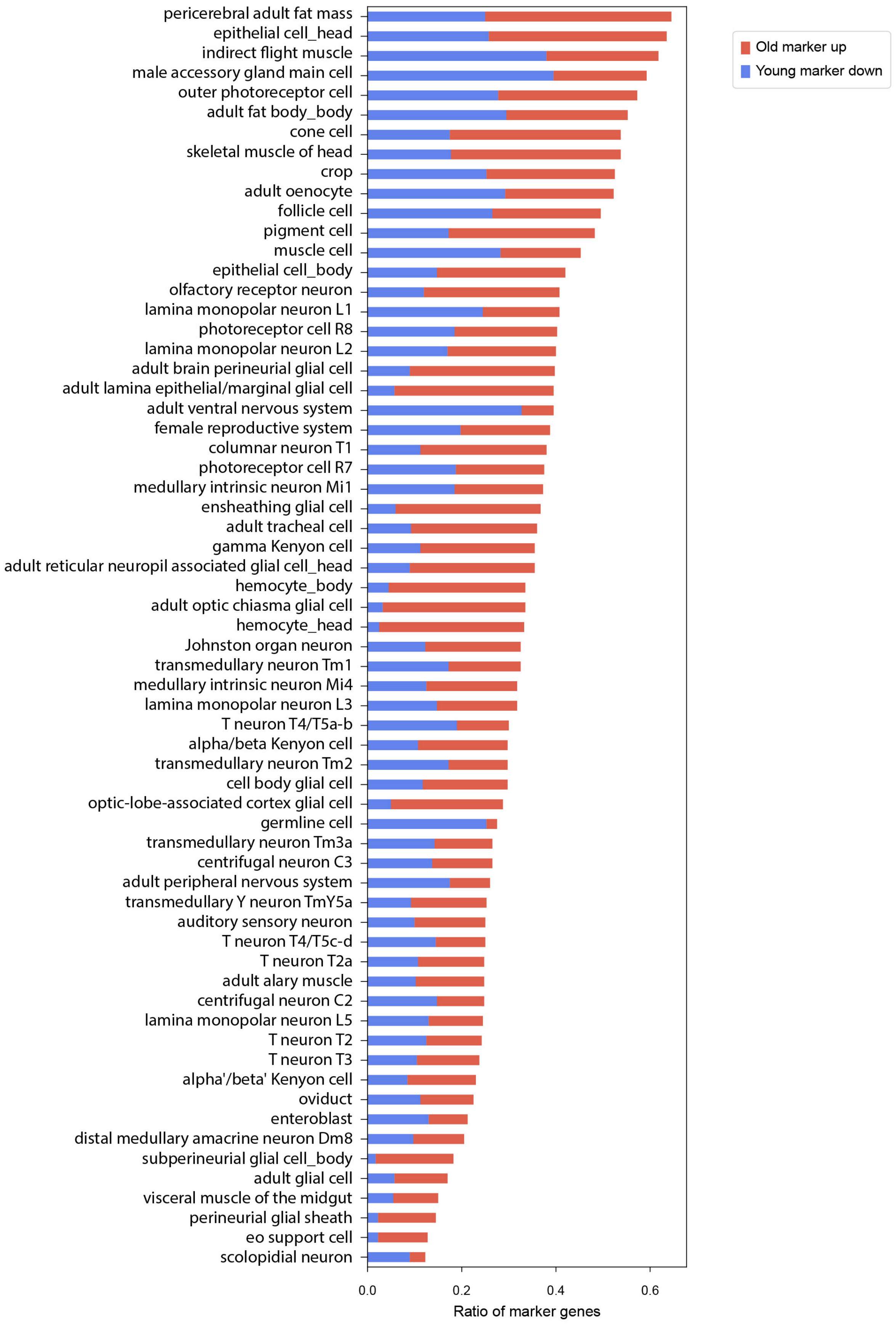

pericerebral adult fat mass
epithelial cell_head
indirect flight muscle
male accessory gland main cell
outer photoreceptor cell
adult fat body_body
cone cell
skeletal muscle of head
crop
adult oenocyte
follicle cell
pigment cell
muscle cell
epithelial cell_body
olfactory receptor neuron
lamina monopolar neuron L1
photoreceptor cell R8
lamina monopolar neuron L2
adult brain perineurial glial cell
adult lamina epithelial/marginal glial cell
adult ventral nervous system
female reproductive system
columnar neuron T1
photoreceptor cell R7
medullary intrinsic neuron Mi1
ensheathing glial cell
adult tracheal cell
gamma Kenyon cell
adult reticular neuropil associated glial cell_head
hemocyte_body
adult optic chiasma glial cell
hemocyte_head
Johnston organ neuron
transmedullary neuron Tm1
medullary intrinsic neuron Mi4
lamina monopolar neuron L3
T neuron T4/T5a-b
alpha/beta Kenyon cell
transmedullary neuron Tm2
cell body glial cell
optic-lobe-associated cortex glial cell
germline cell
transmedullary neuron Tm3a
centrifugal neuron C3
adult peripheral nervous system
transmedullary Y neuron TmY5a
auditory sensory neuron
T neuron T4/T5c-d
T neuron T2a
adult alary muscle
centrifugal neuron C2
lamina monopolar neuron L5
T neuron T2
T neuron T3
alpha'/beta' Kenyon cell
oviduct
enteroblast
distal medullary amacrine neuron Dm8
subperineurial glial cell_body
adult glial cell
visceral muscle of the midgut
perineurial glial sheath
eo support cell
scolopidial neuron
0.0
0.2
0.4
0.6
Ratio of marker genes
Old marker up
Young marker down

**Fig. S27 Changes of cell-type identity during aging.**

Cell types ranked by the decline of cell-type identity during aging. The decrease of young markers and the increase of old markers are shown separately in each cell type.

### Figure S28

Decreased expression of marker gene from the pericerebral adult fat mass

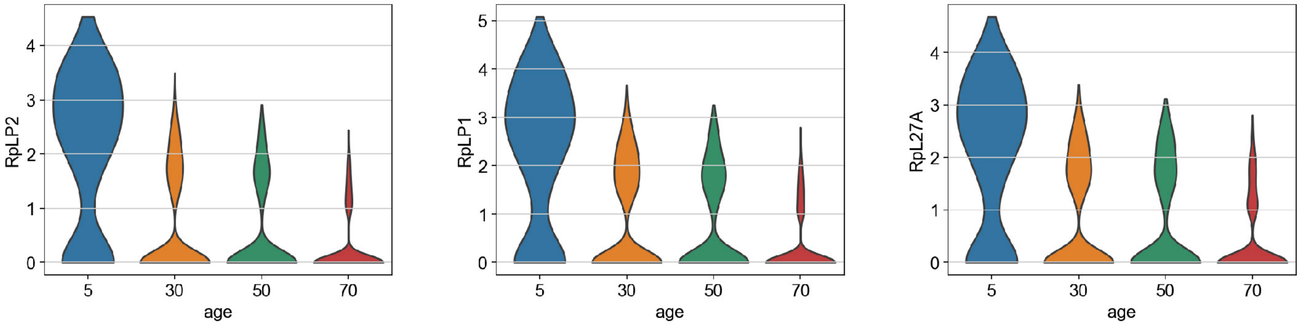


Increased expression of marker gene from the pericerebral adult fat mass

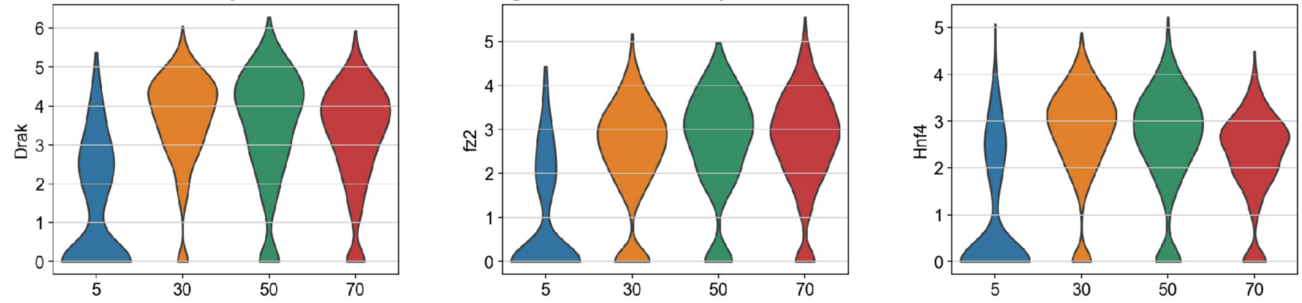


Decreased expression of marker gene from the male accessory gland main cell

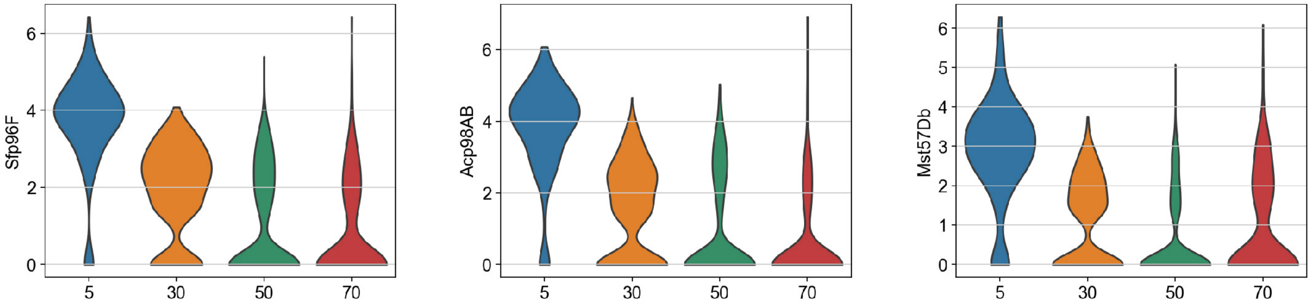


Increase expression of marker gene from the male accessory gland main cell

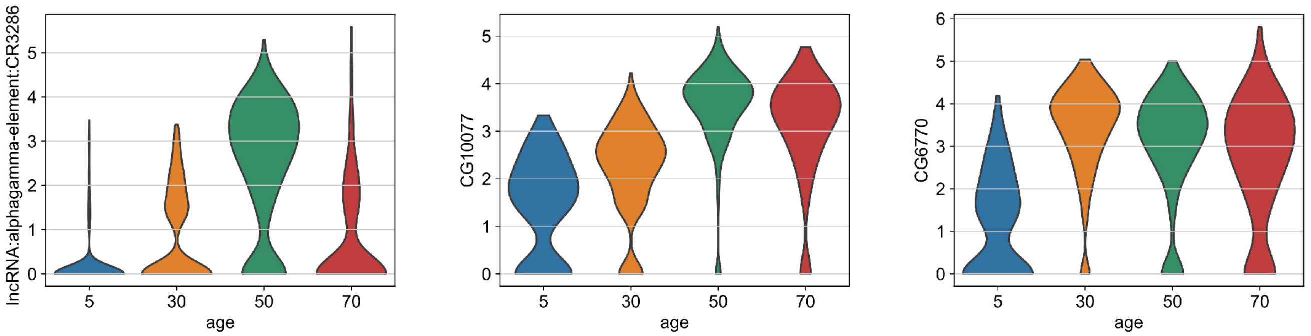

**Fig. S28 Marker genes that affect cell-type identities during aging.**
Examples of marker genes that caused the decline of cell-type identities.

### Figure S29

**A** Ranked by rank sum

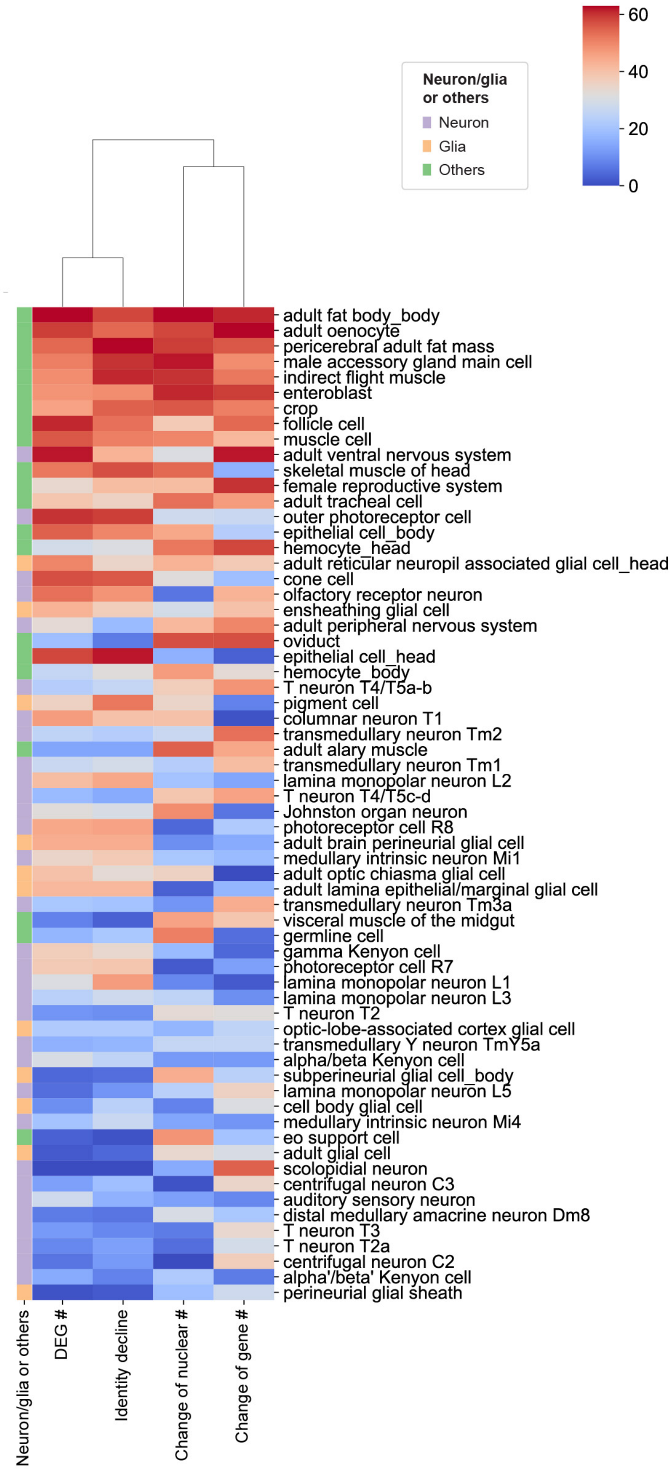


**B** Cluster different aging features

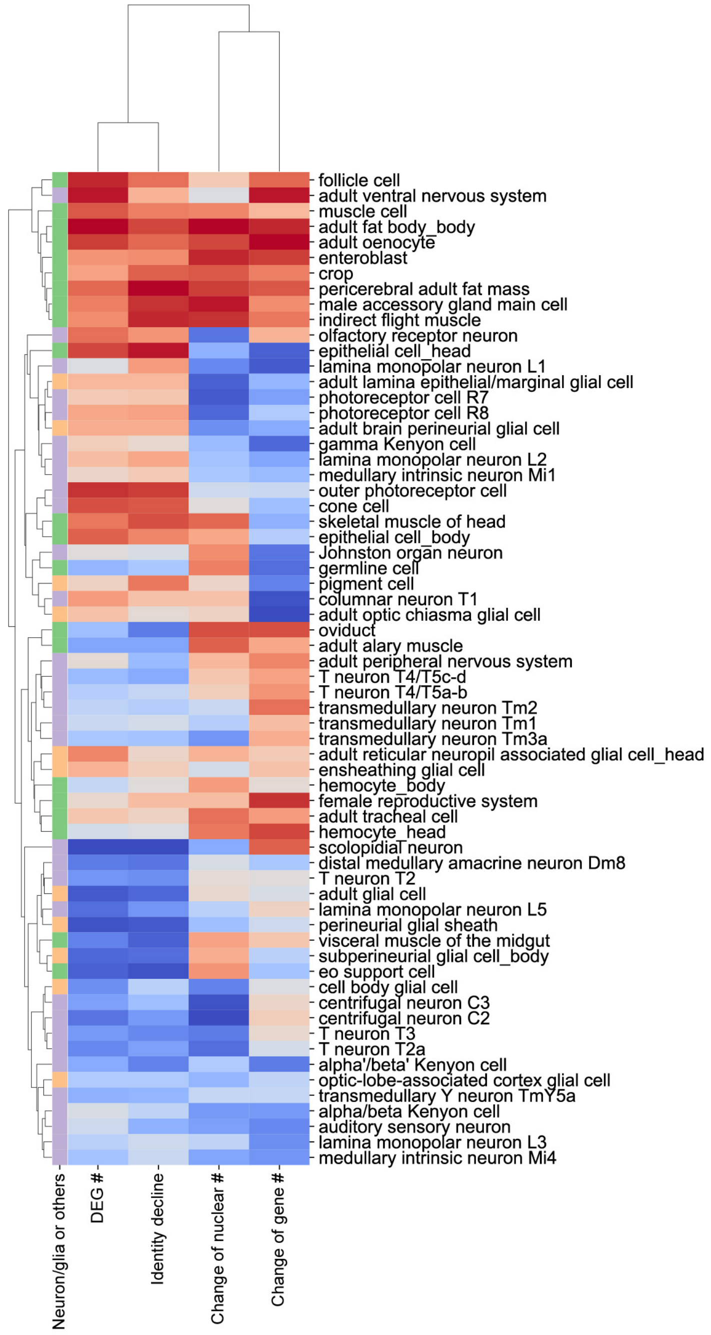

**Fig. S29 Rank sums of different aging features.**
A) Cell types ranked by the sum of different aging feature scores. B) Clustering of rank sum scores across different cell types.

### Figure S30

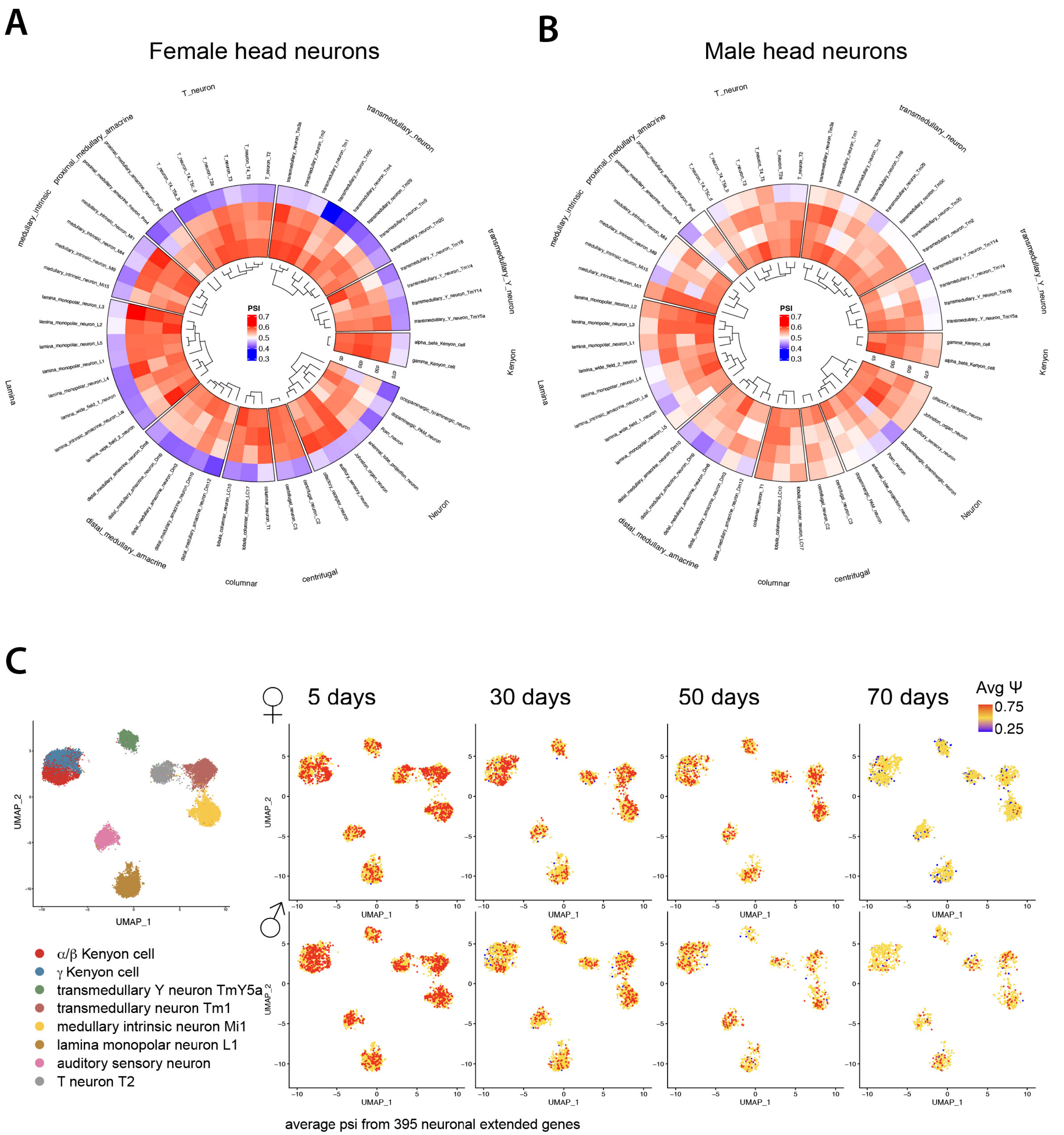
A
Female head neurons
B
Male head neurons
T_neuron
transmedullary_neuron
transmedullary_Y_neuron
Kenyon
Neuron
centrifugal
columnar
distal_medullary_amacrine
Lamina
medullary_intrinsic
proximal_medullary_amacrine
PSI
0.7
0.6
0.5
0.4
0.3
C
5 days
30 days
50 days
70 days
Avg Ψ
0.75
0.25
UMAP_1
UMAP_2
α/β Kenyon cell
γ Kenyon cell
transmedullary Y neuron TmY5a
transmedullary neuron Tm1
medullary intrinsic neuron Mi1
lamina monopolar neuron L1
auditory sensory neuron
T neuron T2
average psi from 395 neuronal extended genes

**Fig. S30 Comparison of alternative polyadenylation (APA) scores across different ages.** A-B) The circular heatmaps summarize APA trends from the annotated neuronal extended genes across different time points in females (A) and males (B). We use the LABRATsc package to calculate $\psi$ and the average value for 391 neuronal extended genes, which are plotted in each of the neuronal types from the AFCA head data. C) UMAP of different neuronal types designated by the AFCA. Average 3' isoform usage of 391 neural extended genes is plotted in individual cells by different aging time points. Both analyses show that neuronal extended 3' isoforms are progressively depleted during aging. This phenotype is stronger in females at 70d.

**Movies S1-S3. 3D reconstructions of two attached nuclei in the same fat body cell.**
The attached nuclei are stained by DAPI and LamC. The membranes of fat body cells are labeled by *cg-GAL4 > UAS-CD8GFP*.
